# Individualized brain-body signatures of positive emotions revealed by large-scale multimodal neurophysiological recording

**DOI:** 10.64898/2026.10.23.761434

**Authors:** Maryam Bijanzadeh, Bijurika Nandi, Vincent Wang, Bhavesh Kalisetti, Richard Campusano, Jennifer Townsend, Roger Anguera-Singla, Isabella D. Hartley, Melissa Osheroff, David A. Ziegler, Theodore P. Zanto, Adam Gazzaley, Reza Abbasi-Asl

## Abstract

Using one of the most comprehensive multimodal biosensing datasets in affective research, densely sampled within individuals, we asked how emotions are instantiated in each person’s brain and body. Four participants viewed 168 emotion-evoking videos during 128-channel EEG and 28 autonomic, ocular and facial recordings. Cross-validated elastic-net classifiers distinguished each of 15 positive and neutral emotions from baseline in every participant (mean accuracy 87-97%). We found that no emotion was predicted by a single marker and each reflected a multimodal ensemble of cortical and autonomic activity whose composition was largely person-specific. Yet physiology-derived groupings converged selectively with self-report; Interest-Awe-Amusement-Satisfaction and Entrancement-Aesthetic Appreciation recurred across individuals based on physiology. Across emotion groups, frontal and posterior delta, posterior alpha and electrodermal activity formed a shared backbone, plausibly reflecting affective and sensory engagement, whereas cardiovascular and facial signals varied between people. This framework yields individualized, objective neurophysiological indicators of emotion, potentially aiding personalized diagnosis.

## Introduction

Emotions shape cognition, decision-making and social behavior, yet remain among the most difficult mental states to measure objectively. An emotional episode is a coordinated brain-body event: cortical oscillations shift, cardiovascular, electrodermal, respiratory and gastric activity change, and facial muscles engage^1,2^. These responses arise from bidirectional interactions between cortical and subcortical circuits and the autonomic nervous system (ANS), with interoceptive hubs such as the insula and anterior cingulate cortex both reading out and regulating visceral state^3–5^. When any or all parts of these systems are dysregulated, mood and anxiety disorders can emerge, thus identifying how brain and body jointly encode emotional experience is crucial for developing biomarkers to monitor and treat them^6,7^.

Emotional experience is also highly individual. The cognitive, semantic, and bodily expressions, accompanying awe, amusement or boredom differ between people and, within a person, with physiological, circadian and mood state^8,9^. This variability is not merely noise: designs that average a few trials across many participants recover shared features but discard the within-person structure of an individual’s emotional repertoire, such that population-derived biomarkers often generalize poorly to individuals^10,11^. Dense and longitudinal sampling of many trials per emotion within the same person, combined with interpretable, rigorous machine learning approaches, instead allows probing personalized response profiles central to precision neuroscience^12–14^, shifting the question from “what is common to emotion across people” to “how a given emotion is instantiated in a particular person’s brain and body”.

Most neurophysiological studies of emotion examine central or peripheral signals in isolation. Autonomic measures such as heart-rate variability, electrodermal activity, blood pressure and respiration chiefly index arousal^1,15^, although their multivariate combinations carry partial information about emotion categories^16,17^. fMRI patterns decode discrete categories^18,19^, and video-evoked emotion is represented in a high-dimensional, distributed neural space with smooth gradients between categories^20–22^. Scalp and intracranial EEG associate spectral features to emotion categories, for example theta-band power during amusement, interest and joy^23^ and gamma-band activity during affective behavior^24^. Yet large comprehensive multimodal datasets typically sample many people briefly rather than each person densely^25,26^.

Here we recorded 128-channel EEG together with 28 peripheral sensors, including electrocardiography, electrodermal activity, facial electromyography (zygomaticus and corrugator), respiration, electrogastrography (EGG), continuous blood pressure, ocular signals and trans-radial electrical bioimpedance velocimetry (TREV), while participants viewed 168 emotionally evocative 1-minute videos^20^ and reported the emotions they experienced. Using interpretable, regularized classifiers with block-wise cross-validation, we asked three questions. First, can each of 15 emotions be distinguished from a no-stimulus baseline within an individual? Second, which neurophysiological features drive these distinctions, and which are shared versus emotion-specific? Third, do emotions group together by their brain-body physiology, and does this grouping agree with self-report^20,21^? The framework recovered individualized neurophysiological signatures of emotion whose organization is largely person-specific, partly shared across individuals, and consistent with the multidimensional structure of reported experience.

## Results

To investigate individualized markers of rated emotions, we collected one of the most comprehensive multimodal biosensing datasets to date: four participants, each completing four 1-hour sessions (168 one-minute trials per participant). In each trial, participants watched a ∼1-minute block of emotionally evocative video clips with similar emotional content, previously validated by crowd workers^20^ (see Video Stimulation in Methods). They then rated their experience on 18 emotion categories (0–5 scale), of which 15 were consistently selected. Finally, they fixated on a central cross for a 30-second no-stimulus baseline (Fig. 1A). One-minute trials accommodate slower autonomic responses such as EDA^27^, and dense repeated sampling provides the within-person statistical power required to estimate personalized emotion signatures.

**Figure 1.**
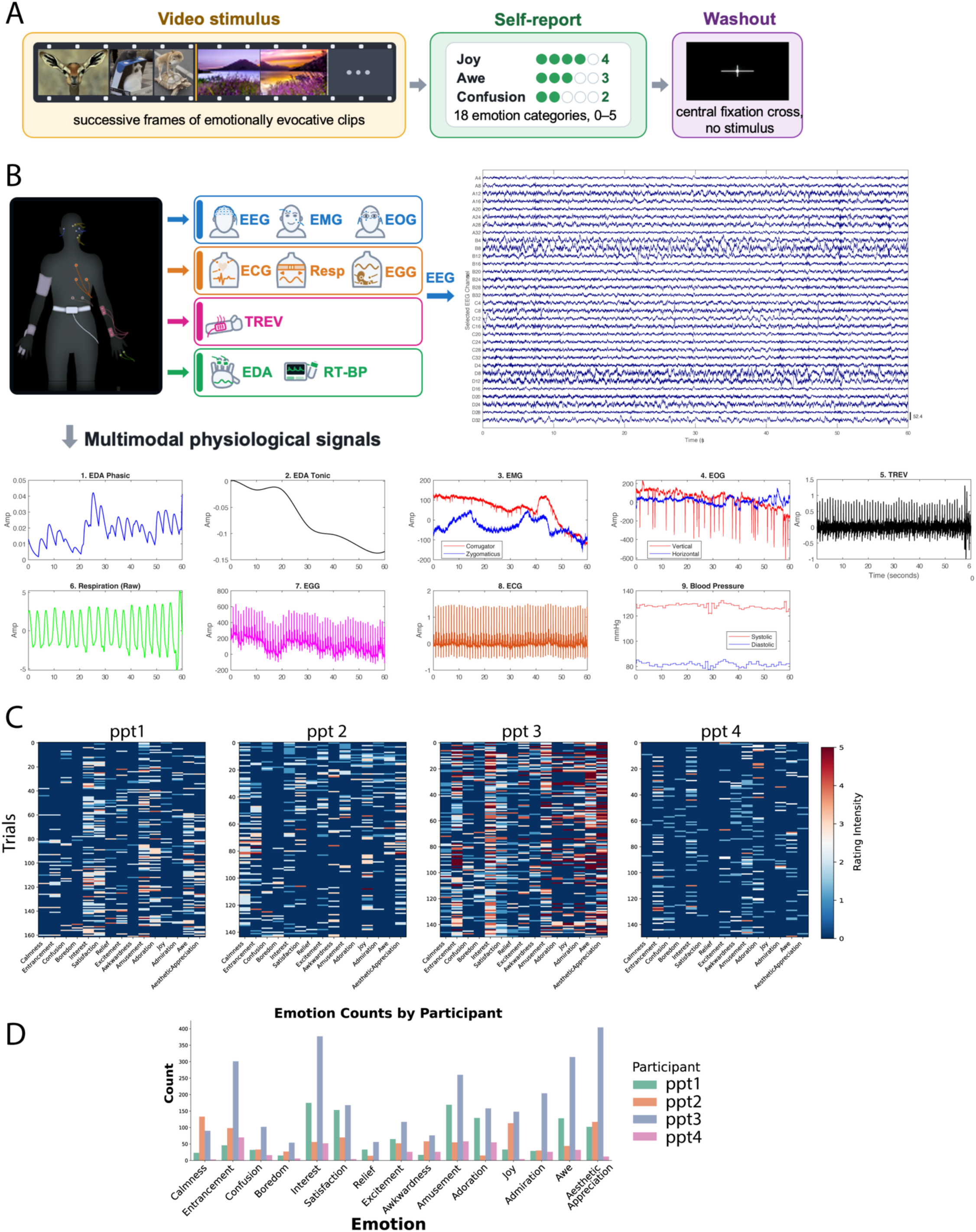
Experimental paradigm, multimodal biosensing setup and self-report structure. **A. Trial structure.** In each trial, participants viewed a ∼1-min block of emotionally evocative video clips (yellow) from a validated normative library, assembled from mutually exclusive clips so that each block carried coherent emotional content. They then rated the emotion(s) they experienced on 18 positive and neutral categories (0–5 scale; green) and fixated a central cross during a 30-s no-stimulus baseline (purple). Four participants each completed four 1-h sessions, yielding 168 one-minute trials per participant. **B. Multimodal biosensing (MMBS) acquisition.** Left, sensor montage: 128-channel BioSemi ActiveTwo EEG with 28 peripheral readouts (BIOPAC and CNAP RT-BP): facial EMG (zygomaticus, corrugator), vertical and horizontal EOG, ECG, respiration, electrogastrography (EGG), electrodermal activity (EDA), trans-radial electrical bioimpedance velocimetry (TREV; a proxy for radial blood flow and relative cardiac contractility) and continuous systolic/diastolic blood pressure. Right, 60 s of preprocessed EEG from a representative subset of channels (0.5–55 Hz, ocular components removed by ICA, common-average referenced; scale bar 52.4 μV). Bottom, simultaneously recorded peripheral signals over the same 60-s window: (1) phasic and (2) tonic EDA (cvxEDA decomposition), (3) corrugator and zygomaticus EMG, (4) vertical and horizontal EOG, (5) TREV, (6) raw respiration, (7) EGG, (8) ECG, and (9) beat-to-beat systolic and diastolic blood pressure. **C. Trial-by-trial self-report ratings** for each participant (ppt 1–4). Rows are trials and columns are the 15 emotion categories retained for analysis (Nostalgia, Romance and Craving were excluded; Methods); color encodes rating intensity (0–5). **D. Emotion endorsement frequency** by participant. Total counts ranged from 390 (ppt 4) to 2,829 (ppt 3), and the rank order of emotions differed across participants (Kendall’s W = 0.504), motivating the within-individual modeling framework used throughout.

We recorded 128-channel EEG and 28 peripheral sensors (EMG, ECG, EDA, EOG, TREV, EGG, respiration and blood pressure; Fig. 1B). After preprocessing (Methods; Extended Data Fig. 1), multitaper power spectra were averaged in six bands (delta 1-4 Hz, theta 4-8 Hz, low alpha 8-10 Hz, alpha 10-12 Hz, beta 12-30 Hz, gamma 30-55 Hz). Band power was then averaged within nine regions of a 3 × 3 topographic grid (posterior, middle, frontal × left, medial, right; Supplementary Fig. 1). We derived peripheral features such as heart rate variability, tonic and phasic EDA, facial EMG activity and cardiac contractility from TREV (Methods). This yielded 102 features per 10-second window: 54 spectral and 48 peripheral (Supplementary Table T1).

Self-reports varied substantially both between and within participants (Fig. 1C, D), consistent with previous reports^20^. Total emotion counts per participant ranged from 390 (Ppt 4) to 2,829 (Ppt 3) (mean = 1,320.75, SD = 1,054). Rank standard deviations of emotion counts across participants ranged from 6.05 (Calmness) to 1 (Confusion) (Extended Data Table 1), indicating moderate consensus (Kendall’s W = 0.504).

### Emotionally-weighted stimuli are reliably classified

We first trained balanced binary classifiers to test whether emotionally evocative stimuli are distinguishable from the no-stimulus baseline using each participant’s multimodal neurophysiological signals (stimulus-driven classifiers). To prevent slow drift and session differences from masquerading as task effects, each feature was z-scored within its recording block, jointly across task and baseline segments (Fig. 2A). Evaluation used block-wise GroupKFold, holding out entire blocks (six sequential trials each) so that temporally adjacent trials never appeared in both training and test sets, guarding against data leakage and overfitting. Majority-class segments were subsampled to balance classes within each fold (Fig. 2A, top right), and hyperparameters were tuned by grid search on a nested validation set (Methods, Extended Data Figure 1). Stimulus-driven classifiers separated emotionally rated epochs from baseline in every participant. Area under the receiver-operating-characteristic curve (AUC) and accuracy were well above chance (50%) for logistic regression with elastic net (LR-EN), linear support vector machine (SVM) and random forest (RF; Fig. 2B, inset); Stimulus-driven mean AUCs were 0.95–1.00. Comparable performance across model families indicates robust discriminative information rather than an algorithmic artifact, although RF performed slightly worse in most participants than LR-EN and SVM (Fig. 2B, inset). We proceeded with LR-EN because its feature weights are interpretable and directional, and it costs less to compute than SVM.

**Figure 2.**
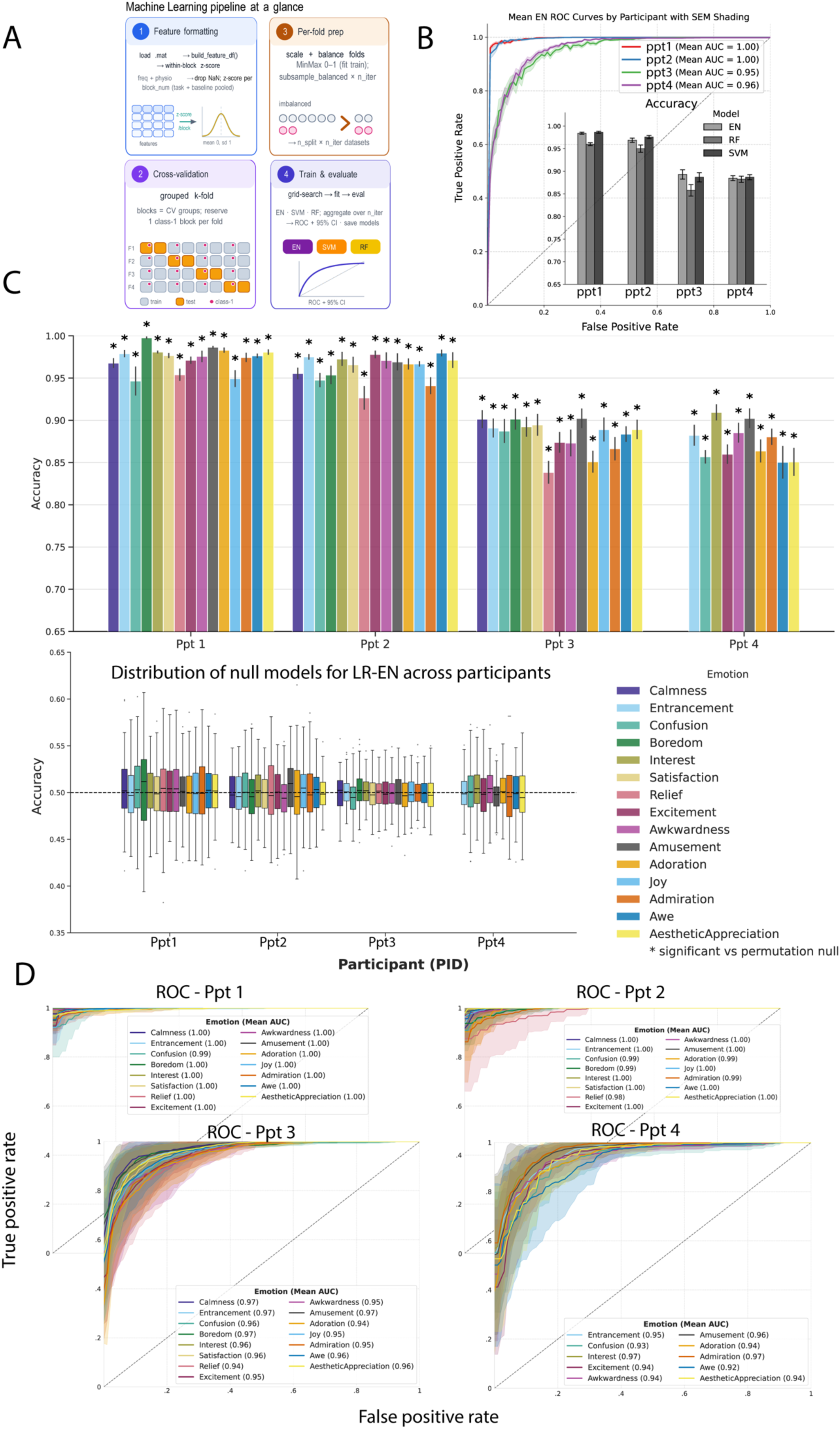
Within-individual decoding of emotional states from multimodal neurophysiology. **A. Analysis pipeline.** (1) Feature formatting: 54 EEG spectral features (6 canonical bands × 9 regional electrode groups, oscillatory power isolated with multitaper spectra and FOOOF) and 48 peripheral physiological features were extracted from 10-s segments with 50% overlap; each feature was z-scored within its recording block, jointly across task and baseline segments, and segments with missing values were dropped. (2) Cross-validation: recording blocks (6 sequential trials each) served as groups in a custom block-grouped k-fold scheme that guaranteed both classes in every fold, so that temporally adjacent trials never appeared in both training and test sets. (3) Per-fold preparation: min–max scaling was fitted on training data only, and the majority class was subsampled five times without replacement, separately in training and test partitions. (4) Training and evaluation: elastic-net logistic regression (EN), linear support vector machine (SVM) and random forest (RF) were fit with per-fold grid-search hyperparameter tuning (internal 2-fold CV optimizing AUC), and performance was aggregated across folds. **B. Stimulus-versus-baseline classification.** Mean ROC curves for elastic-net classifiers separating all stimulus trials (emotion ratings pooled) from no-stimulus baseline segments, shown per participant; shading denotes s.e.m. across cross-validation folds and values in parentheses give mean AUC. Dashed diagonal marks chance. Inset, classification accuracy for the three model families (EN, RF, SVM) in each participant; error bars denote s.e.m. across folds. Performance was well above chance (0.5) for all model families, with RF slightly below EN and SVM in most participants; **C. Per-emotion classification.** Top, accuracy of separate EN classifiers distinguishing each emotion from the no-stimulus baseline, for each participant; bars are colored by emotion, and error bars denote s.e.m. across cross-validation runs (n = 25). Mean accuracy across emotions ranged from 0.87 (ppt 4) to 0.97 (ppt 1). Bottom, accuracy of label-permuted surrogate models trained identically (box plots; dashed line, chance), which centered on chance (mean 0.49–0.50); observed accuracies fell outside the surrogate confidence intervals for every emotion (Supplementary Table T2). **D.** ROC curves for each emotion versus baseline, per participant; shading denotes cross-validation variability and values in parentheses give mean AUC per emotion. Ppt 4 contributes 10 emotions because five were rated in too few trials (Methods).

To test whether each of the 15 positive and neutral emotions was distinguishable from baseline, we trained a separate LR-EN classifier per emotion within the same framework. Each contrasted positively rated trials (class 1) with baseline segments (class 0; Methods). Every emotion was separable from baseline (Fig. 2C, D). Performance approached ceiling for several participants and emotions. It also significantly exceeded label-shuffled surrogate models trained identically, falling outside their confidence intervals (Fig. 2C, bottom; Supplementary Table T2). Thus, within an individual, distinct emotions carry distinguishable multimodal signatures relative to the no-stimulus phase.

We additionally trained one-vs-all classifiers, contrasting trials containing the target emotion with all others (Supplementary Fig. S2). One-vs-all classifiers performed lower and more variably, thus we used emotion-vs-baseline models to probe neurophysiological features. The weaker one-vs-all performance likely reflects the task design: participants could rate multiple emotions per trial, entangling their neurophysiological underpinnings.

### Interpretable models reveal per-emotion feature signatures for each participant

We next asked whether neurophysiological features are selective for specific emotions or shared across several. Each LR-EN classifier assigns every feature a weight reflecting its contribution in recognizing a given emotion. Retaining only stable features (see Methods; Fig. 3A), the resulting profiles were rich and emotion-specific, typically spanning cortical (EEG band-power) and peripheral (e.g., cardiovascular, electrodermal) features (Supplementary Figs. S3–S6). Figure 3 illustrates this architecture for a representative participant, using Joy as an example (Fig. 3A, B).

**Figure 3.**
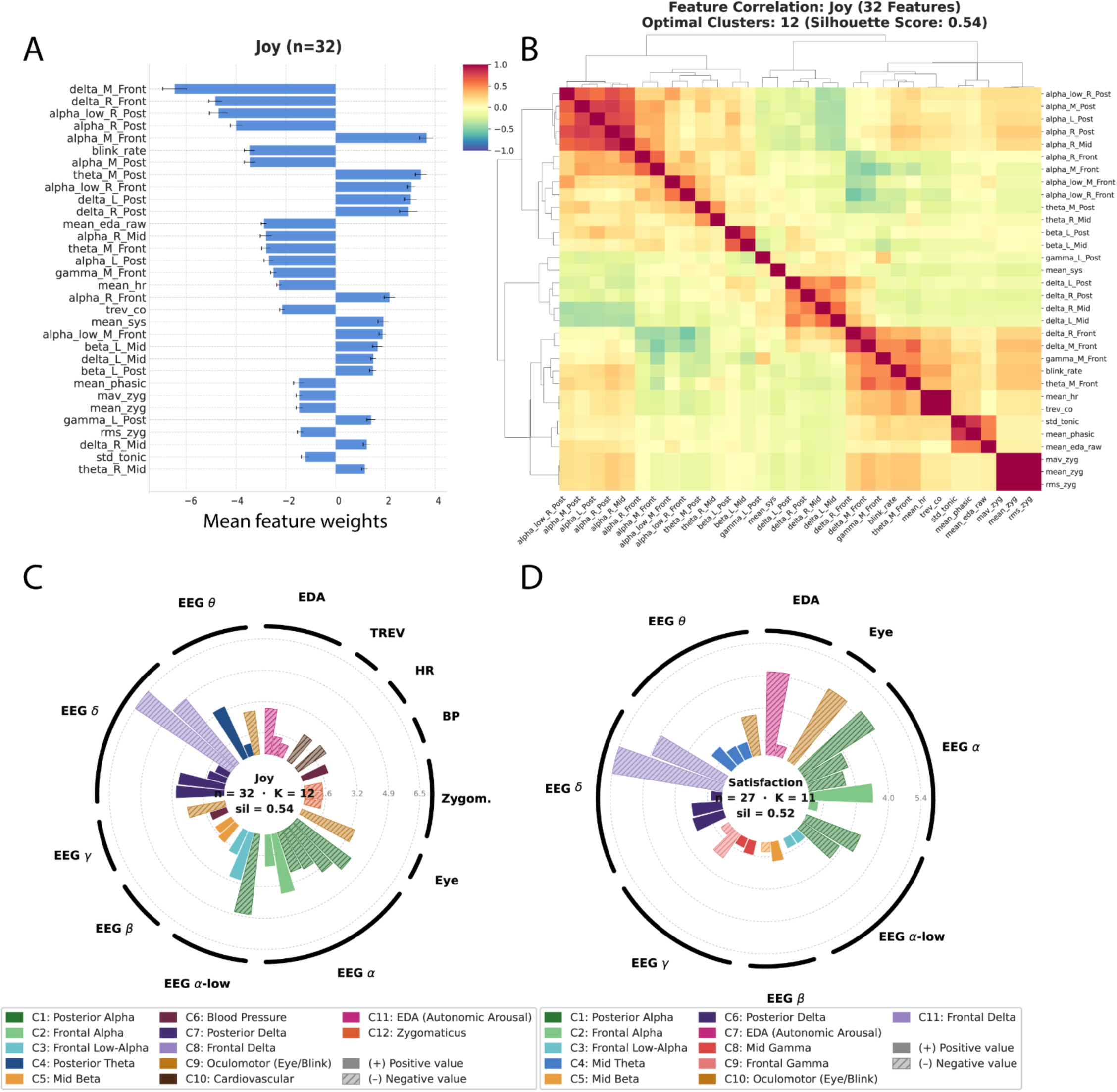
Each emotion recruits a structured, multimodal ensemble of correlated cortical and peripheral features. **A.** Signed mean elastic-net weights of the stable feature set for an example emotion (Joy) in a representative participant (ppt 2). Stable features (non-zero weight in ≥90% of cross-validation runs, trimmed to the top 90% of cumulative absolute importance; n = 32) are ranked by absolute mean weight; error bars show s.e.m. across n = 25 cross-validation runs. **B.** Pearson correlation matrix of the same 32 stable features, computed across the pooled Joy-rated and no-stimulus-baseline samples and reordered by agglomerative hierarchical clustering (Ward’s linkage, Euclidean distance, optimal leaf ordering). The number of clusters was chosen to maximize the average silhouette score (12 clusters; silhouette = 0.54), yielding interpretable functional and anatomical modules. **C.** Circular functional map for Joy. Each stable feature occupies one angular sector; radial extent encodes mean absolute importance, sector color denotes its correlation cluster, hatching denotes negative weights, and outer labels group features by ground truth neurophysiological category. Dashed rings mark fixed fractions of the emotion’s maximum importance; the innermost ring (first quartile) separates dominant from marginal contributors and defines inclusion in Tables 1 and 2. **D.** As in C, for Satisfaction (n = 27 features; 11 clusters, silhouette = 0.52). Satisfaction retained the EDA, blink-rate and cortical delta, alpha and low-alpha clusters seen for Joy but lost its cardiovascular, blood-pressure and zygomaticus features, whereas gamma formed distinct mid and frontal clusters only for Satisfaction.

**Table 1.** Shared and emotion-specific feature clusters in participant 2. Feature clusters (rows) × emotion categories (columns) for participant 2. Only features exceeding the first-quartile importance threshold defined in Fig. 3C were tabulated. Each cell gives the number of contributing features and the signed importance of the feature with the largest absolute magnitude in that cluster; rows are ordered by largest absolute importance, cells are shaded on a diverging color scale, and empty cells indicate clusters below threshold. A shared backbone appeared across emotions: frontal delta was negative for all 15 emotions (and among the strongest predictors), posterior alpha negative for 14/15, posterior delta positive for all 15, and midline theta and frontal (low-)alpha positive for most; the electrodermal cluster was negative for every emotion and the oculomotor cluster appeared in nine. Discriminative information came from near-exclusive clusters: posterior low-alpha for Calmness, posterior theta for Joy, frontal theta for Adoration, and TREV and respiration rate for Awkwardness. Because models were trained separately per emotion, weight magnitudes are not directly comparable across columns.

|  | Admiration | Adoration | Aesthetic Appreciation | Amusement | Awe | Awkwardness | Boredom | Calmness | Confusion | Entrancement | Excitement | Interest | Joy | Relief | Satisfaction |
| --- | --- | --- | --- | --- | --- | --- | --- | --- | --- | --- | --- | --- | --- | --- | --- |
| Frontal Delta | -3.46 (n=2) | -1.27 (n=3) | -8.58 (n=5) | -4.18 (n=3) | -3.68 (n=2) | -3.29 (n=3) | -3.40 (n=2) | -8.07 (n=2) | -3.59 (n=2) | -8.94 (n=2) | -3.83 (n=2) | -5.19 (n=3) | -6.47 (n=2) | -4.30 (n=2) | -5.35 (n=2) |
| Frontal Alpha | +1.83 (n=2) |  |  | +1.79 (n=2) | +2.67 (n=1) |  | +2.15 (n=2) | +4.23 (n=2) | +1.91 (n=2) | +6.16 (n=1) | +2.29 (n=2) | +2.87 (n=2) | +3.66 (n=2) | +1.92 (n=2) | +3.36 (n=1) |
| Posterior Alpha | -2.92 (n=6) | -1.49 (n=6) | -6.15 (n=5) | -3.10 (n=7) | -3.30 (n=5) | -2.76 (n=7) | -2.95 (n=7) |  | -2.21 (n=5) | -5.93 (n=5) | -3.31 (n=7) | -3.46 (n=6) | -4.70 (n=5) | -3.10 (n=5) | -4.27 (n=5) |
| Posterior Low-Alpha |  |  |  |  |  |  |  | -5.75 (n=5) |  |  |  |  |  |  |  |
| Frontal Low-Alpha |  |  | +5.68 (n=1) | +1.89 (n=2) | +1.98 (n=1) | +2.11 (n=4) | +2.19 (n=1) | +3.13 (n=2) | +1.95 (n=1) | +2.66 (n=1) | +2.22 (n=2) | +1.53 (n=2) | +3.05 (n=2) |  |  |
| Posterior Delta | +1.66 (n=2) | +0.63 (n=4) | +4.84 (n=2) | +1.93 (n=4) | +2.03 (n=3) | +1.77 (n=5) | +1.88 (n=5) | +3.97 (n=4) | +0.98 (n=4) | +4.08 (n=2) | +1.58 (n=4) | +2.45 (n=4) | +3.02 (n=3) | +2.06 (n=2) | +2.05 (n=2) |
| EDA (Autonomic Arousal) | -2.80 (n=2) | -1.14 (n=5) | -4.09 (n=1) | -2.62 (n=4) | -4.04 (n=1) | -1.77 (n=4) | -2.52 (n=1) | -2.74 (n=2) | -2.15 (n=1) | -3.80 (n=1) | -3.01 (n=3) | -3.70 (n=3) | -2.88 (n=1) | -1.91 (n=1) | -4.18 (n=2) |
| Oculomotor (Eye/Blink) | -2.55 (n=3) |  |  | -2.82 (n=3) | -2.81 (n=2) |  |  | -4.10 (n=3) |  | -4.05 (n=2) | -3.07 (n=3) |  | -3.46 (n=3) | -2.59 (n=1) | -4.17 (n=2) |
| Frontal Gamma |  |  |  |  | -2.07 (n=1) | -2.71 (n=4) | -2.82 (n=3) |  | -2.45 (n=4) |  |  | -3.80 (n=4) |  |  | -1.94 (n=1) |
| Posterior Theta |  |  |  |  |  |  |  |  |  |  |  |  | +3.43 (n=1) |  |  |
| Blood Pressure |  |  |  |  |  | +1.28 (n=4) |  | +2.59 (n=2) | +0.97 (n=1) | +3.05 (n=1) | +1.33 (n=1) | +1.14 (n=2) | +1.93 (n=1) |  |  |
| Mid Theta | +1.75 (n=4) | +0.90 (n=5) |  | +1.91 (n=3) | +1.62 (n=2) | +1.40 (n=4) | +1.69 (n=3) | +2.55 (n=1) | +1.23 (n=2) |  | +2.38 (n=3) | +2.05 (n=4) |  | +1.45 (n=1) | +1.79 (n=3) |
| Cardiovascular |  | -0.78 (n=3) |  | -1.65 (n=2) | -1.28 (n=1) |  | -0.87 (n=2) |  | -1.64 (n=2) |  | -1.54 (n=3) | -1.70 (n=2) | -2.27 (n=2) |  |  |
| Mid Beta |  | +0.69 (n=2) |  | +1.53 (n=1) | +1.48 (n=1) | +1.47 (n=3) |  |  |  |  | +1.99 (n=1) | +2.05 (n=3) | +1.69 (n=2) | +1.44 (n=1) | +1.54 (n=1) |
| Mid Gamma |  | +0.67 (n=2) |  | +1.68 (n=2) | +1.54 (n=1) | +0.80 (n=1) | +0.83 (n=1) |  |  |  | +1.49 (n=2) | +1.14 (n=2) |  |  | +1.32 (n=1) |
| TREV |  |  |  |  |  | -1.43 (n=2) |  |  |  |  |  |  |  |  |  |
| Frontal Theta |  | -1.39 (n=5) |  |  |  |  |  |  |  |  |  |  |  |  |  |
| Posterior Beta |  |  |  |  |  |  | +0.98 (n=2) |  | +1.17 (n=2) |  |  |  |  |  |  |
| Corrugator |  | -0.55 (n=2) |  |  |  |  | -1.14 (n=1) |  | -0.96 (n=1) |  |  |  |  |  |  |
| Zygomaticus |  | -0.93 (n=5) |  |  |  |  | -0.82 (n=1) |  |  |  |  |  |  |  |  |
| Respiration Rate |  |  |  |  |  | +0.74 (n=3) |  |  |  |  |  |  |  |  |  |

**Table 2.**
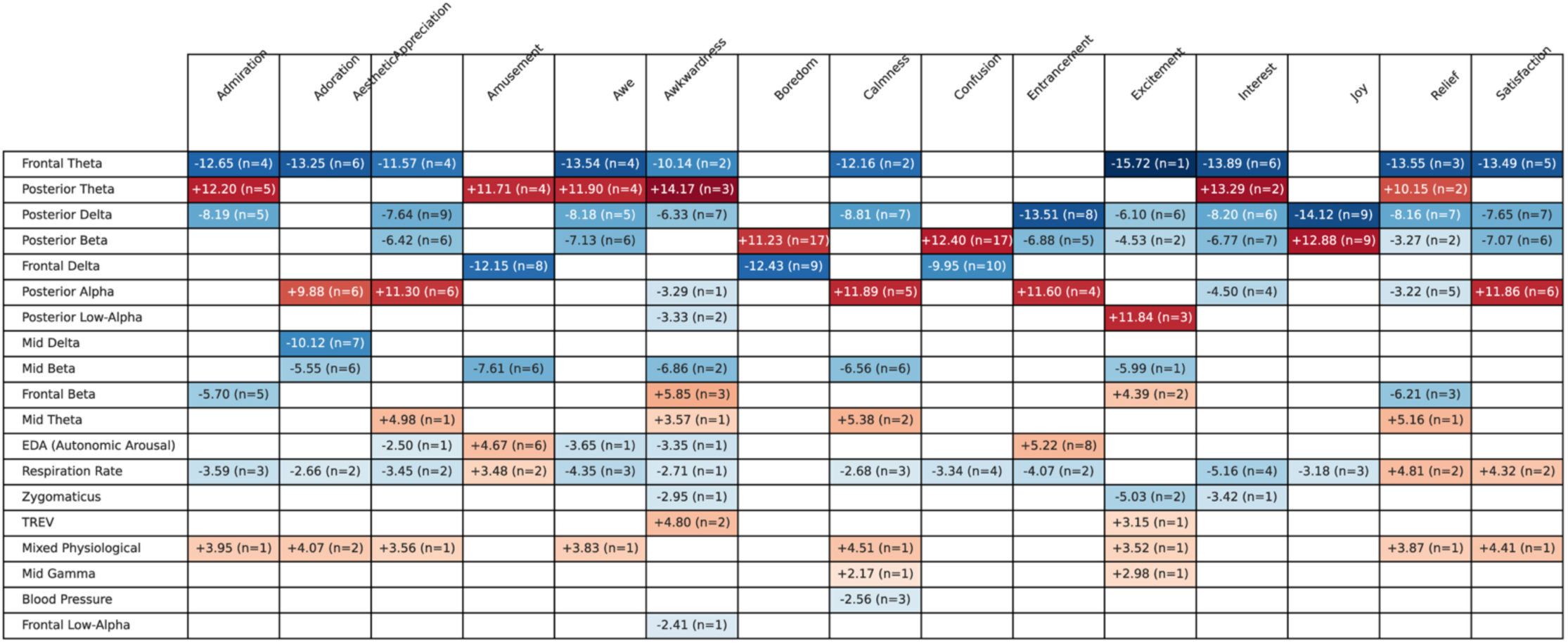
Shared and emotion-specific feature clusters in participant 3. As in Table 1, for participant 3. The shared backbone differs from that of ppt 2: frontal theta was negative wherever it appeared (10/15 emotions), posterior delta negative for 11/15 and posterior theta uniformly positive. Several clusters were near-exclusive (mid delta for Adoration; posterior low-alpha for Awkwardness and Excitement; blood pressure for Calmness; frontal low-alpha for Awkwardness; mid gamma for Calmness and Excitement), and others recurred with emotion-selective sign reversals: such as EDA was positive for Amusement and Entrancement but negative for Awe, Awkwardness and Aesthetic Appreciation; refer to Results for more details. Such sign flips were absent in ppt 2. Corresponding tables for ppt 1 and ppt 4 are provided in Supplementary Tables T3 and T4.

Because many predictors index the same physiological process (e.g., neighboring electrodes and bands, or phasic and tonic EDA), we next examined their covariance structure. For each emotion, the Pearson correlation matrix of stable features across pooled emotion and no-stimulus baseline samples was clustered with Ward linkage, with the number of clusters chosen to maximize the silhouette score (Fig. 3B). The clusters formed interpretable functional and anatomical modules. For Joy, these included frontal and posterior alpha, low-alpha, theta, left beta, frontal and posterior delta and mid-frontal gamma. They also included blink-rate, electrodermal, zygomaticus and cardiovascular (heart rate and contractility) modules. Discriminative information was thus organized into a few correlated sources rather than 32 independent predictors.

To compare this organization across emotions, we projected each emotion’s stable features onto a circular map in which features are grouped by data-driven cluster and color-coded by modality (Joy, Fig. 3C; Satisfaction, Fig. 3D). Two observations stand out. First, each emotion recruited a multimodal signature. For Joy, band-specific EEG clusters were accompanied by several peripheral clusters: electrodermal (raw, tonic and phasic), cardiovascular (heart rate and TREV, a proxy for radial blood flow), blood pressure, zygomaticus and blink rate. Second, the number and composition of clusters shifted by emotion. Joy resolved into 12 clusters (32 features; silhouette = 0.54) and Satisfaction into 11 (27 features; silhouette = 0.52). Satisfaction retained EDA, blink-rate and cortical delta, alpha and low-alpha clusters similar to Joy, but its cardiovascular, blood-pressure and zygomaticus features were no longer stable. These differences indicate partly emotion-specific physiological fingerprints built on a shared multimodal backbone.

We separated low alpha (∼8–10 Hz) from upper alpha (∼10-12 Hz, referred to as alpha), because low alpha may track tonic alertness and non-specific attentional demands, whereas upper alpha is modulated by task-specific processing^28^. Posterior alpha power also varies inversely with external visual engagement, desynchronizing during active viewing and synchronizing when attention is internally directed or external input is gated^29^. Gamma formed distinct mid and frontal clusters for Satisfaction. In Joy, gamma features were absorbed into other clusters including blood pressure and oculomotor. In both emotions, gamma’s contribution (dashed circles) was small relative to other bands. Clustered feature maps for each emotion are shown for an example participant (Ppt 2) in Figure 4 and for other participants in Supplementary Figs. S7–S9. Together, Figs. 3 and 4 show that decoding relied not on an isolated marker but on a multimodal ensemble of correlated EEG and peripheral features, organized into a few interpretable clusters whose membership varied across emotions.

**Figure 4.**
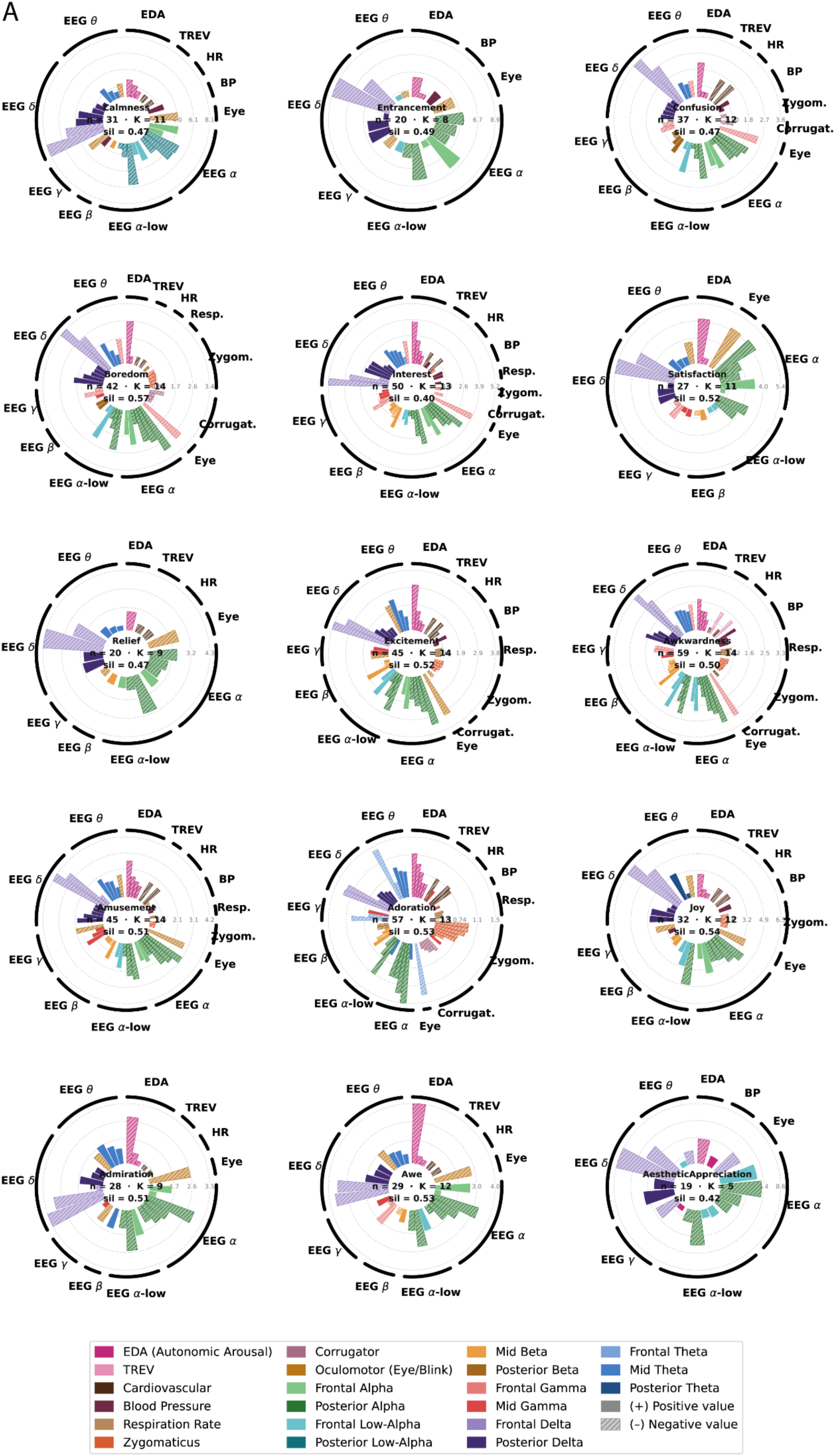
Emotion-specific circular feature maps across the full emotion set in a representative participant. Circular functional maps for all 15 emotion categories in a representative participant (ppt 2), plotted as in Fig. 3 C, D. Radial extent encodes mean absolute elastic-net importance, sector color the data-driven correlation cluster and outer labels the ground truth neurophysiological modality; the innermost dashed ring marks the first quartile of each emotion’s importance range, and the number of stable features (n), clusters (K) and silhouette score (sil) are given for each map. Because models were trained separately for each emotion, radial magnitudes are comparable within but not across maps. Corresponding maps for the other participants are shown in Supplementary Figs. S7–S9.

### Shared versus emotion-specific neurophysiological features

Aggregating stable features across emotions mapped which clusters matter for which emotions (Fig. 4 and Table 1&2 for Ppts 2 & 3; Supplementary Figs. S7-S9 and Tables T3-T4). This separates common features, likely reflecting generic processes (sensory engagement, attention, cognition, arousal), from features specific to one or a few emotions. For Ppt 2, frontal delta was negative for all 15 emotions (and among the strongest predictors), posterior alpha was negative for 14/15, posterior delta was positive for all 15, and midline theta and frontal (low-)alpha were positive for most. Peripherally, the electrodermal cluster was negative for every emotion and the oculomotor cluster appeared in nine. This shared cortical– autonomic pattern has little discriminative power and instead marks the transition from the no-stimulus baseline into an affectively engaged state.

Discriminative information came from a few emotion-specific clusters. Posterior low-alpha appeared only for Calmness, posterior theta only for Joy, frontal theta only for Adoration, and TREV and respiration rate only for Awkwardness.

Corrugator and zygomaticus contributed mainly to Boredom and Adoration, with corrugator alone for Confusion. Blood-pressure and cardiovascular clusters were recruited chiefly by higher-arousal states (Excitement, Interest, Joy), while blood pressure also contributed to low-arousal states (Calmness, Confusion, Entrancement). Note that weight magnitudes are not comparable across emotions, as models were trained separately.

For Ppt 3 (Table 2), frontal theta was negative wherever it appeared (10/15 emotions), posterior delta was negative for 11/15, and posterior theta was uniformly positive. With near-ubiquitous posterior theta/beta, frontal-theta, posterior delta and respiration-rate engagement, this backbone marks general affective engagement. Near-exclusive clusters included mid delta (Adoration), posterior low-alpha (Awkwardness, Excitement), blood pressure (Calmness), frontal low-alpha (Awkwardness) and mid gamma (Calmness, Excitement). Other clusters were emotion-selective and sign-specific. EDA was positive for Amusement and Entrancement but negative for Awe, Awkwardness and Aesthetic Appreciation, and zygomaticus and TREV contributed to few states. Posterior alpha was positive for Adoration, Aesthetic Appreciation, Calmness, Entrancement and Satisfaction but negative for Awkwardness, Interest and Relief. Respiration rate and posterior beta were mostly negative but positive for Amusement, Relief and Satisfaction, and for Boredom, Confusion and Joy, respectively; this sign flip was absent in Ppt 2. Thus, each emotion is defined not by a unique marker but by a shared multimodal backbone plus a compact emotion-specific overlay, whose membership and sign vary with emotion category and individual.

### Neurophysiology and self-report converge on emotion groupings

We next asked whether emotions can be stratified by their neurophysiology. For each emotion we ranked its stable features by LR-EN weight magnitude, computed Spearman rank correlations between every pair of emotions, and clustered the resulting matrix with Ward linkage (Fig. 5A; 79 stable features for ppt 2). For the example participant, at level 4 of clustering 4 groups of emotions emerged:1. Excitement, Interest, Awkwardness, Amusement (with L5, called group 1-A), and Relief, Satisfaction, Admiration and Awe; 2. Adoration; 3. Entrancement and Aesthetic Appreciation, and 4: Boredom, Confusion, Calmness and Joy. While this clustering could reflect emotion characteristics such as arousal (in group 1-A vs. group 4), it demonstrates more nuanced and complex patterns (Supplementary Figure 10, participant 1) where awkwardness, joy and confusion were grouped together.

**Figure 5.**
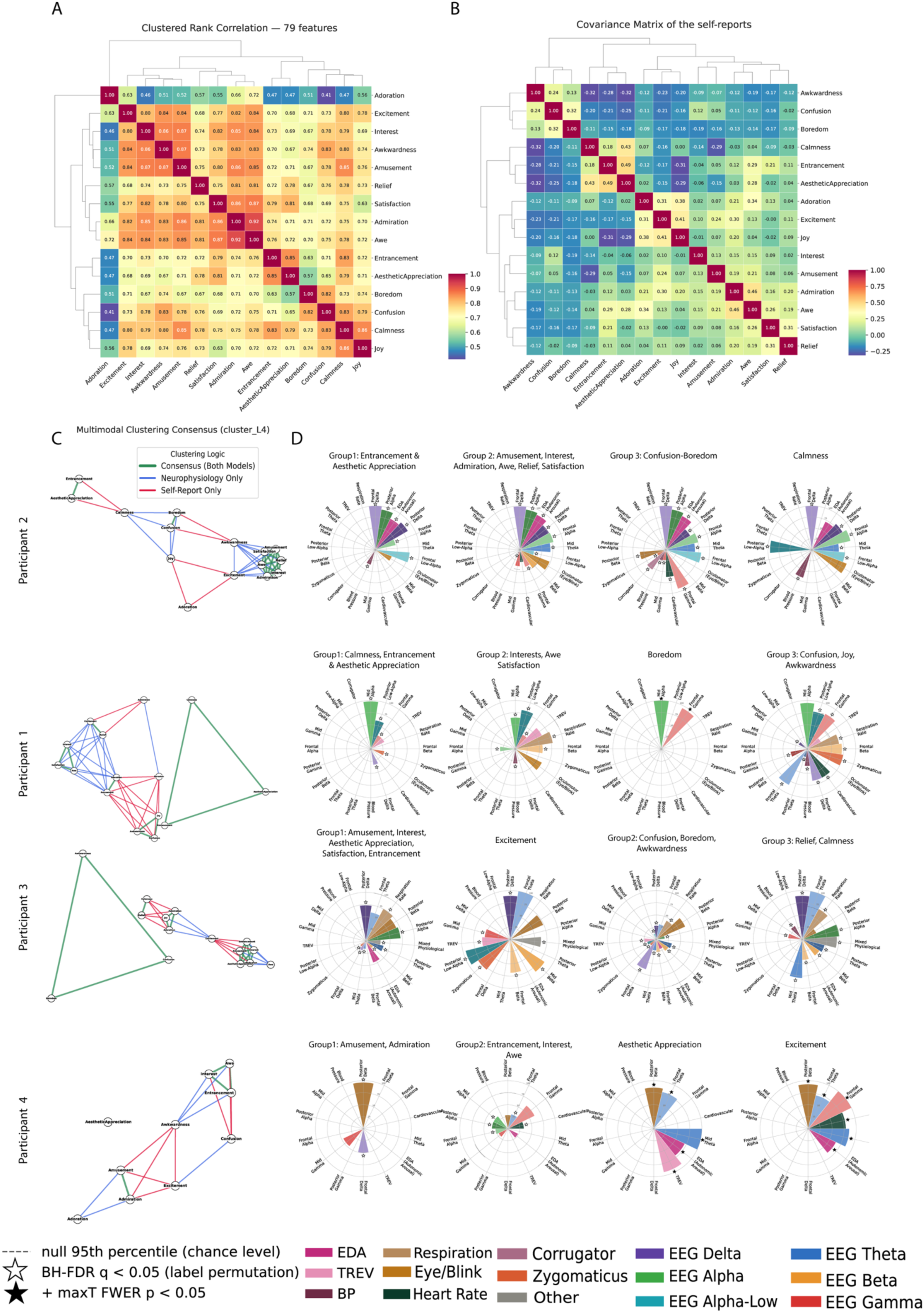
Neurophysiological and self-report emotion structures converge selectively within each participant. **A.** Neurophysiological emotion similarity for a representative participant (ppt 2; n = 79 stable features pooled across emotions). For each emotion, stable features were ranked by absolute elastic-net weight over a common feature space (absent features ranked one step beyond the maximum), Spearman correlations were computed between emotions, and the matrix was clustered with Ward’s linkage and cut at four depths (L2–L5). Correlations are uniformly high (r ≈ 0.4–0.9), reflecting the shared engagement backbone (Table 1). **B.** Self-report emotion similarity for the same participant, from the identical procedure applied to binarized trial-by-trial ratings. Correlations are low and often negative (r ≈ −0.3 to +0.5), separating a positive-appraisal family (Adoration, Excitement, Joy, Interest, Amusement, Admiration, Awe, Satisfaction, Relief) from a disengaged family (Awkwardness, Confusion, Boredom). **C.** Multimodal consensus network for each participant (ppt 1–4). Emotions are nodes; pairs co-clustered at L4 are joined by thick green (both modalities), blue (neurophysiology only) or red (self-report only) edges. For ppt 2, consensus edges formed a clique of positive, high-engagement states (Amusement, Satisfaction, Awe, Relief, Interest, Admiration) and two smaller groups (Entrancement-Aesthetic Appreciation; Boredom-Confusion), whereas Adoration, Joy and Calmness had mainly single-modality edges. **D.** Nightingale rose charts showing the physiology of a selected emotion group per participant (rows, ppt 1–4). Within each emotion, stable features were awarded Borda points (number of features − rank + 1; absent features score zero), summed across the group’s emotions and normalized to the maximum attainable score (bar radius, 0– 100%). Angular position follows one global feature-cluster order and color denotes modality, so roses can be compared directly. The dashed ring marks the 95th percentile of a label-permutation null; open stars, Benjamini–Hochberg FDR q < 0.05; filled stars, also max-T family-wise p < 0.05. For ppt 2, the largest wedges of every group were FDR-significant, although none survived family-wise correction: Entrancement-Aesthetic Appreciation was defined by frontal and posterior delta, posterior alpha, frontal low-alpha and alpha as well as EDA and blood pressure; Confusion-Boredom additionally recruited frontal gamma, posterior beta, cardiovascular and corrugator wedges; and Calmness was distinguished by posterior low-alpha and blood pressure. The overlayed dashed sectors is an uncorrected per-wedge reference; wedges exceeding it without a star did not survive FDR correction across all wedges in that participant, such as oculomotor wedge in Calmness for ppt2. Remaining emotion groups are shown in Supplementary Figs. S12–S15.

We next asked whether emotions that recruit similar physiology are also grouped together in self-report, a question that has long divided emotion theories^30,31^. Clustering binarized trial-by-trial ratings the same way gave a purely behavioral grouping (Fig. 5B). The two matrices (Fig. 5A & Fig. 5B) differed sharply: physiological rank correlations were uniformly high (r ≈ 0.4 to 0.9), reflecting the shared “engagement backbone”, whereas self-report correlations were low and often negative (r ≈ −0.3 to +0.5), because participants rarely endorsed opposing states on the same trial. Both nonetheless showed interpretable structure: Entrancement–Aesthetic Appreciation grouped together in both modalities, and behaviorally a positive-appraisal family (i.e., (Adoration, Excitement, Joy, Interest, Amusement, Admiration, Awe, Satisfaction, Relief) separated from a disengaged family (Awkwardness, Confusion, Boredom).

We formalized this agreement with a multimodal consensus network per participant (See Methods for more detail; Fig. 5C), linking emotion pairs co-clustered at a matched level (L4) by green (both modalities), blue (neurophysiology only) or red (self-report only) edges. For ppt 2, green edges formed a dense consensus clique of positive, high-engagement states (Amusement, Satisfaction, Awe, Relief, Interest and Admiration), with two smaller consensus groups, Entrancement-Aesthetic Appreciation and Boredom-Confusion, that are low-arousal state with positive and neutral-negative tendency, respectively. Adoration, Joy and Calmness had mainly single-modality edges, meaning that the participant grouped them subjectively in ways their physiology did not mirror, or vice versa. This selective correspondence recurred, with participant-specific topology, in ppt 1, ppt 3 and ppt 4.

Finally, Nightingale rose plots show the physiology defining selected emotion group for each participant (Fig. 5D; the rest of emotion groups are shown in supplementary figures S12-S15). Each rose is a Borda consensus of elastic-net rankings, tested against label-permuted models (open stars, FDR q < 0.05; filled stars, also max-T family-wise p < 0.05; see Methods). For ppt 2 (Fig 5D, top row), the largest wedges of every group were FDR-significant, although none survived family-wise correction. Entrancement-Aesthetic Appreciation (i.e., Group 1) was defined by significant frontal delta, posterior alpha, EDA, posterior delta, frontal low-alpha and alpha; Group 2 additionally recruited significant mid theta and frontal gamma; Confusion-Boredom showed the broadest significant profile, adding frontal gamma, posterior beta, cardiovascular and corrugator wedges; and Calmness was distinguished by a significant posterior low-alpha and blood pressure wedges. Size alone did not imply reliability: the oculomotor wedge of Group 2 did not exceed the null, nor did the dominant frontal gamma wedge of ppt 4’s Entrancement-Interest-Awe group. Across participants, similar but not identical groups emerged, built on shared dominant features (such as Frontal and posterior delta, EDA and Frontal low-alpha in ppt2) plus significant emotion group-specific ones.

### Emotion neurophysiological groupings that recur across individuals

We next examined whether participants’ neurophysiology converges on the same emotion groupings. For each participant, emotions were clustered purely from physiological data and merged into a consensus graph employing aforementioned approach (Fig. 6A). Each of the 105 possible emotion pairs (i.e., 15 choose 2) was weighted by how many participants clustered them together. Green edges mark agreement in all included participants, black edges in two of three, and colored edges in one only. Participant 4 was excluded from modeling due to smaller sample size for 5/15 emotions (see Supplementary Fig. S16A for results including this participant). Cliques among consensus edges then defined recurring emotion groups.

**Figure 6.**
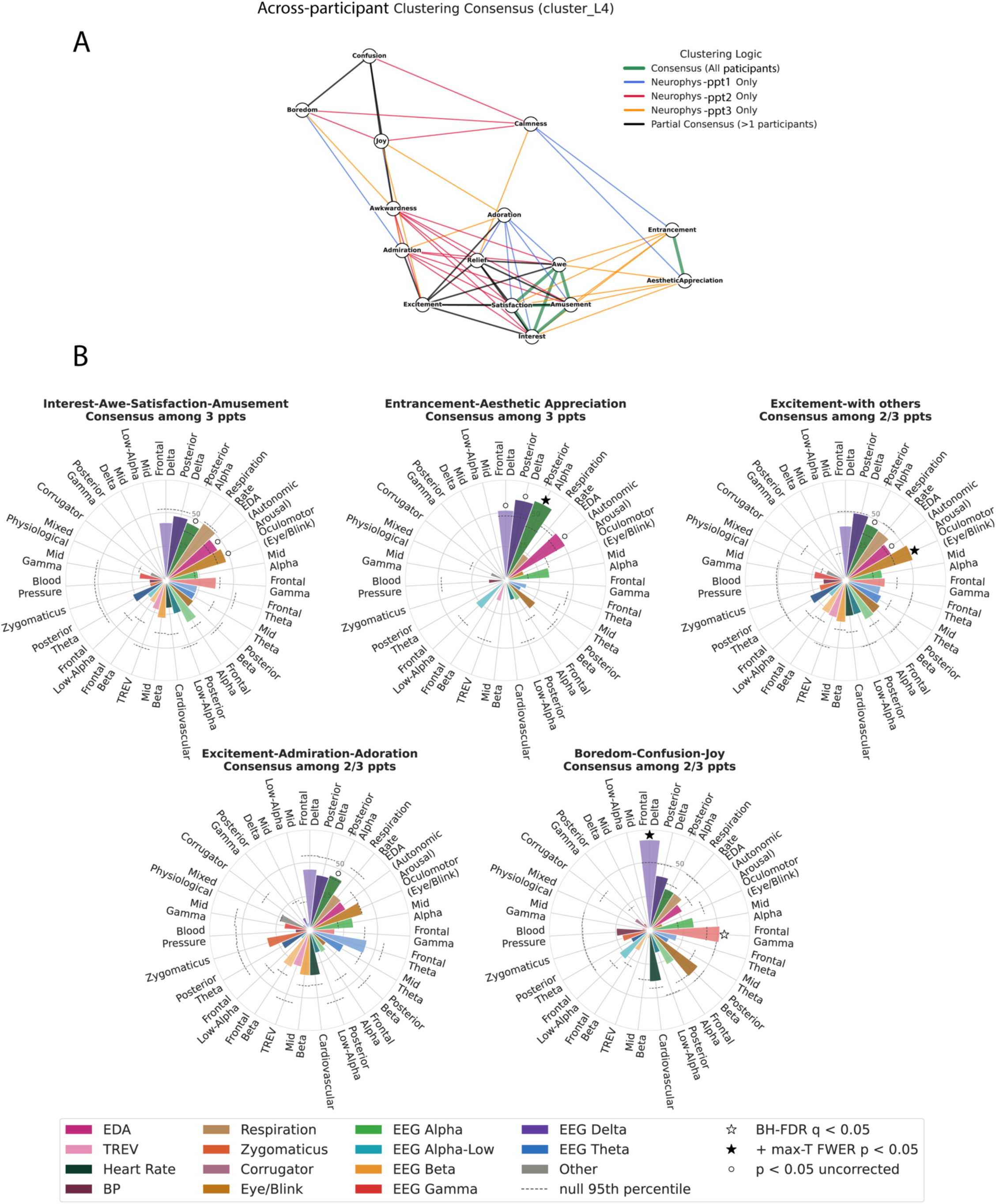
A reproducible neurophysiological scaffold for emotions that recur across individuals. **A.** Consensus network based on neurophysiology. Emotion partitions were derived independently for each participant from neurophysiology alone (Ward clustering of the Spearman rank-correlation matrix of stable elastic-net feature orderings, cut at L4) and entered into a single undirected graph. Each of the 105 possible emotion pairs was evaluated only in participants for whom both emotions existed; thick green edges mark full consensus (all included participants, ppt 1-3), black edges partial consensus (two of three) and participant-colored edges pairings supported by a single participant. Ppt 4 was excluded because only 10 of 15 emotions were modelled (Supplementary Fig. S16A includes this participant). Cliques among consensus edges define the reproducible groups: Interest-Awe-Amusement-Satisfaction and Entrancement-Aesthetic Appreciation reached full consensus, whereas Excitement with others (See Results and Methods), Excitement-Admiration-Adoration and Boredom-Confusion-Joy reached partial consensus. **B.** Nightingale rose plots for each consensus group, extending the Borda procedure of Fig. 5D across participants: each participant’s feature clusters were ranked per emotion and awarded Borda points; clusters supported by at least two of three participants form the consensus set (minority clusters are still drawn), and bar radius gives the percentage of the maximum possible cross-participant score. Significance was assessed with 5,000 within-participant permutations, tested and corrected within each group across the consensus set: dashed arcs, null 95th percentile; open circles, uncorrected p < 0.05 (exploratory); open stars, FDR q < 0.05; filled stars, studentized max-T family-wise p < 0.05. Posterior alpha defined Entrancement-Aesthetic Appreciation and oculomotor activity defined Excitement with others (both family-wise significant), and Boredom-Confusion–Joy was characterized by frontal delta (family-wise) and frontal gamma (FDR); no cluster survived correction in the other two groups, and no single biomarker was significant in every group.

Two groups reached full consensus: Interest-Awe-Amusement-Satisfaction and Entrancement-Aesthetic Appreciation. A larger set reached partial (two-of-three) consensus, extending the first group to Relief and Excitement (Excitement with others) and linking Boredom, Confusion and Joy. Some emotions had a single consensus partner (degree 1): Adoration-Excitement, Admiration-Excitement and Awkwardness-Joy. These recapitulate intuitive affective families: high-engagement positive (Interest, Awe, Satisfaction, Amusement, Excitement), absorbed-aesthetic and possibly low-arousal (Entrancement, Aesthetic Appreciation), and a more neutral, low-arousal cluster (Boredom, Confusion, Joy), although Joy is not usually considered neutral. The remaining edges were participant-specific, showing that much of each individual’s emotion geometry, particularly for Adoration, Calmness, Joy, Boredom and Awkwardness, is idiosyncratic. Because these clusters derive solely from neurophysiology, their convergence reflects shared peripheral–cortical organization rather than shared conceptual structure in self-reports.

To identify each group’s defining features, we extended the Borda procedure across participants (Fig. 6B); bar radius gives each feature cluster’s score as a percentage of the maximum attainable. Significance was tested by 5,000 within-participant permutations, corrected within each group (See Methods). Posterior alpha defined Entrancement–Aesthetic Appreciation (max-T FWER p < 0.05), with frontal delta, posterior delta and EDA reaching nominal significance (uncorrected p < 0.05). Oculomotor activity defined Excitement with others (FWER p < 0.05), with nominal support from posterior alpha and EDA. Boredom-Confusion-Joy was characterized by frontal delta (FWER p < 0.05) and frontal gamma (FDR q < 0.05). In Interest-Awe-Satisfaction-Amusement and Excitement-Admiration-Adoration, no cluster survived correction. In the first of these, posterior delta, EDA and oculomotor activity reached only nominal significance; in the second, only posterior alpha did. Frontal or posterior delta, posterior alpha, electrodermal and oculomotor clusters therefore recurred across several groups. Yet no single biomarker reached significance in all of them, and each group was distinguished by a different combination of features. Cardiovascular modules contributed more variably and were the chief source of individual differences. Note that these tests establish cross-participant agreement, not group specificity (Supplementary Table T5). Together, these results indicate that shared emotion groupings rest on a partly common neurophysiological profile, while the specific biomarkers supporting each group remain substantially idiosyncratic.

## Discussion

We combined dense, comprehensive within-individual multimodal biosensing data with interpretable machine learning to ask how discrete emotions are instantiated in the brain and body of a single person. Within each participant, all 15 emotion categories were separable from a no-stimulus baseline, with mean accuracy across emotions ranging from 0.87 (ppt4) to 0.97 (ppt1), well above permutation surrogates (mean accuracy 0.4993–0.501; Supplementary Table T2). Three results deserve emphasis. First, no emotion was carried by an isolated marker; each was defined by a multimodal ensemble of cortical oscillatory and autonomic features. Second, the composition of that ensemble was largely person-specific. Third, despite this idiosyncrasy, a small scaffold of cortical and autonomic modules recurred across individuals, indicating that personalized characterization and population-level structure are complementary rather than competing goals.

Affective processes are time-varying and non-ergodic: within-person variance can be two to four times larger than between-person variance, so population averages often fail to describe any single individual^10,11^. This has slowed biomarker development in psychiatry and motivated person-specific, high-precision models of psychopathology^6,7,32–34^. Intracranial studies show that affective states are encoded idiosyncratically^24,35,36^, and successful biomarker-driven interventions, such as closed-loop neuromodulation for treatment-resistant depression, have relied on person-specific electrophysiological signatures^37–41^. Precision mapping shows that individual brain networks combine group-level and individual-specific features^12,13^, and dense small-N designs are a principled route to such inference^14^, yet most multimodal affective datasets favor participant numbers over within-person depth^25,26^. Because our 102 cortical and autonomic features are mutually correlated, we used elastic-net regularization^42^, which selects sparse, grouped and interpretable predictors where mass-univariate tests would be underpowered.

An emotional episode unfolds over seconds to minutes and is shaped by regulation^43,44^, physiological background and recent events, so the same nominal emotion varies across occasions and people; this is the central obstacle to physiological markers of emotion. Autonomic specificity emerges only once emotion subtypes are distinguished^1^, effect sizes are heterogeneous within every category^8^, positive emotions show graded differentiation ordered by approach motivation^15^, and ambulatory sampling reveals substantial within-category variation in daily life^9^. Emotional experience is supported by distributed, partly valence-general networks acting with the autonomic nervous system, yet discrete categories remain recoverable from distributed patterns^17–19,21^ and are associated with distinct maps of bodily sensation^45^. However, supervised and unsupervised analyses can yield discordant category structure^46^, so classifier success is not proof of natural kinds. Our design is agnostic to the debate between basic-emotion^2,47,48^ and dimensional or constructionist accounts^30,49,50^: we assume neither one-to-one mappings between labels and physiology nor a two-dimensional space, which our predominantly positive stimuli could not span. The semantic-space work underlying our stimuli reports at least 27 categories bridged by continuous gradients^20^, and research on discrete positive emotions argues for differentiated functions within positive affect^51^. Our data add that a similar pattern appears within a single person: a large, shared component behaving like a general engagement dimension, with a compact, category-selective overlay.

LR-EN models matched other model families while providing interpretable, signed weights (Fig. 2), and clustering each model’s stable features yielded anatomically and functionally interpretable modules (Fig. 3). In the example participant, Joy recruited frontal delta and alpha and posterior alpha alongside every recorded peripheral channel, whereas Satisfaction retained this cortical backbone but lost the cardiovascular, blood-pressure and zygomaticus modules and gained gamma clusters (Figs. 3, 4). Across emotions (Tables 1, 2; Supplementary Tables T3, T4), each emotion emerged from a widely shared neurophysiological foundation plus a compact overlay, typically one or two cortical bands (e.g., mid delta, posterior low-alpha) and specific autonomic or somatic modules (blood pressure, respiration, EDA, facial EMG), whose composition and sign varied systematically with emotion. Because decoding weights reflect noise suppression as well as signal^52^, we interpret clusters rather than individual weights.

A fundamental question in affective neuroscience is how neurophysiology relates to subjective report^17,46^. We compared a neurophysiological geometry of each participant’s emotional repertoire, derived from feature rankings (Fig. 5A), with an independent behavioral geometry from self-reports (Fig. 5B). Physiological similarities were uniformly high, reflecting the shared engagement backbone, whereas self-report similarities were low and often negative because participants rarely endorsed opposing states on the same trial. The consensus network showed selective rather than global agreement (Fig. 5C), concentrated in a high-engagement positive core (Amusement, Satisfaction, Awe, Relief, Interest, Admiration) and two smaller lower-arousal families, Entrancement-Aesthetic Appreciation and the disengaged-neutral Boredom-Confusion, whereas Adoration, Joy and Calmness were grouped by one modality only. This structure echoes fMRI mapping of the same video library, in which emotion categories predicted cortical and subcortical responses better than valence or arousal and responses showed a cluster-like categorical organization^20,21^, a view formalized in semantic space theory^22,53^ and in work mapping dozens of categories onto distributed cortical space^19,54^. A reanalysis of the same fMRI data, however, emphasizes substantial within-category variation^55^, and large EEG datasets now separate finer-grained positive categories such as amusement, inspiration, joy and tenderness^56^. Yet our groupings did not reduce to arousal: the high-engagement core combined high-arousal Amusement and Interest with lower-arousal Satisfaction and Relief, consistent with evidence that positive states are differentiated more by approach motivation than by arousal^57^. Because the maps were derived independently, this selective convergence, which recurred in every participant, is a genuine cross-modal check.

Within individuals, the features defining each family were statistically reliable but idiosyncratic (Fig. 5D; Supplementary Figs. S12–S15). In ppt 2, the largest wedges of every group exceeded label-permuted models (FDR q < 0.05): Entrancement-Aesthetic Appreciation relied on frontal and posterior delta, posterior and frontal (low-)alpha and EDA; Confusion-Boredom showed the broadest profile, adding frontal gamma, posterior beta, cardiovascular and corrugator features; and Calmness was distinguished by posterior low-alpha and blood pressure. Group membership itself also differed, with Entrancement joining Interest and Awe in ppt 4 rather than Aesthetic Appreciation. This fits meta-analytic evidence that categories are partly conserved in what they group together but idiosyncratic in neurophysiological channels that realize them^1,8,20^, as expected under non-ergodicity^10^, a group-average model would pool ppt1’s mid alpha-dominant and ppt4’s EDA-dominant implementations of the same emotional family (Aesthetic Appreciation) and recover neither.

Given that each participant’s emotion geometry was largely individual, we next asked whether any of it generalizes across people. Across participants, two emotion groups reached full (3/3 participants) and three partial consensus (Fig. 6A), with profiles combining a shared vocabulary and group-specific markers (Fig. 6B, Participant 4 to see their results see Supplementary Fig. S16). Posterior and frontal Delta, posterior alpha and electrodermal clusters recurred most consistently, suggesting a common vocabulary for affective and sensory engagement, whereas blink rate, facial-muscle and cardiovascular modules contributed variably and were the main source of individual differences; zygomaticus and heart-rate variability, for example, were absent from Entrancement-Aesthetic Appreciation. Three groups had a significant defining marker: posterior alpha for Entrancement-Aesthetic Appreciation, oculomotor activity for Excitement with other emotions, and frontal delta and gamma for Boredom-Confusion-Joy. Prior scalp work linked awe-related positive emotions to fronto-central alpha and beta, interest and amusement to theta, and serenity to increased parieto-occipital alpha^23^; our data converges on alpha involvement and is consistent with posterior alpha and low-alpha association in absorbed states.

Alpha is not a unitary phenomenon. Lower alpha (∼8–10 Hz) tracks tonic alertness and general attentional demands, upper alpha (∼10–12 Hz) reflects task-specific processing, and posterior alpha decreases with external visual engagement and increases when attention is internally directed or external input is gated^28,29^. Interest and Awe, which engage an information-rich external scene, should therefore rely less on posterior alpha than the absorbed, internally weighted Entrancement and Aesthetic Appreciation, and posterior alpha was indeed the only significant marker of the absorbed group. This aligns with imaging evidence that intense aesthetic experience engages the default-mode network, signaling self-relevance^58,59^, whereas awe reduces default-mode activity and engages fronto-parietal regions directed at the external scene^60^. Peripherally, the absorbed group showed weak oculomotor and respiratory contributions, whereas excitement-anchored groups relied on blink rate, which marks attentional shifts during video viewing^61^. Peripheral meta-analysis and behavioral studies likewise show greater sympathetic reactivity for excitement than for awe or contentment^15,51,62^. Two caveats apply: alpha peak frequency and topography vary between and within individuals^63^, and our 3 × 3 reduction averages left, medial and right regions, so frontal-posterior contrasts should be read as a broad anterior–posterior gradient rather than as specific cortical sources.

Frontal gamma was a significant marker of Boredom-Confusion-Joy but contributed little to the excitement-anchored groups, the reverse of an arousal-based prediction^64,65^; gamma may instead index local evaluative processing. Our intracranial work shows gamma increases during affective relative to neutral behaviors throughout the mesolimbic network^24^, orbitofrontal gamma accompanies outcome evaluation in emotional contexts^66^, and gamma is one of the most informative band in scalp emotion classification^67^. In boredom and confusion, a stimulus fails to resolve into something worth attending to^68^, leaving unresolved evaluation alongside internally generated thought, with default-mode engagement^69^ and elevated alpha^70^. Accordingly, the boredom-anchored group lacked an oculomotor component, consistent with internally directed processing under low external engagement. This interpretation requires an explicit caveat. Scalp gamma (30–55 Hz) overlaps the spectrum of cranial muscle activity^71,72^, and miniature saccades generate task-modulated gamma-band artifacts^73,74^. Because facial and eye movements covary with emotion, such artifacts would mimic neural effects. We mitigated this with ICA, spectral parameterization that separates oscillatory from aperiodic components^75^, and by entering zygomaticus, corrugator and electrooculographic features into the same models so that they compete with cortical features for explanatory weight. Nonetheless, the frontal gamma findings are the most provisional of our results, and confirmation with high-resolution eye tracking or intracranial recordings^76^ is warranted.

The no-stimulus baseline is also unlikely to be affectively neutral. Emotional episodes outlast their eliciting stimuli, and the same autonomic states recur during emotional reactivity and undirected rest^77,78^, although the rating epoch preceding each baseline partly buffers this carryover. Task epochs also differ from baseline in audiovisual input: posterior alpha desynchronizes with external visual engagement^29^, low-frequency activity entrains to continuous audiovisual streams^79,80^, blink rate^61^ and EDA track viewing engagement independently of emotion category^27,81^, and visual cortex itself carries emotion-relevant information^82^. Both considerations support reading the shared backbone as marking the transition from rest into a perceptually and affectively engaged state, which is why the sparser overlay clusters, not the backbone, discriminate emotions. The sign of several backbone clusters differed between participants (Tables 1, 2; Supplementary Tables T3, T4), consistent with individual tonic set points. Importantly, residual baseline emotion should reduce rather than inflate separability, and shared sensory drive cannot explain why emotions differ. Stimulus-matched neutral clips and jittered washouts could test whether the preceding emotion is decodable from subsequent rest, a person-level measure akin to emotional inertia, which predicts depression and maladjustment^83^.

Several further limitations apply. Dense sampling means few participants, so the cross-participant consensus shows that reproducible structure exists rather than its full extent. The stimulus library^20^ evoked predominantly positive, low-threat states, so generalization to negative and threat-related emotions, for which most of the autonomic literature was developed^1,8^, remains to be established. Video elicitation lacks the temporal and social structure of naturalistic emotion^9,24^, and scalp EEG with a nine-region reduction cannot resolve mesolimbic sources that intracranial work implicates most strongly. Self-reports were collected per block and are retrospective, non-exclusive and language dependent. Finally, distinguishing emotions from one another, rather than from baseline, remains harder and more clinically relevant (Supplementary Fig. S2), owing to the shared backbone and a design in which participants could rate multiple emotions per trial. All analyses are correlational, and the brain–body relationships reported here motivate but do not establish causal accounts.

In summary, interpretable, regularized models trained on many trials from one person, combined with multimodal recording of central and peripheral physiology, can move affective neuroscience from population description toward individualized characterization. Such signatures could support precise characterization and longitudinal monitoring of mood and anxiety disorders^7,41^, and as peripheral and neural sensing becomes available in wearable form^26^, the framework can be carried into the settings where emotion, and its disorders, actually unfold.

## Methods

### Participants and experiment design

Four adult participants each completed four 1-hour sessions of the experimental paradigm, providing 168 one-minute trials per participant. In each trial a participant viewed a short emotionally evocative video selected from a validated emotion-elicitation library^20^ (see Stimulus construction) and subsequently reported the emotion(s) experienced by choosing among 18 positive and neutral emotion categories and rating intensity on a 0-5 scale. Each session comprised 7 blocks of 6 trials, and no-stimulus segments were recorded to provide a baseline condition. All procedures were approved by the University of California, San Francisco Institutional Review Board. Participants provided written informed consent to participate in the experiment. Data collection and analysis were not performed blind to the conditions of the experiments.

### Stimulus construction

Stimuli were assembled from a validated normative library of short emotional video clips (roughly 2,000 of them), each of which comes with normative affect ratings from Cowen et.al.^20^. Using 18 positive and neutral emotions out of 27 distinct emotion categories, we selected the most representative clips for each emotion by ranking all clips in descending order of that emotion’s normative rating and taking the top 20 highest-rated exemplars. Because a clip could rank highly for more than one emotion, overlaps were resolved to enforce mutual exclusivity: any clip appearing in multiple emotions’ lists was retained only for the emotion in which its rating was highest, removed from the others, and each emotion’s list was backfilled with the next-highest-ranked clips, iterating until every emotion had a disjoint set of clips and each clip was assigned to a single emotion. The clips selected for each emotion were then concatenated end to end into that emotion’s stimulus, yielding the ∼1-minute trial videos presented during the experiment; the exact clip composition and ordering of each trial were fixed in advance.

The 18 emotions were: *Calmness, Entrancement, Confusion, Boredom, Nostalgia, Romance, Interest, Satisfaction, Relief, Excitement, Awkwardness, Amusement, Adoration, Joy, Admiration, Craving, Awe, Aesthetic Appreciation. Nostalgia, Romance and Craving* were excluded from subsequent analyses because they form separate clusters outside the smooth gradients of the normative emotion space (see Fig. 2 of ref. ^20^) and were rarely selected by our participants.

### Self-report rating variability

To quantify the inter-participant variability in emotion reporting, two distinct statistical measures were calculated based on the participants’ behavioral rating distributions. First, the raw occurrence counts for each of the fifteen emotions are normalized to the maximum count within each individual participant, standardizing the data to a relative frequency scale ranging from 0 to 1. The **Standard Deviation of Normalized Counts (Count SD)** was then computed for each emotion across all participants, serving as a parametric measure of absolute variance in how frequently an emotion is experienced relative to an individual’s own baseline. To account for potential scaling differences and isolate categorical priorities, a second non-parametric metric, the **Standard Deviation of Ranks (Rank SD)**, was calculated. This was achieved by transforming each participant’s normalized counts into descending ordinal ranks (1 representing their most frequent emotion, 15 their least) and computing the standard deviation of these ranks across the cohort. Together, these metrics provide a robust evaluation of variability in subjective rating among participants. These findings are shown in Extended Table 1.

### Multimodal Bio Sensing (MMBS) data acquisition

Our multimodal biosensing (MMBS) recording combined a 128-channel BioSemi Active 2 EEG system with 28 physiological readouts from BIOPAC and external BioSemi sensors: electrocardiography (3 lead ECG, BIOPAC); electromyography (EMG, External sensors of BioSemi) over the zygomaticus and corrugator muscles; electrooculography (EOG) of vertical and horizontal eye movements; a respiration belt (RSP, BIOPAC); electrogastrography (EGG) of gastric myoelectrical activity; trans-radial electrical impedance velocimetry (TREV, BIOPAC), which provides a non-invasive proxy for relative cardiac contractility and stroke volume from radial blood flow; electrodermal activity (EDA, BIOPAC) as skin conductance; and continuous systolic and diastolic blood pressure recorded with a CNAP RT-BP system. Fig. 1B shows the sensor layout.

### Preprocessing

#### EEG preprocessing

EEG was recorded with a 128-channel BioSemi ActiveTwo system alongside synchronized peripheral physiology as external sensors including EOG, facial EMG, EGG in addition to ECG, EDA, TREV and respiration that were acquired through a BioPAC MP160 system and its Acqknowledge Software. Blood pressure was recorded through CNAP RT-BP system. Preprocessing followed a six-stage pipeline in which each stage operated on the saved output of the preceding one. In the first stage, raw BioSemi EEG (.bdf) and BioPAC (.mat) files were imported and integrated in MATLAB using EEGLAB^84^ with FieldTrip utilities^85^. EEG was downsampled by a factor of four to 512 Hz, detrended, and low-pass filtered at 55 Hz with a zero-phase FIR filter (applied via filtfilt); two zero-phase IIR notch filters (centered at 28 Hz and 48.5 Hz) were additionally applied to suppress CNAP equipment-and line-related noise. Channel locations were assigned from a 128-channel template, and non-EEG channels (EOG, EMG, EGG) were separated into dedicated streams. BioPAC signals were downsampled to 500 Hz and low-pass filtered; ECG R-peaks were detected using the Pan–Tompkins algorithm^86^ to derive heart rate. Event markers from the two acquisition systems were cross-checked for temporal synchronization before behavioral data were merged into a single per-block data structure.

Noisy electrodes were then identified from the imported data using a two-stage procedure: channels whose standard deviation exceeded twice the mean of all channel standard deviation were flagged automatically, and this set was verified through visual inspection of continuous traces and of scalp topographies averaged over five successive time windows. In the next stage, flagged electrodes were removed and reconstructed by spherical interpolation (EEGLAB pop-interp)^84^. The interpolated data were high-pass filtered at 0.5 Hz (zero-phase FIR), yielding an effective analysis band of approximately 0.5–55 Hz, after which a deliberately lenient amplitude-based artifact rejection (±500 μV) removed only gross transients prior to independent component analysis.

To remove eye blinks and horizontal eye movement from the EEG signal, independent component analysis was run separately for each block on the concatenated EEG and EOG data from the previous stage using extended infomax ICA^87^ (runica), the resulting weights were saved and, in the following stage, used to compute component activations for manual review. Components reflecting ocular activity, i.e., eye blinks and horizontal/vertical eye movements, were identified by jointly inspecting component scalp topographies and time courses against the raw EOG-referenced traces, and the indices of the artifactual components were saved. In the final stage, those components were projected out of the data, the cleaned EEG was re-referenced to a common average computed over the retained (non-interpolated) channels, and the data were re-epoched by trial. The resulting clean, common-average-referenced, trial-epoched EEG was saved per block for subsequent analysis. We then applied another amplitude-based artifact rejection (±100 μV) removed and interpolated only gross transients. Because baseline period contained more sudden movement transients, 2.5 x mean of standard deviation across all cleaned signals were chosen as the threshold for amplitude-based artifact rejection. This pipeline was adapted from our previous study^88^. All preprocessing steps were programmed in MATLAB (2024a).

#### Peripheral physiology preprocessing

Electrodermal activity was decomposed into phasic and tonic components using the cvxEDA convex-optimization model ^89^, and low-frequency EGG and heart-rate-variability spectra were estimated by Morlet wavelet decomposition. Data were then epoched by trial, with epochs temporarily concatenated within each block for the subsequent decomposition.

### Feature extraction

#### Spectral feature extraction

First, the continuous EEG signal for each trial and electrode was divided into 10-s segments with 50% overlap, which augmented the dataset for model training. Power spectral density (PSD) was computed for each segment using a multitaper fast Fourier transform (mtmfft), and the FOOOF algorithm^75^ was used to isolate oscillatory power by removing the aperiodic (fractal) component. For each 128 channels, spectral power was then averaged within six canonical bands: delta (1-4 Hz), theta (4-8 Hz), low alpha (8 - 10 Hz), alpha (10 - 12 Hz), beta (13 - 30 Hz) and gamma (30 - 50 Hz). Finally, band power was averaged across good electrodes within each region of a 3 × 3 montage spanning the anterior-posterior and left-right axes (left-, mid- and right-frontal; left-, mid- and right-medial; left-, mid- and right-posterior; Supplementary Figure 1), yielding 54 spectral features (9 regions × 6 bands) per 10-s segment.

#### Peripheral physiology feature extraction

Peripheral signals contributed 48 features summarizing heart rate (mean and variability); tonic and phasic EDA (mean and standard deviation of raw, tonic and phasic components); systolic and diastolic blood pressure (mean, maximum, minimum, standard deviation and kurtosis); respiration (rate plus waveform-length, average-amplitude-change, maximum-fractal-length, mean-absolute-value and root-mean-square statistics); EGG (the same waveform statistics); blink rate; zygomaticus and corrugator EMG (mean, maximum, minimum, standard deviation, kurtosis, variance of the difference, mean-absolute-value and root-mean-square); and radial blood-flow (TREV) metrics. Features were computed within overlapping (50%) 10-s segments and summarized per trial. Trials or segments containing missing values were removed prior to modeling. In total, 102 features were extracted per window: 54 spectral features (9 regions × 6 bands) and 48 peripheral features. The full feature set is listed in Supplementary Table T1.

### Block-wise normalization and cross validation

To control for slow drift and session-to-session differences, task epochs (when participants watched the ∼1-minute videos) and no-stimulus baseline data from each recording block (6 trials) were concatenated, each feature was z-scored jointly across both, and the data were then split back into task and baseline sets. After block-wise z-score normalization, each feature matrix retained a block identifier that was used to keep all trials from a given recording block together during cross-validation, ensuring that data (both video-clip epochs and no-stimulus baseline) from the same block never appeared in both the training and test partitions. Instead of a standard grouped k-fold split (k = 4), we employed a custom block-grouped cross-validation scheme designed to guarantee that both classes were represented in every training and test fold. The procedure first identified all blocks containing at least one positive (class-1, emotion class) sample and verified that enough such blocks existed to populate the requested number of folds; when this was not the case, the number of folds was reduced accordingly, and emotions with too few positive-carrying blocks were skipped (e.g., 5/15 emotions were skipped for ppt4). One positive block was then assigned to each fold’s test set, and the remaining blocks were distributed across folds to balance fold sizes, so that each fold’s training and test sets contained both classes by construction. Within each fold, features were min-max scaled to [0, 1] using parameters fitted on the training data (scikit-learn^90^), and the majority class was subsampled five times without replacement to balance classes, yielding the training and test sets used for grid search and model fitting. A fixed random seed was used for reproducibility. Subsampling was performed separately for training and test sets to prevent data leakage.

### Classification

For each participant, separate binary classifiers were trained to distinguish task (all emotions combined together) vs baseline, and each emotion vs. the no-stimulus baseline or all other emotions combined (refer to the next paragraph). An emotion’s training set comprised the trials positively rated for that emotion together with all baseline segments. Three model families were used: logistic regression (saga solver; l1-ratio searched over linear five values in [0,1] and inverse elastic net regularization strength C over {0.5, 1, 5, 10}); a linear support vector machine (C over {0.5, 1, 5, 10}, with probability estimates); and a random forest (entropy criterion, out-of-bag scoring; 100 or 300 trees and maximum depth in {1, 3, 6, none}). Hyperparameters were selected by grid search within each training fold using an internal 2-fold cross-validation that optimized the area under the receiver operating characteristic curve (AUC). We included only emotions rated in more than 10 trials (∼100 augmented 10-s segments). This excluded five emotions (Calmness, Boredom, Satisfaction, Relief and Joy) for one participant (ppt 4) in the emotion-versus-baseline analyses.

The three classification schemes differed (task versus baseline, Fig. 2B; each emotion versus baseline, Fig. 2C–D and Supplementary Figure S2) only in how the positive and negative classes were defined prior to this shared fold-construction step. In the per-emotion analysis (i.e., each emotion versus baseline), folds were built separately for each emotion: task trials rated positive for that emotion were labeled as the positive class and combined with all baseline trials as the negative class, producing a distinct, class-balanced set of folds for each emotion-versus-baseline contrast. In the task-versus-baseline analysis, all task trials were instead pooled and labeled as the positive class against all baseline trials, and a single set of folds was constructed to capture the general task-versus-baseline distinction rather than any individual emotion. In the “each emotion versus all other” (i.e., one-vs-all) models, we used trials in which the target emotion was present as class 1 versus all remaining trials in which it was absent (though other emotions could be present). The block-grouping, train-only scaling, and balanced-subsampling and cross-validation steps were identical across both analyses; only the upstream label definition distinguished them.

Performance was summarized by AUC, and accuracy, and ROC curves were aggregated across folds to show cross-validation variability. To evaluate classification performance against chance, we constructed surrogate permutation models by shuffling class labels and training them exactly as the main emotion models (Fig. 2C, bottom panel and Supplementary Figure 2, bottom panel); model performance was compared with the confidence interval of these surrogates.

The data preparation, machine learning and feature clustering is visualized in Extended Data Figure 1.

### Feature attribution and stability

For interpretation, all clustering analyses were performed separately for each participant and each emotion, and were restricted to the features that the classifiers relied on consistently. For a given emotion, a feature was retained as “stable” only if it received a non-zero elastic-net weight in at least 90% of cross-validation runs. For each stable feature we recorded its mean signed weight, the standard error across runs, and its selection frequency, and ranked features by absolute mean weight. To remove the long tail of marginally contributing predictors, features were ordered by descending absolute importance and only those falling within the top 90% of the cumulative absolute-importance distribution were carried forward, leaving a compact, participant-and emotion-specific set of stable features (refer to Supplementary Figures S3-6).

### Feature clustering

To express each emotion’s signature at the level of physiology, we then asked, for each participant, how these stable features covaried across samples. The trials rated positive for that emotion were combined with all no-stimulus baseline trials, the resulting sample matrix was restricted to that emotion’s stable features, and a Pearson correlation matrix was computed between every pair of features across samples. This feature × feature matrix was submitted to agglomerative hierarchical clustering using Ward’s linkage on a Euclidean distance metric with optimal leaf ordering. Rather than fixing the number of clusters a priori, we swept candidate solutions from two clusters up to a maximum determined by the number of available features (the smaller of 15 and the number of stable features) and selected the partition maximizing the average silhouette score; the corresponding silhouette value and cluster count were retained for reporting. Results were displayed as symmetric correlation matrices overlaid with matched row and column dendrograms, yielding one cluster map per emotion per participant (Fig 3B).

The feature clustering findings were summarized in two complementary formats. First, for each emotion we generated a circular (polar) plot (Figure 3C and 3D, Figure 4 & Supplementary Figures S7-S9). Every stable feature occupies one angular sector of the ring. The radial extent of each sector encodes the feature’s mean absolute importance. The sector color denotes its assigned correlation cluster, and a legend maps each color to its cluster. Features are grouped around the ring by neurophysiological category (ground truth in black). Concentric dashed reference rings mark fixed fractions of each emotion’s maximum absolute importance; the innermost ring, at the first quartile of the importance range, serves as a visual threshold separating dominant from marginal contributors.

Second, features exceeding this first-ring cutoff were tabulated per emotion (Tables 1 & 2 & Supplementary Tables T3 and T4). Within each table, features were aggregated by cluster, and cluster labels differing only by an ordinal suffix were merged. For each emotion × cluster pair we reported two values: the number of contributing features, and the signed importance of the feature with the largest absolute magnitude in that cluster. These summaries were arranged with clusters as rows and emotions as columns. Rows were ordered by the largest absolute importance observed across emotions. Cells were shaded on a diverging color scale, such that the sign and strength of each cluster’s contribution can be read at a glance. Empty cells indicate clusters falling below the first-ring threshold.

### Behavioral clustering and the neurophysiological-behavioral consensus network

#### Emotion clustering

To assess whether different emotions relied on similar physiological features, we clustered emotions based on stable features’ rankings from the model training. For each emotion, its trimmed stable features were ordered by descending absolute importance and assigned integer ranks, producing a feature × emotion rank matrix; features absent from a given emotion’s stable set were assigned a rank one step beyond the maximum observed rank so that all emotions were defined over a common feature space. Spearman rank correlations were computed between every pair of emotion columns, and the resulting emotion × emotion similarity matrix was clustered using Ward’s linkage on a Euclidean metric and visualized as an annotated cluster map. Cluster memberships were extracted at four successive depths of the dendrogram (two through five clusters), allowing emotion groupings to be examined at multiple levels of granularity.

For the self-report grouping, the participant’s categorical ratings were binarized and clustered by their correlation structure. In more detail, for each participant, we took the trial × emotion matrix of ratings and computed the Pearson correlation between every pair of emotion columns across trials. This yields an emotion × emotion covariance structure describing which emotions tended to be endorsed together. Similar to the emotion clustering by features, this matrix was clustered with Ward’s linkage on a Euclidean metric and displayed as an annotated cluster map. Cluster memberships were extracted at four dendrogram depths (two through five clusters), matching the depths used for the neurophysiological clustering.

#### Consensus network

Emotion clustering analyses provided two independent partitions of the same emotions. One came from neurophysiology, based on how similarly the classifiers ranked neurophysiological features across emotions. The other came from self-reports, based on how similarly participants rated the emotions across trials. To compare them directly, we built an undirected graph using NetworkX^91^. Each emotion is a node. Every pair of emotions was then checked twice: once for whether they fell in the same neurophysiological cluster, and once for whether they fell in the same behavioral cluster. An edge was drawn whenever at least one of those two checks was positive.

Edges were color-coded and weighted by which modalities agreed. If both the neurophysiological and behavioral partitions placed a pair in the same cluster, the pair was joined by a thick green consensus edge. If only the neurophysiological partition grouped them, the edge was thinner and blue. If only the self-reports grouped them, the edge was thinner and red. Pairs that were separated in both partitions received no edge at all. Nodes were laid out one connected component at a time using a spring layout, with components spaced apart so that isolated groups of emotions remain visually distinct. The resulting figure makes the degree of cross-modal convergence readable.

### Common emotion clustering scheme across all participants

To search for consensus among participants on emotion clustering, for each participant, the emotion-level cluster assignments derived from the feature (i.e., neurophysiology)-ranking analysis were stored and then pooled. This gave one independent emotion partition per participant, all defined at the same dendrogram depth (four clusters). We then built a single undirected graph with NetworkX^91^. Similar to the previous section, each emotion is a node, and the node set is the union of all emotions present in any participant, so that emotions missing from one participant are still represented. Every pair of emotions was then evaluated across participants. For each participant, we simply asked whether the two emotions fell in the same cluster, and we counted how many participants agreed. Pairs were only compared within a participant if both emotions existed for that participant.

Edges were then color-coded by the level of agreement in a similar way mentioned above (See Consensus network). A pair grouped together in every participant received a thick green edge, marking full consensus. A pair grouped together in more than one (Fig. 6A) or two (Supplementary Figure S16A) participants, but not all, received a black edge, marking partial consensus. A pair grouped together in exactly one participant received a thinner edge colored by that participant’s identity, marking an idiosyncratic grouping. These cliques identify sets of emotions that were mutually co-clustered in the majority of participants, and they define the emotion groupings we report as reproducible among participants.

### Probing feature clusters based on emotion grouping (using Borda rose plots)

Each rose summarizes the features that most consistently drive the elastic-net models within an emotion group: within each emotion, the stable features are ranked by absolute elastic-net weight, and a feature receives consensus points equal to (total number of features − rank + 1), so that top-ranked features earn the most points and features absent from an emotion’s stable set score zero. Points are summed across all emotions belonging to that group, normalized by the maximum attainable score (100% = a feature ranked first for every member emotion), and plotted as a polar bar whose radius encodes this consensus score (0–100). Critically, the angular positions are fixed to a single global feature-cluster ordering shared by every rose, and each wedge is colored by its modality/feature cluster (EDA, TREV, heart rate, blood pressure, respiration, zygomaticus, corrugator, eye/blink, and EEG delta/theta/alpha/low-alpha/beta/gamma), so that roses can be compared directly by shape and color rather than read individually.

To test whether a feature’s consensus exceeds chance, the same score was recomputed from the label-permuted (surrogate) elastic-net models to form a per-feature null distribution: the dashed ring marks the null 95th percentile, open stars denote features significant after Benjamini–Hochberg FDR correction (q < 0.05), and filled stars denote features that additionally survive max-T family-wise error correction (p < 0.05). Because correction was applied across all drawn wedges within a participant, and because single-emotion groups yield discrete, frequently tied Borda scores, a wedge can exceed its null 95th percentile (an uncorrected per-wedge reference) without reaching FDR significance.

#### Cross-participant Borda consensus

Briefly, each participant (except ppt 4) contributes one cluster × emotion weight table, and all tables are first reindexed onto the union of feature clusters and emotions so that every participant is scored on the same scale. Akin to what is mentioned above, within each participant, feature clusters are ranked by absolute weight per emotion and awarded Borda points equal to (total clusters − rank + 1), with absent clusters scoring zero. A participant then “votes” for a feature cluster if it earns any points in their data, and clusters reaching a strict majority of votes (≥2 of 3 participants) are retained as the consensus set, ordered by their pooled points; minority clusters are still drawn, appended afterwards, so they appear as short or absent bars and the full union remains visible. Points are summed across participants and across the group’s member emotions and normalized by the maximum attainable score (clusters × emotions × participants), so that bar radius expresses the percentage of the maximum possible cross-participant consensus (100% = ranked first for every emotion in every participant).

### Statistics

Reported associations between physiological measures (e.g. EEG band power and tonic EDA) used Spearman rank correlation with two-sided P values. Cross-validation provided the principal control against overfitting; all performance metrics and feature attributions were computed on held-out blocks and aggregated across folds. Emotion vs. baseline models’ performance was compared with the confidence interval of the label-permuted surrogate models (Supplementary Table T2 and star values on Supplementary figure 2A and Figure 2C). Feature extraction, classification and all clustering analyses were implemented in Python using scikit-learn^90^, SciPy, NumPy, pandas, and NetworkX.

#### Statistical assessment of the Borda consensus rose plots

To determine whether the physiological clusters highlighted in the Nightingale rose plots reflect genuine, above-chance consensus rather than artifacts of the feature-selection pipeline, we constructed a label-permutation null distribution for every plotted wedge. For each of N (e.g., 100) permutations, we shuffled the trial labels prior to Elastic Net (EN) fitting and passed the resulting per-fold coefficients through the identical downstream pipeline used to generate the observed figure: a stability filter (feature retained if non-zero in ≥90% of cross-validation folds), a cumulative-importance plateau trim (retaining features up to 90% of total |importance|), a ring-1 magnitude cutoff (|importance| ≥ 25% of that emotion’s maximum), collapse to physiological feature clusters, and Borda rank-aggregation into a normalized (0-100) consensus score per emotion-group. This yielded, for every cluster × emotion-group wedge, a null distribution of 100 surrogate consensus scores obtained under the identical feature clustering and processing pipeline as the observed data. We computed an exact one-sided empirical p-value for each wedge as p = (1 + Σ[S(null) ≥ S(obs)]) / (N of perm + 1), a construction that avoids a p-value of exactly zero even at the extreme of the null distribution ^92^. Because feature to cluster assignment (a fixed mapping derived from the observed analysis, with any unmapped features selected solely under the null aggregated into fallback pseudo-clusters so they still compete in the Borda ranking without inflating null scores) and emotion groupings were common to both the observed and permuted pipelines, wedge scores under the null and the alternative were directly comparable. To correct for testing multiple wedges within a participant, we applied the Benjamini-Hochberg procedure (FDR < 0.05) across all drawn wedges; a more conservative Westfall-Young max-T correction (comparing each wedge to the null distribution of the single largest wedge score per permutation within its emotion group) controlled the family-wise error rate as a secondary, assumption-free check. Wedges surviving FDR correction are marked with an open star in the annotated rose plots; wedges additionally surviving the max-T correction are marked with a filled star. Because correction was applied across all drawn wedges within a participant, and because single-emotion groups yield discrete, frequently tied Borda scores, a wedge can exceed its null 95th percentile (an uncorrected per-wedge reference) without reaching FDR significance, such as oculomotor wedge in ppt2’s Calmness. (Fig. 5D)

#### Statistics for cross-participant rose plots

To test whether participants agree on which feature clusters characterize each consensus emotion group, we used a within-participant rank-shuffle permutation test on the cross-participant Borda scores. For each participant, every emotion’s feature clusters were ranked by absolute elastic-net weight and assigned Borda points (number of clusters − rank + 1; absent clusters scored zero). A cluster’s observed group score was the sum of its points across the group’s member emotions and across participants, normalized by the maximum attainable score. The null distribution was built by randomly reassigning each participant’s Borda points among that participant’s own clusters, independently for every emotion, and recomputing the group score (5,000 permutations). This keeps each participant’s set of scores and cluster vocabulary intact but breaks any correspondence between clusters across participants and emotions. A cluster’s score therefore exceeds the null only if participants consistently rank the same cluster highly for the group’s emotions. One-sided empirical p-values were computed as (1 + number of null scores ≥ observed) / (1 + number of permutations).

Multiple comparisons were corrected within each emotion group. The test family was restricted, before testing, to clusters present in at least two participants’ feature sets. A cluster found in only one participant cannot show cross-participant agreement, and because each participant’s cluster set is fixed under the permutation, this restriction does not depend on the observed scores. Within this family we applied Benjamini–Hochberg FDR correction (open stars, q < 0.05). We also applied a stricter family-wise correction using a studentized max-T procedure, in which each cluster’s score was standardized against its own null before taking the maximum across clusters (filled stars, p < 0.05). This prevents clusters that score highly for every emotion from dominating the maximum. In the rose plots, the dashed arc on each wedge marks the null 95th percentile.

Open circles mark wedges within the test family exceeding this threshold (uncorrected p < 0.05) that did not survive correction, and these are reported as exploratory. Note that in these statistical tests a cluster present in only one participant will never get an open circle, even if its bar visually clears the dashed 95th-percentile arc on the plot

## Data availability

The multimodal biosensing dataset that supports the findings of this study is available from corresponding authors upon reasonable request, subject to participant-privacy and institutional constraints.

## Code availability

The software package for the methods in this paper and example code is available at https://github.com/abbasilab/Emosense.

## Acknowledgements

We thank the members of Neuroscape and the Abbasi-Asl laboratory at the University of California, San Francisco, Joseph Chen, for technical assistance and discussion, and the study participants for their time. The authors gratefully acknowledge support from the Patrick J. McGovern Foundation, UCSF Innovation Ventures, and the UCSF Weill Institute for Neurosciences (UCSF Weill Catalyst Award).

## Author Contributions

RA-A and AG conceived the study. RA-A, AG, TZ, and DZ designed the data collection protocol with contributions from BN, JT, and RC. BK, RAS, and RA-A developed the stimulus paradigm. BN, RC, VW, BK, IH assisted with subject recruitment and data collection. MB conceptualized the analytical framework, performed all analysis and modeling with contributions from RA-A. MB, RA-A, AG, and TZ interpreted the findings. MO contributed to data alignment under MB’s supervision. MB wrote the manuscript with contributions from RA-A, AG, and TZ. All authors reviewed and revised the manuscript.

---

**Extended Data Figure 1.**
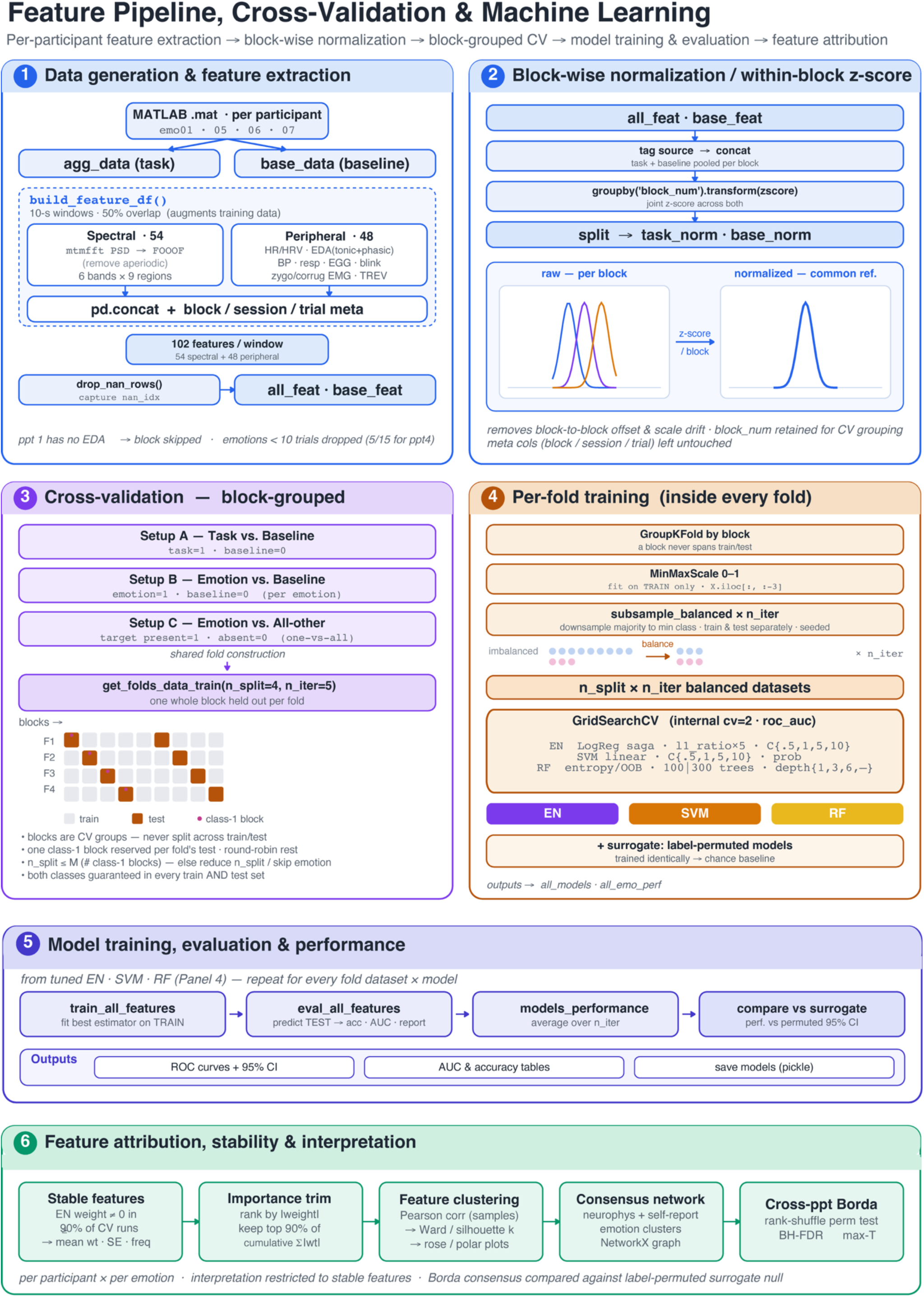
Demonstrates Machine Learning pipeline.

**Extended Data Table 1.**
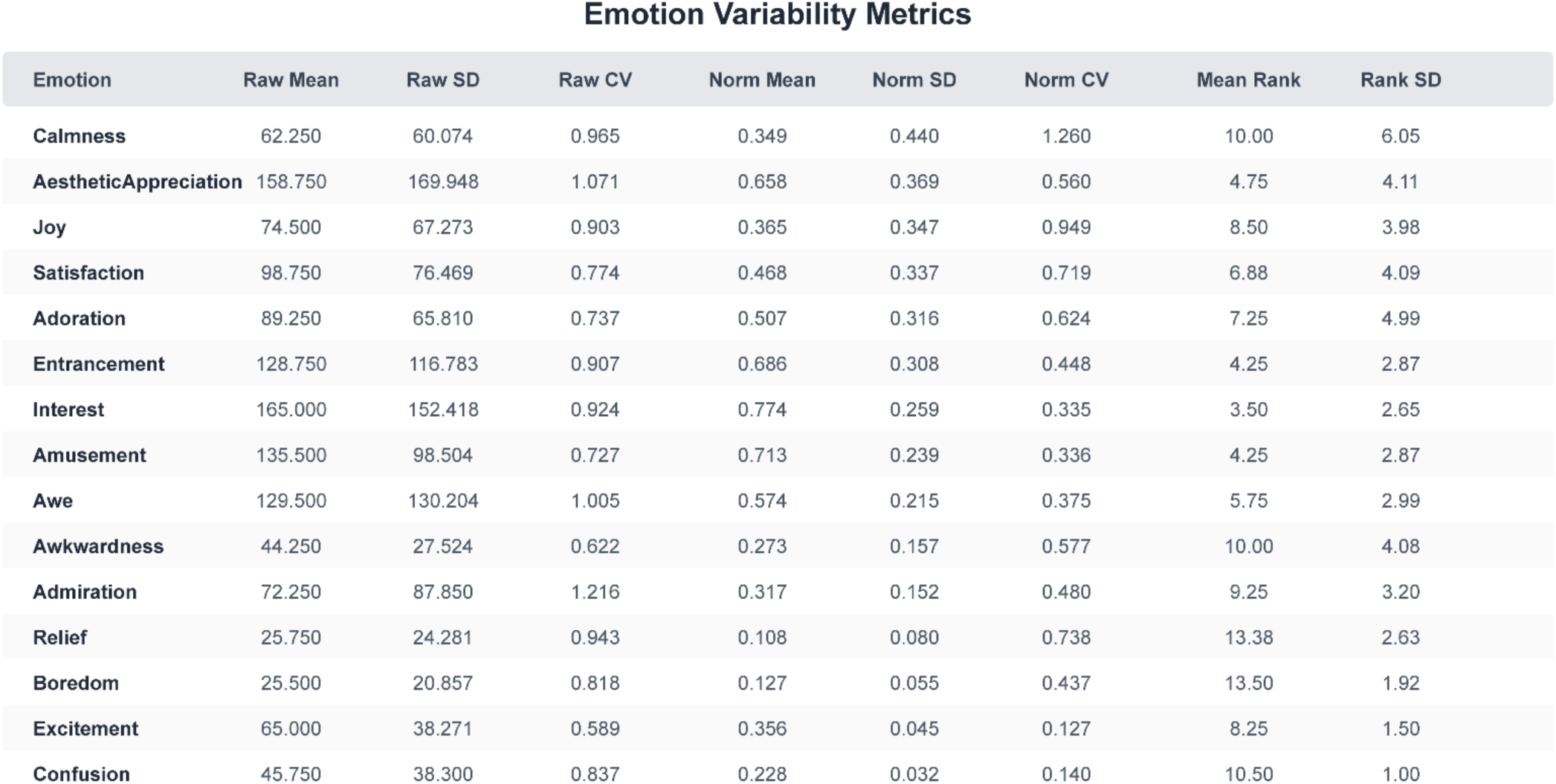
Variability of emotion ratings across participants. For each emotion, the mean, s.d. and coefficient of variation (CV) of raw and normalized counts across participants, and the mean and s.d. of within-participant ranks (1 = most frequent). Rank s.d. ranged from 6.05 (Calmness) to 1.00 (Confusion); agreement in rank order was moderate (Kendall’s W = 0.504).

Emotion Variability Metrics
| Emotion | Raw Mean | Raw SD | Raw CV | Norm Mean | Norm SD | Norm CV | Mean Rank | Rank SD |
| --- | --- | --- | --- | --- | --- | --- | --- | --- |
| Calmness | 62.250 | 60.074 | 0.965 | 0.349 | 0.440 | 1.260 | 10.00 | 6.05 |
| AestheticAppreciation | 158.750 | 169.948 | 1.071 | 0.658 | 0.369 | 0.560 | 4.75 | 4.11 |
| Joy | 74.500 | 67.273 | 0.903 | 0.365 | 0.347 | 0.949 | 8.50 | 3.98 |
| Satisfaction | 98.750 | 76.469 | 0.774 | 0.468 | 0.337 | 0.719 | 6.88 | 4.09 |
| Adoration | 89.250 | 65.810 | 0.737 | 0.507 | 0.316 | 0.624 | 7.25 | 4.99 |
| Entrancement | 128.750 | 116.783 | 0.907 | 0.686 | 0.308 | 0.448 | 4.25 | 2.87 |
| Interest | 165.000 | 152.418 | 0.924 | 0.774 | 0.259 | 0.335 | 3.50 | 2.65 |
| Amusement | 135.500 | 98.504 | 0.727 | 0.713 | 0.239 | 0.336 | 4.25 | 2.87 |
| Awe | 129.500 | 130.204 | 1.005 | 0.574 | 0.215 | 0.375 | 5.75 | 2.99 |
| Awkwardness | 44.250 | 27.524 | 0.622 | 0.273 | 0.157 | 0.577 | 10.00 | 4.08 |
| Admiration | 72.250 | 87.850 | 1.216 | 0.317 | 0.152 | 0.480 | 9.25 | 3.20 |
| Relief | 25.750 | 24.281 | 0.943 | 0.108 | 0.080 | 0.738 | 13.38 | 2.63 |
| Boredom | 25.500 | 20.857 | 0.818 | 0.127 | 0.055 | 0.437 | 13.50 | 1.92 |
| Excitement | 65.000 | 38.271 | 0.589 | 0.356 | 0.045 | 0.127 | 8.25 | 1.50 |
| Confusion | 45.750 | 38.300 | 0.837 | 0.228 | 0.032 | 0.140 | 10.50 | 1.00 |

## References

1. Kreibig, S. D. Autonomic nervous system activity in emotion: A review. Biol. Psychol. 84, 394–421 (2010).

2. Levenson, R. W. The Autonomic Nervous System and Emotion. Emot. Rev. 6, 100–112 (2014).

3. Kluger, D. S., Allen, M. G. & Gross, J. Brain–body states embody complex temporal dynamics. Trends Cogn. Sci. 28, 695–698 (2024).

4. Uddin, L. Q., Kinnison, J., Pessoa, L. & Anderson, M. L. Beyond the Tripartite Cognition–Emotion– Interoception Model of the Human Insular Cortex. J. Cogn. Neurosci. 26, 16–27 (2014).

5. Craig, A. D. How do you feel? Interoception: the sense of the physiological condition of the body. Nat. Rev. Neurosci. 3, 655–666 (2002).

6. Insel, T. et al. Research domain criteria (RDoC): toward a new classification framework for research on mental disorders. Am. J. Psychiatry 167, 748–751 (2010).

7. Wright, A. G. C. & Woods, W. C. Personalized models of psychopathology. Annu Rev Clin Psychol 16, 49– 74 (2020).

8. Siegel, E. H. et al. Emotion fingerprints or emotion populations? A meta-analytic investigation of autonomic features of emotion categories. Psychol. Bull. 144, 343–393 (2018).

9. Hoemann, K. Context-aware experience sampling reveals the scale of variation in affective experience. Sci Rep 10, 12459 (2020).

10. Fisher, A. J., Medaglia, J. D. & Jeronimus, B. F. Lack of group-to-individual generalizability is a threat to human subjects research. Proc. Natl. Acad. Sci. 115, E6106–E6115 (2018).

11. Molenaar, P. C. M. & Campbell, C. G. The New Person-Specific Paradigm in Psychology. Curr. Dir. Psychol. Sci. 18, 112–117 (2009).

12. Gordon, E. M. et al. Precision Functional Mapping of Individual Human Brains. Neuron 95, 791–807.e7 (2017).

13. Gratton, C. Functional brain networks are dominated by stable group and individual factors, not cognitive or daily variation. Neuron 98, 439–452 (2018).

14. Smith, P. L. & Little, D. R. Small is beautiful: in defense of the small-N design. Psychon Bull Rev 25, 2083–2101 (2018).

15. Behnke, M., Kreibig, S. D., Kaczmarek, L. D., Assink, M. & Gross, J. J. Autonomic nervous system activity during positive emotions: A meta-analytic review. Emot. Rev. 14, 132–160 (2022).

16. Stephens, C. L., Christie, I. C. & Friedman, B. H. Autonomic specificity of basic emotions: Evidence from pattern classification and cluster analysis. Biol. Psychol. 84, 463–473 (2010).

17. Kragel, P. A. & Labar, K. S. Multivariate pattern classification reveals autonomic and experiential representations of discrete emotions. Emotion 13, 681–690 (2013).

18. Kragel, P. A. & LaBar, K. S. Decoding the Nature of Emotion in the Brain. Trends Cogn. Sci. 20, 444–455 (2016).

19. Saarimäki, H. et al. Discrete Neural Signatures of Basic Emotions. Cereb. Cortex 26, 2563–2573 (2016).

20. Cowen, A. S. & Keltner, D. Self-report captures 27 distinct categories of emotion bridged by continuous gradients. Proc. Natl. Acad. Sci. 114, E7900–E7909 (2017).

21. Horikawa, T., Cowen, A. S., Keltner, D. & Kamitani, Y. The Neural Representation of Visually Evoked Emotion Is High-Dimensional, Categorical, and Distributed across Transmodal Brain Regions. iScience 23, 101060 (2020).

22. Cowen, A. S. & Keltner, D. Semantic space theory: a computational approach to emotion. Trends Cogn Sci 25, 124–136 (2021).

23. Hu, X. et al. EEG Correlates of Ten Positive Emotions. Front. Hum. Neurosci. 11, 26 (2017).

24. Bijanzadeh, M. et al. Decoding naturalistic affective behaviour from spectro-spatial features in multiday human iEEG. Nat. Hum. Behav. 6, 823–836 (2022).

25. Koelstra, S. et al. DEAP: A Database for Emotion Analysis Using Physiological Signals. IEEE Trans. Affect. Comput. 3, 18–31 (2012).

26. Saganowski, S. et al. Emognition dataset: emotion recognition with self-reports, facial expressions, and physiology using wearables. Sci. Data 9, 158 (2022).

27. Dawson, M. E., Schell, A. M. & Filion, D. L. The electrodermal system. in Handbook of psychophysiology, 3rd ed 159–181 (Cambridge University Press, New York, NY, US, 2007). doi:10.1017/CBO9780511546396.007.

28. Klimesch, W. EEG alpha and theta oscillations reflect cognitive and memory performance: a review and analysis. Brain Res. Rev. 29, 169–195 (1999).

29. Klimesch, W. Alpha-band oscillations, attention, and controlled access to stored information. Trends Cogn. Sci. 16, 606–617 (2012).

30. Barrett, L. F. & Russell, J. A. The Structure of Current Affect: Controversies and Emerging Consensus. Curr. Dir. Psychol. Sci. 8, 10–14 (1999).

31. Siedlecka, E. & Denson, T. F. Experimental Methods for Inducing Basic Emotions: A Qualitative Review. Emot. Rev. 11, 87–97 (2019).

32. Drysdale, A. T. et al. Resting-state connectivity biomarkers define neurophysiological subtypes of depression. Nat. Med. 23, 28–38 (2017).

33. Fried, E. I. & Nesse, R. M. Depression is not a consistent syndrome: an investigation of unique symptom patterns in the STAR*D study. J. Affect. Disord. 172, 96–102 (2015).

34. Tozzi, L. et al. Personalized brain circuit scores identify clinically distinct biotypes in depression and anxiety. Nat. Med. 30, 2076–2087 (2024).

35. Sani, O. G. et al. Mood variations decoded from multi-site intracranial human brain activity. Nat. Biotechnol. 36, 954–961 (2018).

36. Kirkby, L. A. et al. An Amygdala-Hippocampus Subnetwork that Encodes Variation in Human Mood. Cell 175, 1688–1700.e14 (2018).

37. Bijanki, K. R. et al. Cingulum stimulation enhances positive affect and anxiolysis to facilitate awake craniotomy. J. Clin. Invest. 129, 1152–1166 (2019).

38. Scangos, K. W. et al. Closed-loop neuromodulation in an individual with treatment-resistant depression. Nat. Med. 27, 1696–1700 (2021).

39. Rao, V. R. et al. Direct Electrical Stimulation of Lateral Orbitofrontal Cortex Acutely Improves Mood in Individuals with Symptoms of Depression. Curr. Biol. 28, 3893–3902.e4 (2018).

40. Metzger, B. A. et al. Intracranial stimulation and EEG feature analysis reveal affective salience network specialization. Brain 146, 4366–4377 (2023).

41. Alagapan, S. Cingulate dynamics track depression recovery with deep brain stimulation. Nature 622, 130– 138 (2023).

42. Zou, H. & Hastie, T. Regularization and variable selection via the elastic net. J. R. Stat. Soc. Ser. B Stat. Methodol. 67, 301–320 (2005).

43. Ochsner, K. & Gross, J. The cognitive control of emotion. Trends Cogn. Sci. 9, 242–249 (2005).

44. Gross, J. J. Emotion regulation: Current status and future prospects. Psychol. Inq. 26, 1–26 (2015).

45. Nummenmaa, L., Glerean, E., Hari, R. & Hietanen, J. K. Bodily maps of emotions. Proc. Natl. Acad. Sci. 111, 646–651 (2014).

46. Azari, B. Comparing supervised and unsupervised approaches to emotion categorization in the human brain, body, and subjective experience. Sci Rep 10, 20284 (2020).

47. Ekman, P., Levenson, R. W. & Friesen, W. V. Autonomic nervous system activity distinguishes among emotions. Science 221, 1208–1210 (1983).

48. Ekman, P. & Cordaro, D. What is Meant by Calling Emotions Basic. Emot. Rev. 3, 364–370 (2011).

49. Barrett, L. F. The theory of constructed emotion: an active inference account of interoception and categorization. Soc. Cogn. Affect. Neurosci. 12, 1–23 (2017).

50. Russell, J. A. A circumplex model of affect. J. Pers. Soc. Psychol. 39, 1161–1178 (1980).

51. Shiota, M. N., Neufeld, S. L., Yeung, W. H., Moser, S. E. & Perea, E. F. Feeling good: autonomic nervous system responding in five positive emotions. Emotion 11, 1368–1378 (2011).

52. Haufe, S. On the interpretation of weight vectors of linear models in multivariate neuroimaging. NeuroImage 87, 96–110 (2014).

53. Keltner, D., Brooks, J. A. & Cowen, A. Semantic space theory: data-driven insights into basic emotions. Curr Dir Psychol Sci 32, 242–249 (2023).

54. Koide-Majima, N., Nakai, T. & Nishimoto, S. Distinct dimensions of emotion in the human brain and their representation on the cortical surface. NeuroImage 222, 117258 (2020).

55. Westlin, C., Singh, A., Erdogmus, D., Stratis, G. & Barrett, L. F. Exploring theory-laden observations in the brain basis of emotional experience. https://arxiv.org/abs/2507.00320 (2025).

56. Chen, J. A large finer-grained affective computing EEG dataset. Sci Data 10, 740 (2023).

57. Gable, P. & Harmon-Jones, E. The motivational dimensional model of affect: implications for breadth of attention, memory, and cognitive categorisation. Cogn Emot 24, 322–337 (2010).

58. Vessel, E. A., Starr, G. G. & Rubin, N. Art reaches within: aesthetic experience, the self and the default mode network. Front. Neurosci. 7, 258 (2013).

59. Vessel, E. A., Starr, G. G. & Rubin, N. The brain on art: intense aesthetic experience activates the default mode network. Front. Hum. Neurosci. 6, 66 (2012).

60. van Elk, M., Arciniegas Gomez, M. A., van der Zwaag, W., van Schie, H. T. & Sauter, D. The neural correlates of the awe experience: reduced default mode network activity during feelings of awe. Hum Brain Mapp 40, 3561–3574 (2019).

61. Nakano, T., Yamamoto, Y., Kitajo, K., Takahashi, T. & Kitazawa, S. Synchronization of spontaneous eyeblinks while viewing video stories. Proc. R. Soc. B Biol. Sci. 276, 3635–3644 (2009).

62. Campos, B., Shiota, M. N., Keltner, D., Gonzaga, G. C. & Goetz, J. L. What is shared, what is different? Core relational themes and expressive displays of eight positive emotions. Cogn. Emot. 27, 37–52 (2013).

63. Haegens, S., Cousijn, H., Wallis, G., Harrison, P. J. & Nobre, A. C. Inter-and intra-individual variability in alpha peak frequency. NeuroImage 92, 46–55 (2014).

64. Aftanas, L. I., Reva, N. V., Varlamov, A. A., Pavlov, S. V. & Makhnev, V. P. Analysis of evoked EEG synchronization and desynchronization in conditions of emotional activation in humans: temporal and topographic characteristics. Neurosci. Behav. Physiol. 34, 859–867 (2004).

65. Keil, A. et al. Effects of emotional arousal in the cerebral hemispheres: a study of oscillatory brain activity and event-related potentials. Clin. Neurophysiol. 112, 2057–2068 (2001).

66. Jung, J. et al. Intracerebral γ modulations reveal interaction between emotional processing and action outcome evaluation in the human orbitofrontal cortex. Int. J. Psychophysiol. Off. J. Int. Organ. Psychophysiol. 79, 64–72 (2011).

67. Zheng, W.-L. & Lu, B.-L. Investigating critical frequency bands and channels for EEG-based emotion recognition with deep neural networks. IEEE Trans Auton Ment Dev 7, 162–175 (2015).

68. Eastwood, J. D., Frischen, A., Fenske, M. J. & Smilek, D. The unengaged mind: defining boredom in terms of attention. Perspect Psychol Sci 7, 482–495 (2012).

69. Danckert, J. & Merrifield, C. Boredom, sustained attention and the default mode network. Exp Brain Res 236, 2507–2518 (2018).

70. Compton, R. J., Gearinger, D. & Wild, H. The wandering mind oscillates: EEG alpha power is enhanced during moments of mind-wandering. Cogn. Affect. Behav. Neurosci. 19, 1184–1191 (2019).

71. Muthukumaraswamy, S. High-frequency brain activity and muscle artifacts in MEG/EEG: A review and recommendations. Front. Hum. Neurosci. 7, 138 (2013).

72. Whitham, E. M. Scalp electrical recording during paralysis: quantitative evidence that EEG frequencies above 20 Hz are contaminated by EMG. Clin Neurophysiol 118, 1877–1888 (2007).

73. Yuval-Greenberg, S., Tomer, O., Keren, A. S., Nelken, I. & Deouell, L. Y. Transient induced gamma-band response in EEG as a manifestation of miniature saccades. Neuron 58, 429–441 (2008).

74. Hipp, J. F. & Siegel, M. Dissociating neuronal gamma-band activity from cranial and ocular muscle activity in EEG. Front Hum Neurosci 7, 338 (2013).

75. Donoghue, T. et al. Parameterizing neural power spectra into periodic and aperiodic components. Nat. Neurosci. 23, 1655–1665 (2020).

76. Guillory, S. A. & Bujarski, K. A. Exploring emotions using invasive methods: review of 60 years of human intracranial electrophysiology. Soc. Cogn. Affect. Neurosci. 9, 1880–1889 (2014).

77. Davidson, R. J. Affective Style and Affective Disorders: Perspectives from Affective Neuroscience. Cogn. Emot. 12, 307–330 (1998).

78. Pasquini, L. et al. Dynamic autonomic nervous system states arise during emotions and manifest in basal physiology. Psychophysiology 60, e14218 (2023).

79. Lakatos, P., Karmos, G., Mehta, A. D., Ulbert, I. & Schroeder, C. E. Entrainment of neuronal oscillations as a mechanism of attentional selection. Science 320, 110–113 (2008).

80. Hasson, U., Malach, R. & Heeger, D. J. Reliability of cortical activity during natural stimulation. Trends Cogn. Sci. 14, 40–48 (2010).

81. Bradley, M. M. Natural selective attention: orienting and emotion. Psychophysiology 46, 1–11 (2009).

82. Kragel, P. A., Reddan, M. C., LaBar, K. S. & Wager, T. D. Emotion schemas are embedded in the human visual system. Sci. Adv. eaaw4358 (2019).

83. Kuppens, P. et al. Emotional inertia prospectively predicts the onset of depressive disorder in adolescence. Emotion 12, 283–289 (2012).

84. Delorme, A. & Makeig, S. EEGLAB: an open source toolbox for analysis of single-trial EEG dynamics including independent component analysis. J. Neurosci. Methods 134, 9–21 (2004).

85. Oostenveld, R., Fries, P., Maris, E. & Schoffelen, J.-M. FieldTrip: Open source software for advanced analysis of MEG, EEG, and invasive electrophysiological data. Comput. Intell. Neurosci. 2011, 156869 (2011).

86. Pan, J. & Tompkins, W. J. A real-time QRS detection algorithm. IEEE Trans. Biomed. Eng. 32, 230–236 (1985).

87. Makeig, S., Bell, A., Jung, T.-P. & Sejnowski, T. Independent Component Analysis of Electroencephalographic Data. in Advances in Neural Information Processing Systems vol. 8 (MIT Press, 1995).

88. Pan, T.-F. et al. When more is not better: Predicting human emotional states with brain and bodily biosensors. Biomed. Signal Process. Control 113, 109038 (2026).

89. Greco, A., Valenza, G., Lanata, A., Scilingo, E. P. & Citi, L. cvxEDA: A Convex Optimization Approach to Electrodermal Activity Processing. IEEE Trans. Biomed. Eng. 63, 797–804 (2016).

90. Pedregosa, F. Scikit-learn: machine learning in Python. J Mach Learn Res 12, 2825–2830 (2011).

91. Hagberg, A. A., Schult, D. A. & Swart, P. J. Exploring network structure, dynamics, and function using NetworkX. in Proc. 7th Python in Science Conference (SciPy2008) 11–15 (2008).

92. Phipson, B. & Smyth, G. K. Permutation p-values should never be zero: calculating exact p-values when permutations are randomly drawn. Stat. Appl. Genet. Mol. Biol. 9, (2010).

