## Supplemental Information for "Individualized brain-body signatures of positive emotions revealed by large-scale multimodal neurophysiological recording"

**This PDF file includes:**

Supplementary text
Figures S1 to S16
Supplementary Tables T1 to T5
SI References 5
Legends for Extended Figure and Table

### Supplementary findings

#### One-vs-all classification

To test whether emotions could also be distinguished from one another, we trained one-vs-all LR-EN classifiers that contrasted trials in which the target emotion was rated (class 1) with all remaining trials, regardless of other emotions rated on those trials (Methods). Performance was markedly lower and more variable than for the emotion-versus-baseline models (Fig. S2A): mean accuracies ranged from ~0.42 to ~0.73, with the highest values for Amusement in ppt 1 and Adoration and Joy in ppt 2. Accuracy exceeded the 97.5th percentile of label-permuted surrogate models (Fig. S2B) for 30 of 55 models (11/15, 8/15, 6/15 and 5/10 emotions in ppt 1-4). This weaker separability is expected from the task design: because participants could endorse several emotions per trial, the negative class contained trials that shared physiology with the target emotion. Feature interpretation in the main text was therefore based on the emotion-versus-baseline models.

#### Comparison with label-permuted surrogate models

Each emotion-versus-baseline model was compared with surrogate models trained with shuffled class labels but otherwise identical block-grouped cross-validation, class balancing and hyperparameter search (Methods), a standard test of classifier performance against chance[1,2]. A model was considered significant when its mean cross-validated accuracy exceeded the upper bound of the surrogates' 95% interval (2.5th–97.5th percentiles), a one-sided criterion at $\alpha = 0.025$. Surrogate accuracies were centered on chance, with intervals spanning ~0.41–0.59 across emotions and participants. All 55 emotion models exceeded this bound (Supplementary Table T2), with accuracies of 0.946–0.998 in ppt 1, 0.927–0.980 in ppt 2, 0.838–0.902 in ppt 3 and 0.850–0.909 in ppt 4.

#### Cross-participant emotion groupings including participant 4

Including participant 4, who contributed only 10 emotions, reduced the node set and the number of comparable emotion pairs but preserved the core structure of Fig. 6 (Fig. S16). Only Interest-Awe reached full consensus across all four participants, whereas Interest-Awe-Satisfaction-Amusement and Entrancement-Aesthetic Appreciation reached partial consensus in three of four participants (Fig. S16 A). Cross-participant rose plots for these groups were computed and tested as in Fig. 6B, using 5,000 within-participant permutations with Benjamini–Hochberg FDR and studentized max-T corrections within each group[3,4] (S3, S4). No cluster survived correction (Fig. S16 B). Several wedges reached only nominal significance (uncorrected $p < 0.05$), including posterior delta, posterior alpha and EDA for Entrancement-Aesthetic Appreciation and frontal gamma for Interest-Awe. These overlap with the dominant clusters of the three-participant analysis but are weaker, as expected when consensus must also include a participant with a reduced emotion set.

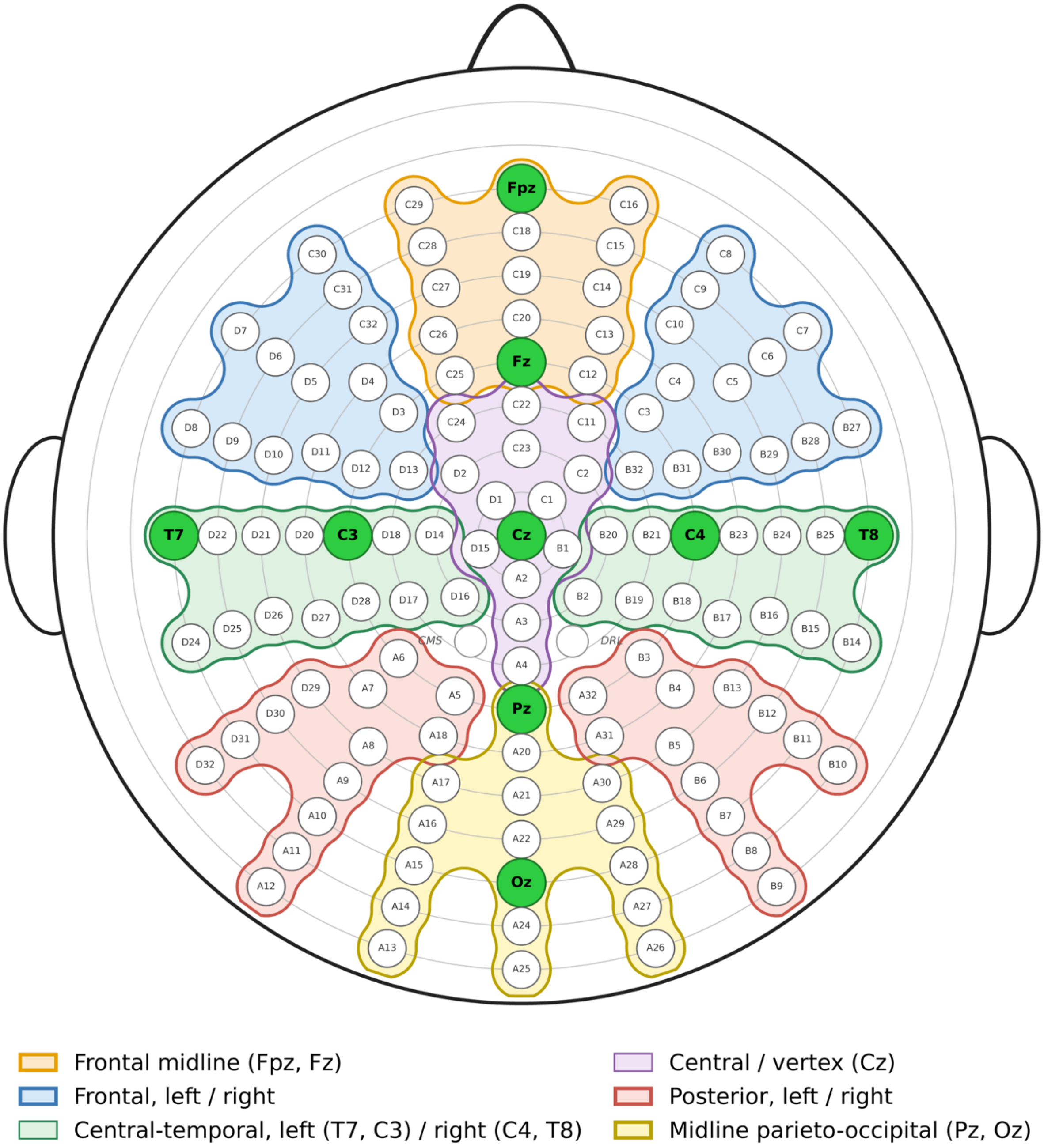


**Fig. S1.** BioSemi 128-channel montage and the regional grouping used for spectral features. Electrodes were grouped into a 3 × 3 grid (frontal, central–temporal and posterior rows × left, midline and right columns); band power was averaged within each region (Supplementary Table T1).

Supplementary Table T1. **All Neurophysiological features,** all features were computed within 10-second segments (50% overlap) of each 1-minute trial. EEG band power: multitaper spectra (FieldTrip mtmfft, ±0.5 Hz spectral smoothing, 1–51 Hz); the aperiodic (fractal) component was estimated with FOOOF and removed by taking the log of the ratio of the original spectrum to its aperiodic fit. Band features are the mean log oscillatory power within each band, averaged across the good electrodes of each of the nine montage regions (9 regions × 6 bands = 54 features).

EDA components were obtained with cvxEDA after z-scoring and linear detrending. Heart rate was derived from Pan–Tompkins R-peak detection. EMG envelopes (corrugator supercilii, zygomaticus major) were detrended, band-passed (10–200 Hz), rectified, smoothed, and normalized to the trial maximum; segments with SD > 5× the median SD were excluded.

SD = standard deviation; SCL/SCR = skin conductance level/response; TREV features are summarized as contractility and cardiac output index.

| No. | Modality | Feature | Description |
|---|---|---|---|
| 1 | **EEG** | Delta power left frontal | Log oscillatory (FOOOF-corrected) multitaper power in 1–4 Hz, averaged across electrodes of the region |
| 2 | | Theta power left frontal | Log oscillatory (FOOOF-corrected) multitaper power in 4–8 Hz, averaged across electrodes of the region |
| 3 | | Low alpha power left frontal | Log oscillatory (FOOOF-corrected) multitaper power in 8–10 Hz, averaged across electrodes of the region |
| 4 | | High alpha power left frontal | Log oscillatory (FOOOF-corrected) multitaper power in 10–12 Hz, averaged across electrodes of the region |
| 5 | | Beta power left frontal | Log oscillatory (FOOOF-corrected) multitaper power in 13–30 Hz, averaged across electrodes of the region |
| 6 | | Gamma power left frontal | Log oscillatory (FOOOF-corrected) multitaper power in 30–50 Hz, averaged across electrodes of the region |
| 7 | | Delta power frontal midline (Fpz, Fz) | Log oscillatory (FOOOF-corrected) multitaper power in 1–4 Hz, averaged across electrodes of the region |
| 8 | | Theta power frontal midline (Fpz, Fz) | Log oscillatory (FOOOF-corrected) multitaper power in 4–8 Hz, averaged across electrodes of the region |
| 9 | | Low alpha power frontal midline (Fpz, Fz) | Log oscillatory (FOOOF-corrected) multitaper power in 8–10 Hz, averaged across electrodes of the region |
| 10 | | High alpha power frontal midline (Fpz, Fz) | Log oscillatory (FOOOF-corrected) multitaper power in 10–12 Hz, averaged across electrodes of the region |
| 11 | | Beta power frontal midline (Fpz, Fz) | Log oscillatory (FOOOF-corrected) multitaper power in 13–30 Hz, averaged across electrodes of the region |
| 12 | | Gamma power frontal midline (Fpz, Fz) | Log oscillatory (FOOOF-corrected) multitaper power in 30–50 Hz, averaged across electrodes of the region |
| 13 | | Delta power right frontal | Log oscillatory (FOOOF-corrected) multitaper power in 1–4 Hz, averaged across electrodes of the region |
| 14 | | Theta power right frontal | Log oscillatory (FOOOF-corrected) multitaper power in 4–8 Hz, averaged across electrodes of the region |
| 15 | | Low alpha power right frontal | Log oscillatory (FOOOF-corrected) multitaper power in 8–10 Hz, averaged across electrodes of the region |
| 16 | | High alpha power right frontal | Log oscillatory (FOOOF-corrected) multitaper power in 10–12 Hz, averaged across electrodes of the region |
| 17 | | Beta power right frontal | Log oscillatory (FOOOF-corrected) multitaper power in 13–30 Hz, averaged across electrodes of the region |
| 18 | | Gamma power right frontal | Log oscillatory (FOOOF-corrected) multitaper power in 30–50 Hz, averaged across electrodes of the region |

| No. | Modality | Feature | Description |
|---|---|---|---|
| 19 |  | Delta power left central–temporal (T7, C3) | Log oscillatory (FOOOF-corrected) multitaper power in 1–4 Hz, averaged across electrodes of the region |
| 20 |  | Theta power left central–temporal (T7, C3) | Log oscillatory (FOOOF-corrected) multitaper power in 4–8 Hz, averaged across electrodes of the region |
| 21 |  | Low alpha power left central–temporal (T7, C3) | Log oscillatory (FOOOF-corrected) multitaper power in 8–10 Hz, averaged across electrodes of the region |
| 22 |  | High alpha power left central–temporal (T7, C3) | Log oscillatory (FOOOF-corrected) multitaper power in 10–12 Hz, averaged across electrodes of the region |
| 23 |  | Beta power left central–temporal (T7, C3) | Log oscillatory (FOOOF-corrected) multitaper power in 13–30 Hz, averaged across electrodes of the region |
| 24 |  | Gamma power left central–temporal (T7, C3) | Log oscillatory (FOOOF-corrected) multitaper power in 30–50 Hz, averaged across electrodes of the region |
| 25 |  | Delta power central / vertex (Cz) | Log oscillatory (FOOOF-corrected) multitaper power in 1–4 Hz, averaged across electrodes of the region |
| 26 |  | Theta power central / vertex (Cz) | Log oscillatory (FOOOF-corrected) multitaper power in 4–8 Hz, averaged across electrodes of the region |
| 27 |  | Low alpha power central / vertex (Cz) | Log oscillatory (FOOOF-corrected) multitaper power in 8–10 Hz, averaged across electrodes of the region |
| 28 |  | High alpha power central / vertex (Cz) | Log oscillatory (FOOOF-corrected) multitaper power in 10–12 Hz, averaged across electrodes of the region |
| 29 |  | Beta power central / vertex (Cz) | Log oscillatory (FOOOF-corrected) multitaper power in 13–30 Hz, averaged across electrodes of the region |
| 30 |  | Gamma power central / vertex (Cz) | Log oscillatory (FOOOF-corrected) multitaper power in 30–50 Hz, averaged across electrodes of the region |
| 31 |  | Delta power right central–temporal (C4, T8) | Log oscillatory (FOOOF-corrected) multitaper power in 1–4 Hz, averaged across electrodes of the region |
| 32 |  | Theta power right central–temporal (C4, T8) | Log oscillatory (FOOOF-corrected) multitaper power in 4–8 Hz, averaged across electrodes of the region |
| 33 |  | Low alpha power right central–temporal (C4, T8) | Log oscillatory (FOOOF-corrected) multitaper power in 8–10 Hz, averaged across electrodes of the region |
| 34 |  | High alpha power right central–temporal (C4, T8) | Log oscillatory (FOOOF-corrected) multitaper power in 10–12 Hz, averaged across electrodes of the region |
| 35 |  | Beta power right central–temporal (C4, T8) | Log oscillatory (FOOOF-corrected) multitaper power in 13–30 Hz, averaged across electrodes of the region |
| 36 |  | Gamma power right central–temporal (C4, T8) | Log oscillatory (FOOOF-corrected) multitaper power in 30–50 Hz, averaged across electrodes of the region |
| 37 |  | Delta power left posterior | Log oscillatory (FOOOF-corrected) multitaper power in 1–4 Hz, averaged across electrodes of the region |
| 38 |  | Theta power left posterior | Log oscillatory (FOOOF-corrected) multitaper power in 4–8 Hz, averaged across electrodes of the region |
| 39 |  | Low alpha power left posterior | Log oscillatory (FOOOF-corrected) multitaper power in 8–10 Hz, averaged across electrodes of the region |
| 40 |  | High alpha power left posterior | Log oscillatory (FOOOF-corrected) multitaper power in 10–12 Hz, averaged across electrodes of the region |
| 41 |  | Beta power left posterior | Log oscillatory (FOOOF-corrected) multitaper power in 13–30 Hz, averaged across electrodes of the region |
| 42 |  | Gamma power left posterior | Log oscillatory (FOOOF-corrected) multitaper power in 30–50 Hz, averaged across electrodes of the region |
| 43 |  | Delta power midline parieto–occipital (Pz, Oz) | Log oscillatory (FOOOF-corrected) multitaper power in 1–4 Hz, averaged across electrodes of the region |

| No. | Modality | Feature | Description |
|---|---|---|---|
| 44 |  | Theta power midline parieto–occipital (Pz, Oz) | Log oscillatory (FOOOF-corrected) multitaper power in 4–8 Hz, averaged across electrodes of the region |
| 45 |  | Low alpha power midline parieto–occipital (Pz, Oz) | Log oscillatory (FOOOF-corrected) multitaper power in 8–10 Hz, averaged across electrodes of the region |
| 46 |  | High alpha power midline parieto–occipital (Pz, Oz) | Log oscillatory (FOOOF-corrected) multitaper power in 10–12 Hz, averaged across electrodes of the region |
| 47 |  | Beta power midline parieto–occipital (Pz, Oz) | Log oscillatory (FOOOF-corrected) multitaper power in 13–30 Hz, averaged across electrodes of the region |
| 48 |  | Gamma power midline parieto–occipital (Pz, Oz) | Log oscillatory (FOOOF-corrected) multitaper power in 30–50 Hz, averaged across electrodes of the region |
| 49 |  | Delta power right posterior | Log oscillatory (FOOOF-corrected) multitaper power in 1–4 Hz, averaged across electrodes of the region |
| 50 |  | Theta power right posterior | Log oscillatory (FOOOF-corrected) multitaper power in 4–8 Hz, averaged across electrodes of the region |
| 51 |  | Low alpha power right posterior | Log oscillatory (FOOOF-corrected) multitaper power in 8–10 Hz, averaged across electrodes of the region |
| 52 |  | High alpha power right posterior | Log oscillatory (FOOOF-corrected) multitaper power in 10–12 Hz, averaged across electrodes of the region |
| 53 |  | Beta power right posterior | Log oscillatory (FOOOF-corrected) multitaper power in 13–30 Hz, averaged across electrodes of the region |
| 54 |  | Gamma power right posterior | Log oscillatory (FOOOF-corrected) multitaper power in 30–50 Hz, averaged across electrodes of the region |
| 55 | **EDA** | Mean raw EDA | Mean of z-scored, detrended skin conductance per segment |
| 56 | | Mean tonic EDA (SCL) | Mean of cvxEDA tonic component per segment |
| 57 | | Mean phasic EDA (SCR) | Mean of cvxEDA phasic component per segment |
| 58 | | SD raw EDA | Standard deviation of z-scored skin conductance per segment |
| 59 | | SD tonic EDA | Standard deviation of tonic component per segment |
| 60 | | SD phasic EDA | Standard deviation of phasic component per segment |
| 61 | **ECG / HR** | Mean heart rate | Mean of beat-derived instantaneous heart rate (Pan–Tompkins R-peaks) per segment |
| 62 | | SD heart rate | Standard deviation of instantaneous heart rate per segment |
| 63 | **TREV** | Contractility | Mean first difference of the TREV waveform per segment |
| 64 | | Cardiac output index | Integrated TREV velocity × mean heart rate of the segment |
| 65 | **BP** | Mean systolic BP | Mean beat-to-beat systolic pressure per segment |
| 66 | | Max systolic BP | Maximum systolic pressure per segment |
| 67 | | Min systolic BP | Minimum systolic pressure per segment |
| 68 | | SD systolic BP | Standard deviation of systolic pressure per segment |
| 69 | | Kurtosis systolic BP | Kurtosis of systolic pressure per segment |
| 70 | | Mean diastolic BP | Mean beat-to-beat diastolic pressure per segment |
| 71 | | Max diastolic BP | Maximum diastolic pressure per segment |
| 72 | | Min diastolic BP | Minimum diastolic pressure per segment |

| No. | Modality | Feature | Description |
|---|---|---|---|
| 73 | | SD diastolic BP | Standard deviation of diastolic pressure per segment |
| 74 | | Kurtosis diastolic BP | Kurtosis of diastolic pressure per segment |
| 75 | **EOG** | Blink rate | Blinks per minute from band-passed (1–10 Hz) vertical EOG peak detection |
| 76 | **EGG** | Waveform length | Sum of absolute first differences of the filtered EGG per segment |
| 77 | | Average amplitude change | Mean of absolute first differences per segment |
| 78 | | Mean fractal length | log10 of the root of summed squared first differences per segment |
| 79 | | Mean absolute value | Mean of absolute signal amplitude per segment |
| 80 | | Root mean square | RMS amplitude of the filtered EGG per segment |
| 81 | **Respiration** | Respiration rate | Breaths per minute from peak detection on the detrended respiration trace |
| 82 | | Waveform length | Sum of absolute first differences per segment |
| 83 | | Average amplitude change | Mean of absolute first differences per segment |
| 84 | | Mean fractal length | log10 of the root of summed squared first differences per segment |
| 85 | | Mean absolute value | Mean of absolute signal amplitude per segment |
| 86 | | Root mean square | RMS amplitude of the respiration trace per segment |
| 87 | **EMG** | Mean of corrugator | Mean of the preprocessed Corrugator EMG envelope per segment |
| 88 | | Max of corrugator | Maximum of the Corrugator EMG envelope per segment |
| 89 | | Min of corrugator | Minimum of the Corrugator EMG envelope per segment |
| 90 | | SD of corrugator | Standard deviation of the Corrugator EMG envelope per segment |
| 91 | | Kurtosis of corrugator | Kurtosis of the Corrugator EMG envelope per segment |
| 92 | | Variance of change of corrugator | Mean moving variance of the differenced Corrugator EMG envelope per segment |
| 93 | | Mean absolute value of corrugator | Mean absolute amplitude of the Corrugator EMG envelope per segment |
| 94 | | RMS of corrugator | RMS amplitude of the Corrugator EMG envelope per segment |
| 95 | | Mean of zygomaticus | Mean of the preprocessed Zygomaticus EMG envelope per segment |
| 96 | | Max of zygomaticus | Maximum of the Zygomaticus EMG envelope per segment |
| 97 | | Min of zygomaticus | Minimum of the Zygomaticus EMG envelope per segment |
| 98 | | SD of zygomaticus | Standard deviation of the Zygomaticus EMG envelope per segment |
| 99 | | Kurtosis of zygomaticus | Kurtosis of the Zygomaticus EMG envelope per segment |
| 100 | | Variance of change of zygomaticus | Mean moving variance of the differenced Zygomaticus EMG envelope per segment |
| 101 | | Mean absolute value of zygomaticus | Mean absolute amplitude of the Zygomaticus EMG envelope per segment |
| 102 | | RMS zof ygomaticus | RMS amplitude of the Zygomaticus EMG envelope per segment |

**Supplementary Table T2.** Emotion-versus-baseline accuracy relative to surrogate models. Each cell gives mean accuracy ± s.e.m. across cross-validation runs and, in brackets, the 2.5th–97.5th percentile interval of label-permuted surrogate models (ppt 1–4 labelled emo01, emo05, emo06 and emo07). All models exceeded the surrogate upper bound; empty cells are emotions not modelled for ppt 4.

| | Admiration | Adoration | AestheticAppreciation | Amusement | Awe | Awkwardness | Boredom | Calmness |
|---|---|---|---|---|---|---|---|---|
| ppt 1 | 0.975 ± 0.005 [0.440, 0.569] | 0.983 ± 0.003 [0.447, 0.559] | 0.981 ± 0.003 [0.451, 0.559] | 0.986 ± 0.002 [0.445, 0.555] | 0.976 ± 0.003 [0.447, 0.549] | 0.976 ± 0.007 [0.436, 0.564] | 0.998 ± 0.002 [0.413, 0.588] | 0.968 ± 0.006 [0.440, 0.582] |
| ppt 2 | 0.941 ± 0.010 [0.454, 0.567] | 0.967 ± 0.007 [0.425, 0.562] | 0.971 ± 0.009 [0.467, 0.537] | 0.969 ± 0.010 [0.449, 0.555] | 0.980 ± 0.004 [0.446, 0.546] | 0.971 ± 0.010 [0.448, 0.542] | 0.954 ± 0.011 [0.443, 0.566] | 0.955 ± 0.007 [0.458, 0.541] |
| ppt 3 | 0.866 ± 0.014 [0.472, 0.532] | 0.851 ± 0.013 [0.458, 0.528] | 0.889 ± 0.011 [0.466, 0.541] | 0.902 ± 0.012 [0.464, 0.538] | 0.884 ± 0.009 [0.470, 0.535] | 0.873 ± 0.016 [0.470, 0.535] | 0.902 ± 0.012 [0.462, 0.541] | 0.901 ± 0.011 [0.464, 0.531] |
| ppt 4 | 0.881 ± 0.010 [0.446, 0.553] | 0.864 ± 0.014 [0.463, 0.537] | 0.851 ± 0.017 [0.443, 0.555] | 0.902 ± 0.012 [0.460, 0.530] | 0.850 ± 0.019 [0.452, 0.543] | 0.885 ± 0.012 [0.455, 0.546] | | |

| | Confusion | Entrancement | Excitement | Interest | Joy | Relief | Satisfaction |
|---|---|---|---|---|---|---|---|
| ppt 1 | 0.946 ± 0.017 [0.445, 0.570] | 0.979 ± 0.004 [0.441, 0.566] | 0.971 ± 0.004 [0.454, 0.553] | 0.981 ± 0.002 [0.453, 0.541] | 0.949 ± 0.010 [0.441, 0.558] | 0.954 ± 0.007 [0.429, 0.572] | 0.977 ± 0.003 [0.453, 0.563] |
| ppt 2 | 0.948 ± 0.008 [0.450, 0.544] | 0.975 ± 0.003 [0.455, 0.541] | 0.978 ± 0.005 [0.441, 0.546] | 0.973 ± 0.008 [0.456, 0.542] | 0.967 ± 0.003 [0.453, 0.553] | 0.927 ± 0.014 [0.436, 0.568] | 0.966 ± 0.009 [0.451, 0.539] |
| ppt 3 | 0.887 ± 0.014 [0.456, 0.532] | 0.891 ± 0.011 [0.471, 0.532] | 0.874 ± 0.012 [0.466, 0.530] | 0.892 ± 0.012 [0.472, 0.530] | 0.889 ± 0.014 [0.462, 0.534] | 0.838 ± 0.013 [0.464, 0.533] | 0.895 ± 0.013 [0.469, 0.533] |
| ppt 4 | 0.857 ± 0.008 [0.442, 0.549] | 0.882 ± 0.012 [0.467, 0.541] | 0.860 ± 0.011 [0.440, 0.554] | 0.909 ± 0.009 [0.455, 0.553] | | | |

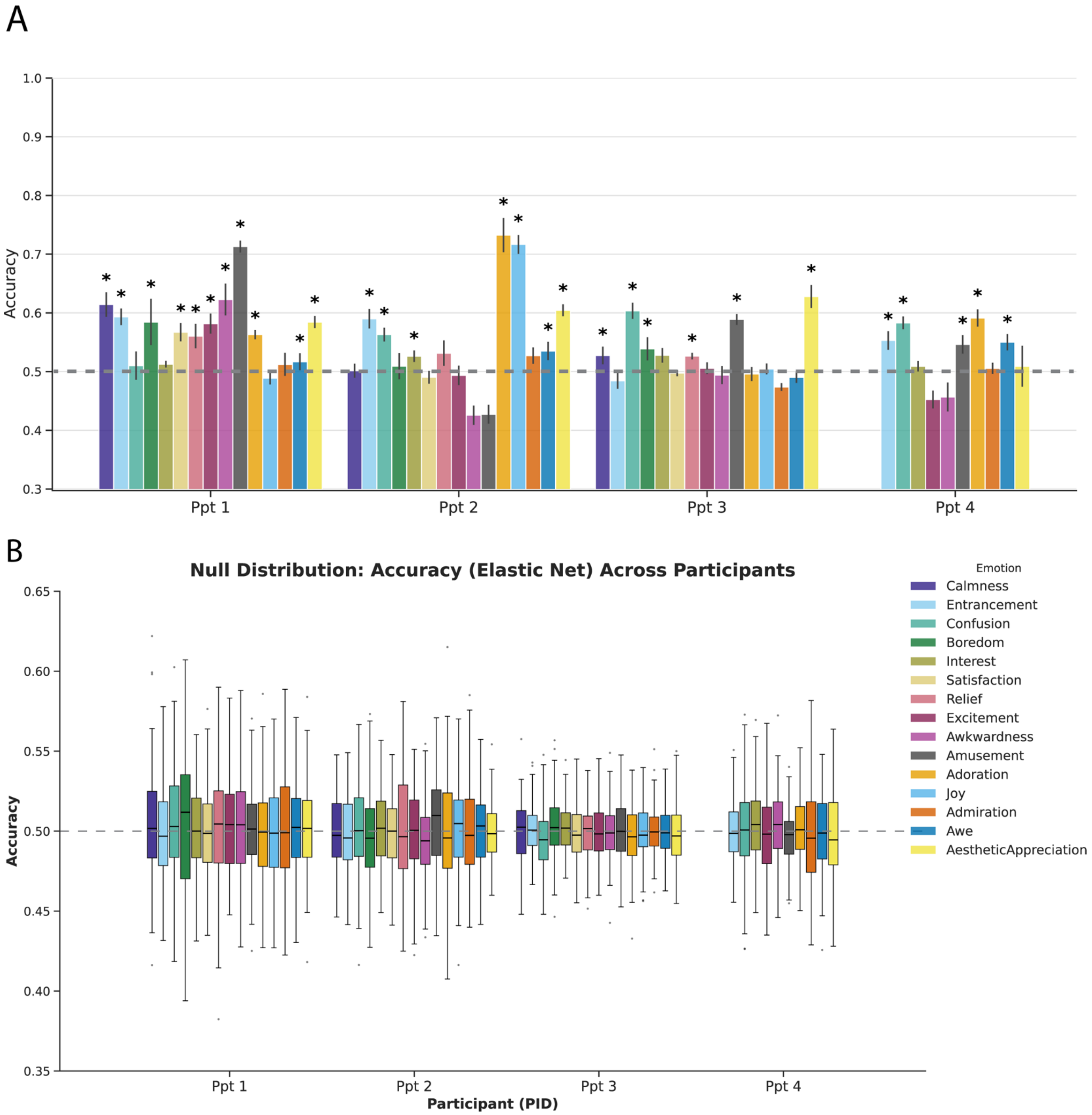


**Fig. S2.** One-vs-all classification. A, Accuracy (mean ± s.e.m. across cross-validation runs) of LR-EN models contrasting trials in which each emotion was rated with all other trials; asterisks mark models exceeding the 97.5th percentile of label-permuted surrogates, and the dashed line marks chance. B, Accuracy distributions of the surrogate models, trained identically with shuffled labels.

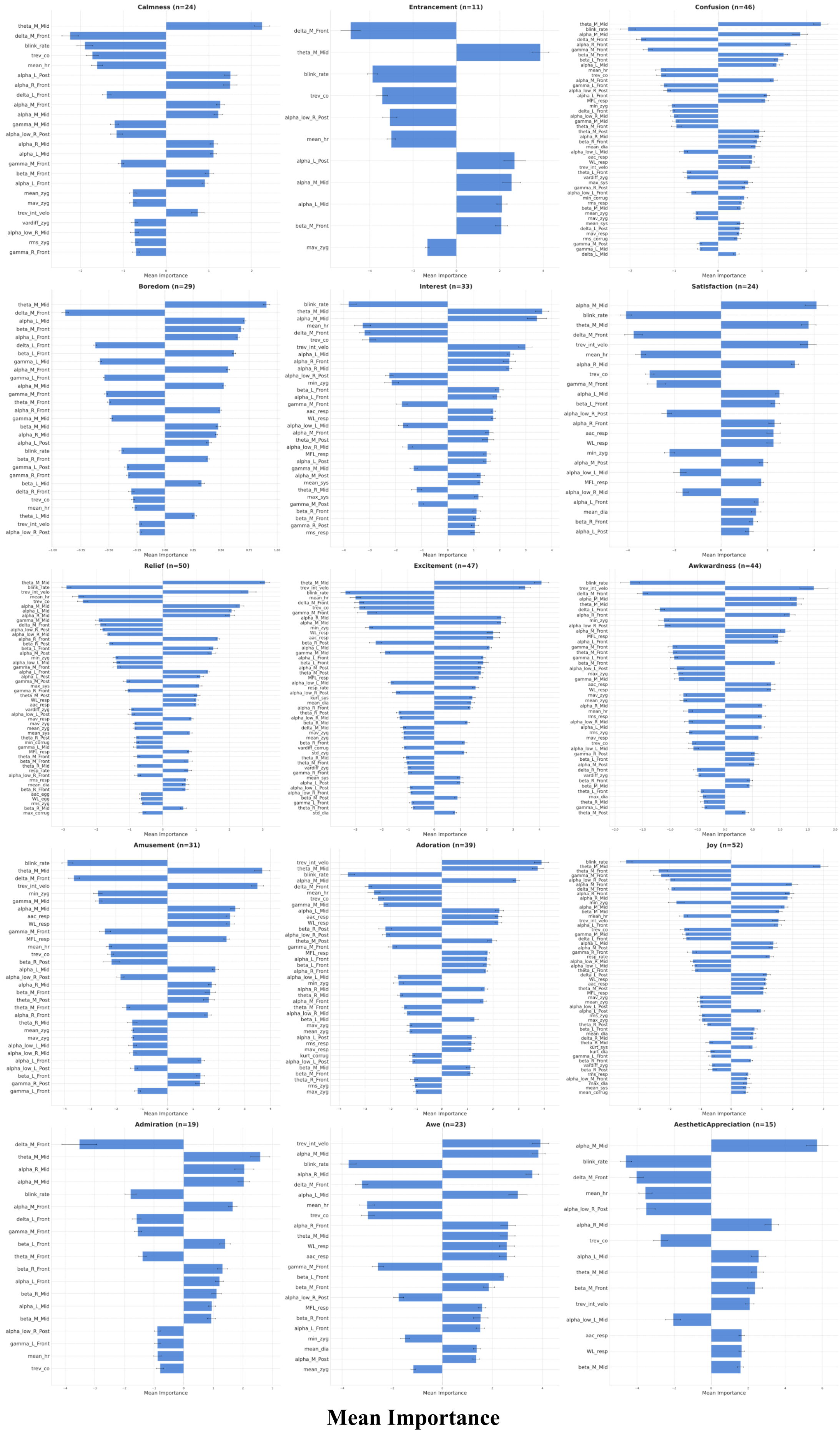


**Fig. S3.** Stable elastic-net feature weights for each emotion versus baseline in participant 1. Bars show mean signed weights (± s.e.m. across 25 cross-validation runs) of features with non-zero weight in ≥90% of runs, trimmed to the top 90% of cumulative absolute importance (Methods).

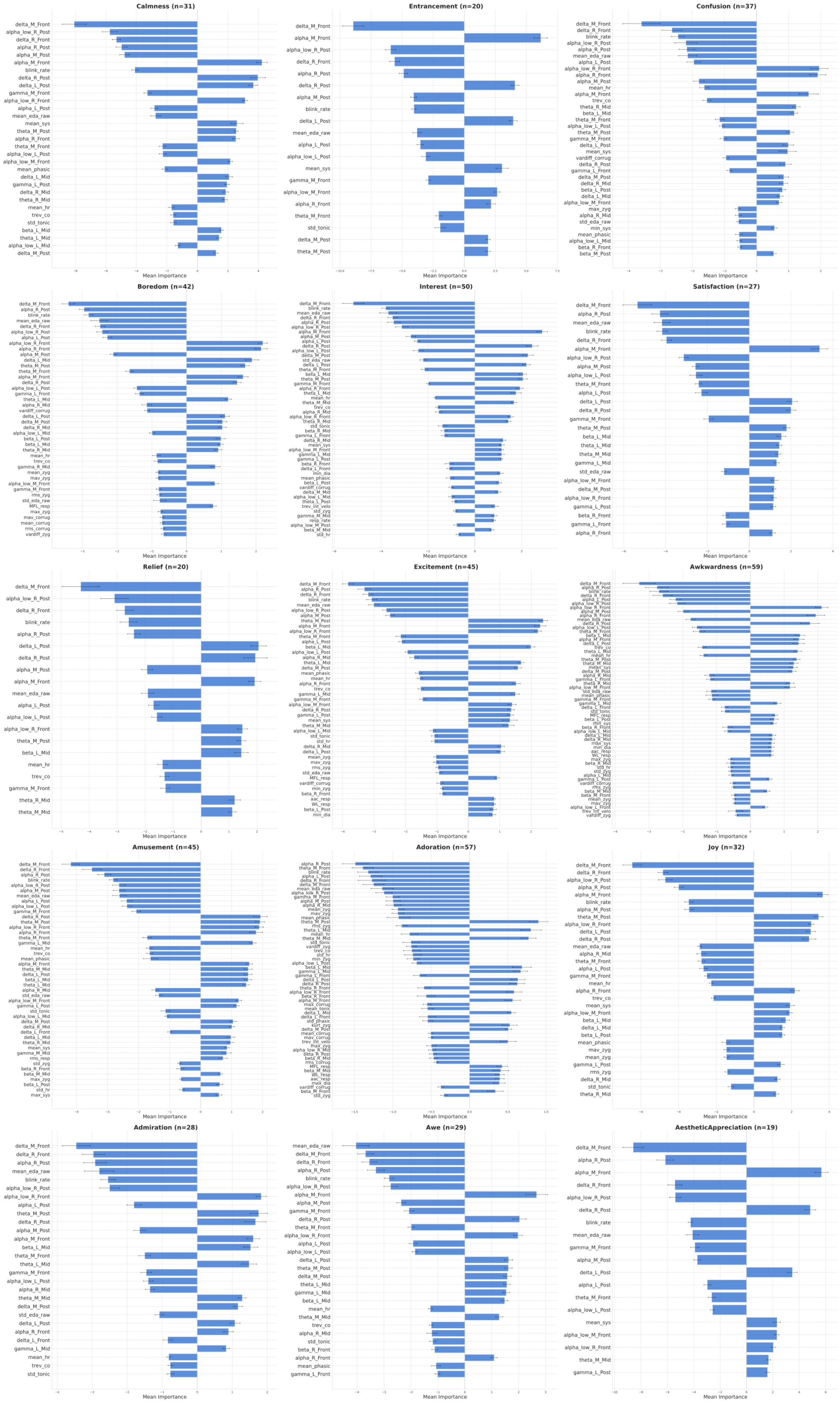




**Mean Importance**

**Fig. S4.** Stable elastic-net feature weights for each emotion versus baseline in participant 2. Legend is similar as Fig. S3

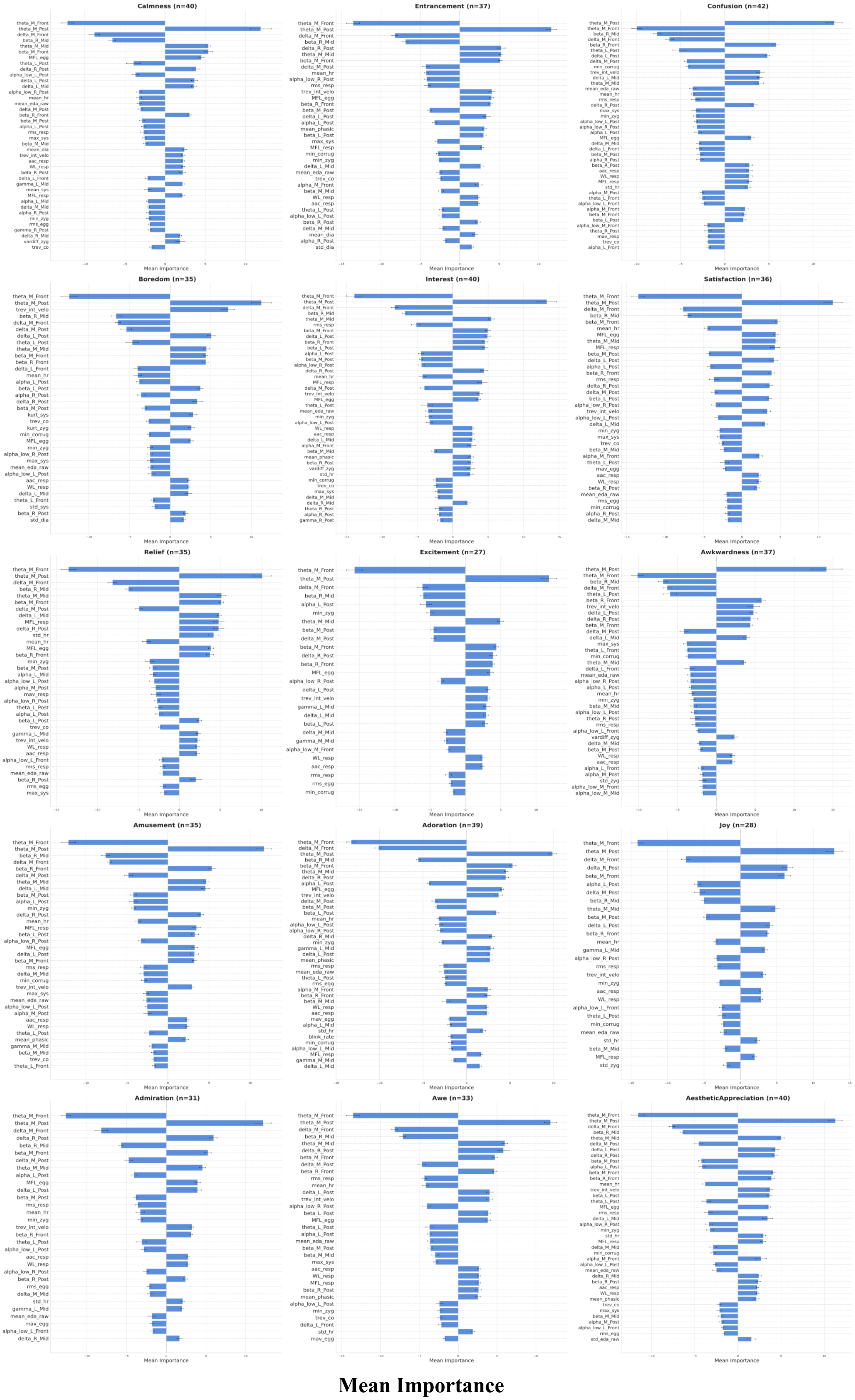


**Fig. S5.** Stable elastic-net feature weights for each emotion versus baseline in participant 3. Legend is similar as Fig. S3

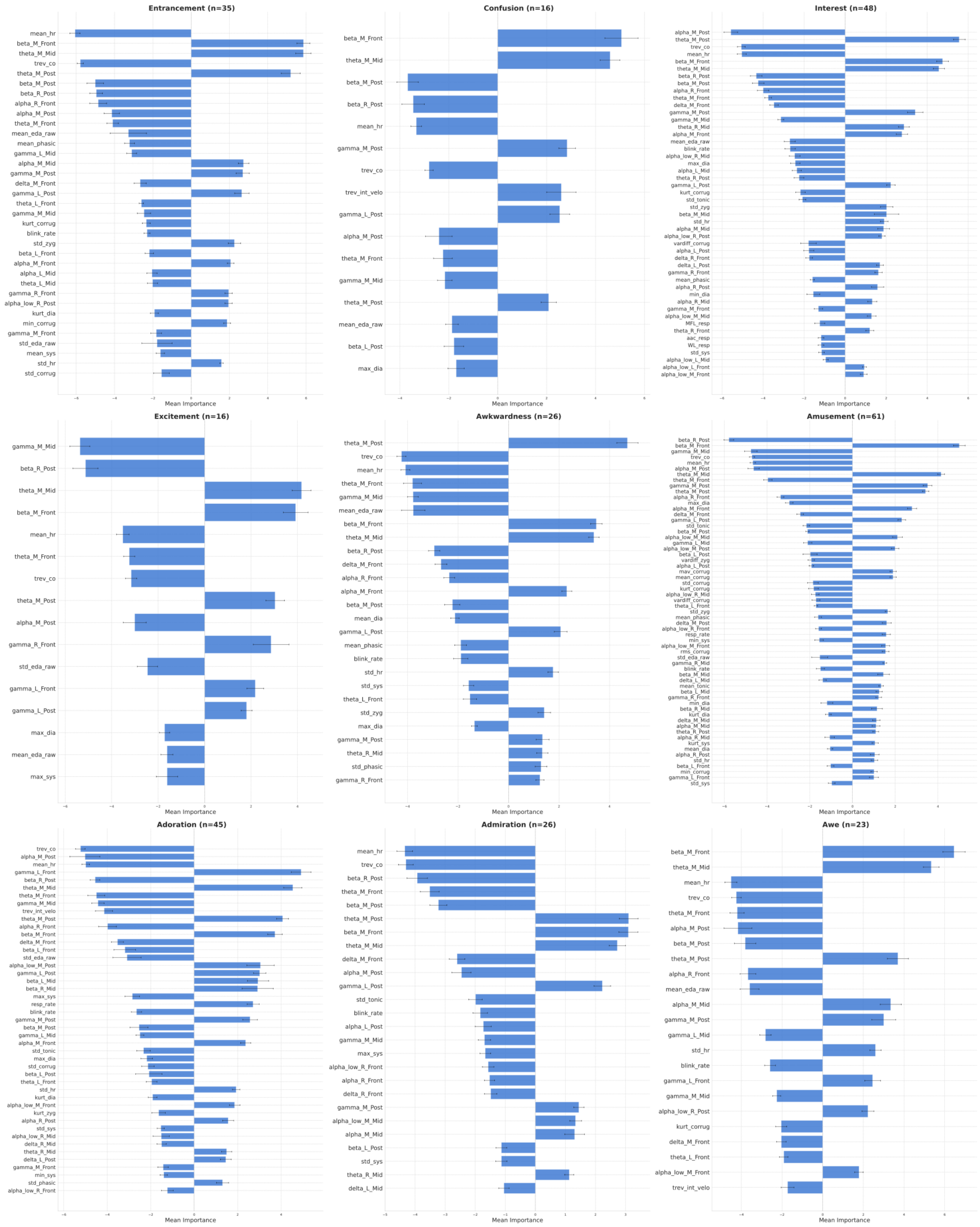


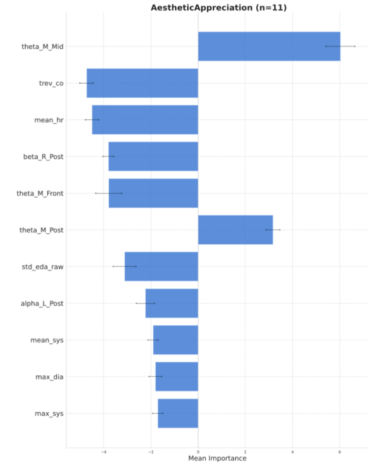


**Fig. S6.** Stable elastic-net feature weights for each emotion versus baseline in participant 4. Bars show mean signed weights (± s.e.m. across 25 cross-validation runs) of features with non-zero weight in ≥90% of runs, trimmed to the top 90% of cumulative absolute importance (Methods). Only 10 emotions were modelled for this participant.

**Mean Importance**

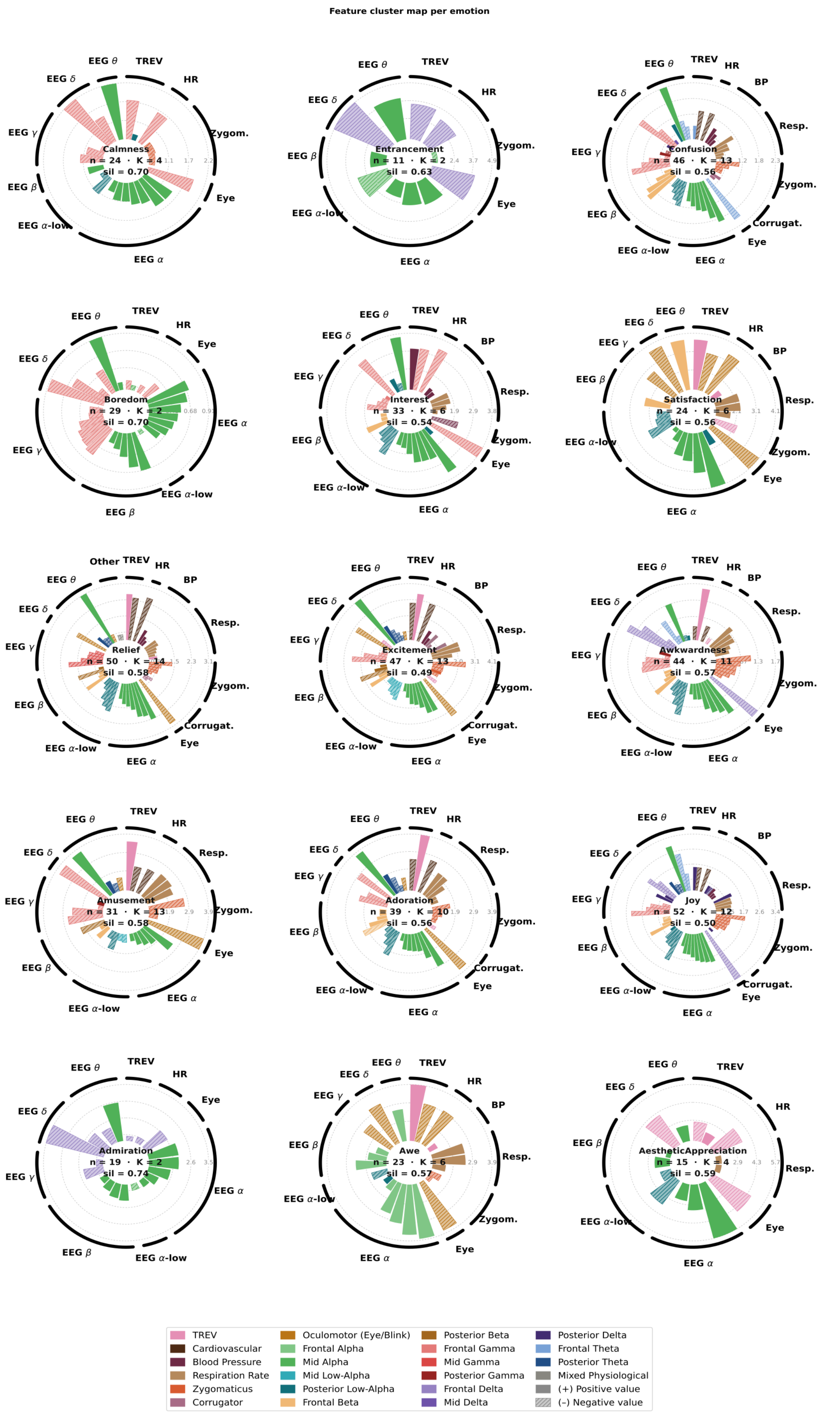


**Fig. S7.** Circular feature-cluster maps for each emotion in participant 1, plotted as in Fig. 4. Emotion name, number of stable features (n), number of clusters (K) and silhouette score (sil) are shown inside each map.

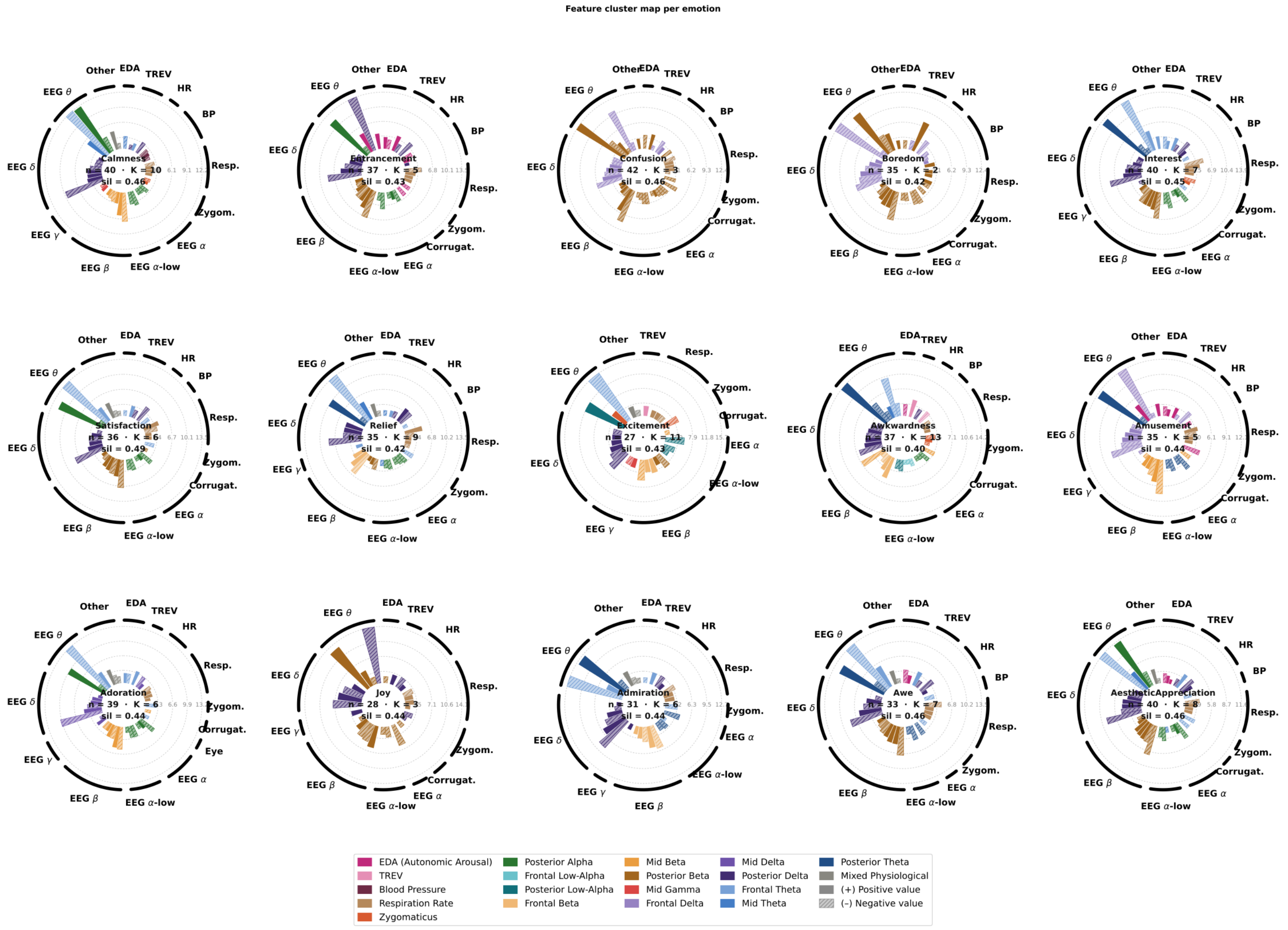


**Fig. S8.** Circular feature-cluster maps for each emotion in participant 3, plotted as in Fig. 4. Emotion name, number of stable features (n), number of clusters (K) and silhouette score (sil) are shown inside each map.

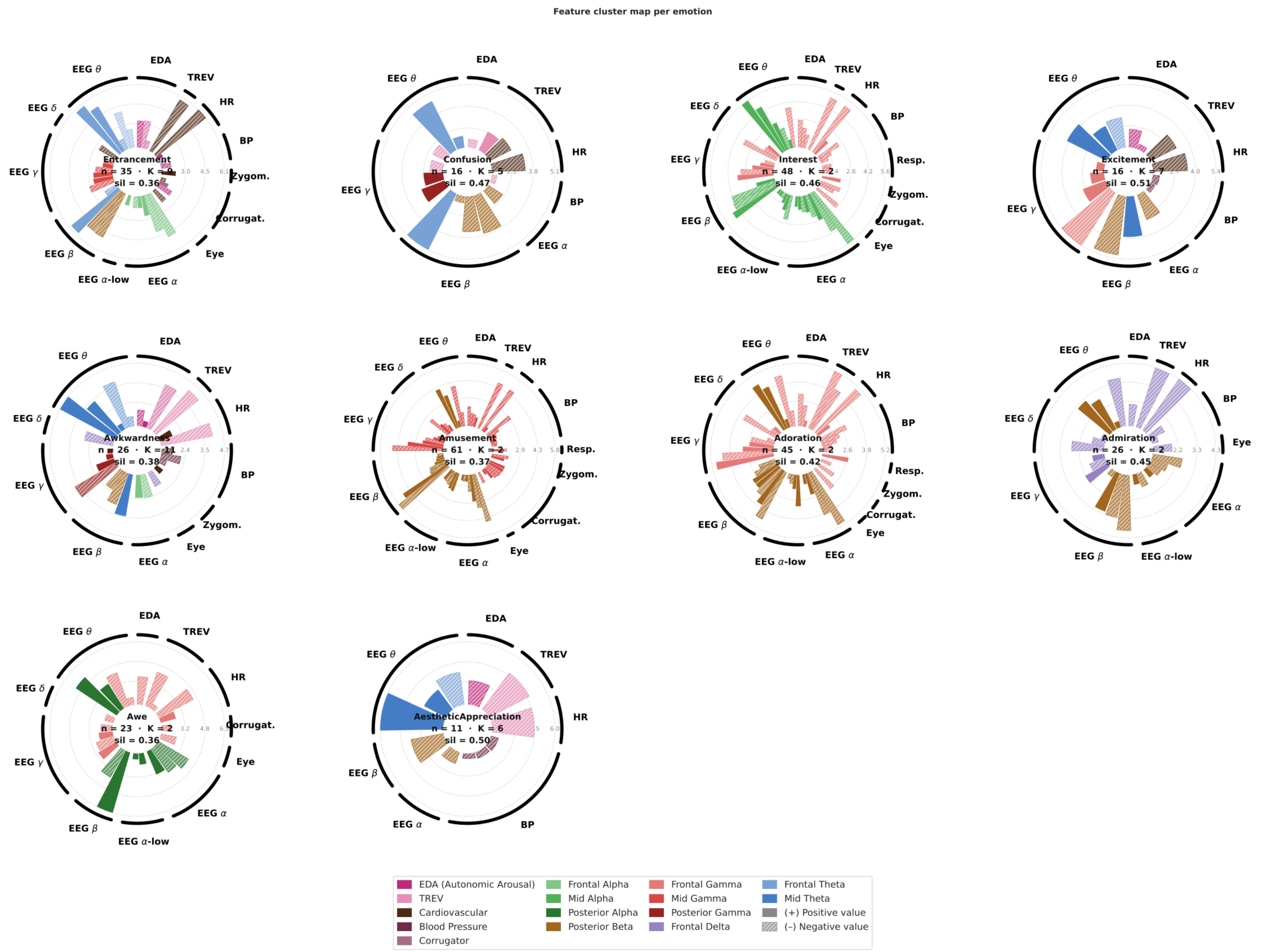


**Fig. S9.** Circular feature-cluster maps for each emotion in participant 4, plotted as in Fig. 4. Emotion name, number of stable features (n), number of clusters (K) and silhouette score (sil) are shown inside each map. Only 10 emotions were modelled for this participant.

**Supplementary Table T3. Shared and emotion-specific feature clusters in participant 1.** As in Table 1.

| | Admiration | Adoration | AestheticAppreciation | Amusement | Awe | Awkwardness | Boredom | Calmness | Confusion | Entrancement | Excitement | Interest | Joy | Relief | Satisfaction |
|---|---|---|---|---|---|---|---|---|---|---|---|---|---|---|---|
| Mid Alpha | +2.59 (n=9) | +3.75 (n=7) | +5.73 (n=5) | +3.69 (n=5) | | +1.30 (n=7) | +0.91 (n=12) | +2.24 (n=9) | +2.33 (n=7) | +3.88 (n=4) | +4.08 (n=7) | +3.63 (n=8) | +2.91 (n=9) | +3.07 (n=8) | +4.11 (n=4) |
| Frontal Delta | -3.52 (n=5) | | | | | -1.73 (n=3) | | | | -4.88 (n=4) | | | -3.41 (n=3) | | |
| TREV | | +3.90 (n=1) | -4.61 (n=5) | +3.50 (n=1) | +3.92 (n=1) | +1.61 (n=1) | | | | | +3.46 (n=3) | | | +2.57 (n=1) | +3.75 (n=2) |
| Oculomotor (Eye/Blink) | | -3.67 (n=2) | | -3.89 (n=2) | -3.74 (n=5) | | | | | | -3.38 (n=2) | | | -2.90 (n=2) | -4.08 (n=5) |
| Frontal Alpha | | | | | +3.84 (n=7) | | | | | | | | | | |
| Frontal Gamma | | -2.87 (n=3) | | -3.64 (n=3) | | -0.96 (n=3) | -0.89 (n=9) | -2.24 (n=7) | -1.75 (n=5) | | -2.56 (n=2) | -3.81 (n=6) | -2.27 (n=3) | | |
| Frontal Beta | | -2.21 (n=3) | | +1.66 (n=1) | | +0.90 (n=1) | | | +1.49 (n=3) | | +1.87 (n=2) | +1.98 (n=1) | +1.56 (n=2) | +1.52 (n=1) | +3.76 (n=2) |
| Posterior Low-Alpha | | -2.18 (n=3) | -3.52 (n=1) | -1.83 (n=1) | -1.75 (n=1) | -1.10 (n=4) | | -1.16 (n=1) | -1.16 (n=5) | | | -2.25 (n=4) | -1.98 (n=4) | -1.81 (n=4) | -2.33 (n=3) |
| Cardiovascular | | -2.65 (n=2) | | -2.28 (n=2) | | -0.67 (n=2) | | | -1.30 (n=2) | | -2.99 (n=2) | | -1.55 (n=2) | -2.54 (n=2) | |
| Blood Pressure | | | | | | | | | +0.85 (n=2) | | +1.57 (n=3) | +2.99 (n=2) | +0.72 (n=1) | +1.09 (n=2) | |
| Zygomaticus | | -1.69 (n=3) | | -2.70 (n=2) | | -1.11 (n=5) | | -0.78 (n=1) | -1.05 (n=2) | | -2.48 (n=4) | | -1.78 (n=5) | -1.42 (n=4) | |
| Respiration Rate | | +2.21 (n=3) | | +2.44 (n=3) | +2.58 (n=3) | +0.96 (n=5) | | | +1.07 (n=3) | | +2.24 (n=3) | +1.75 (n=3) | +1.12 (n=3) | +1.01 (n=4) | +2.26 (n=2) |
| Frontal Theta | | | | | | -0.95 (n=1) | | | -2.05 (n=4) | | | | -2.35 (n=2) | | |
| Posterior Beta | | | | -2.18 (n=1) | | | | | | | -2.23 (n=2) | | | -1.61 (n=1) | |
| Posterior Theta | | +1.97 (n=2) | | +1.61 (n=2) | | | | | | | +1.78 (n=2) | | +1.06 (n=2) | +1.03 (n=2) | |
| Mid Gamma | | | | | | | | | | | | | | -1.93 (n=5) | |
| Mid Low-Alpha | | | | | | | | | | | -1.66 (n=3) | | | | |
| Posterior Delta | | | | | | | | | | | | | +1.54 (n=3) | | |
| Corrugator | | | | | | | | | | | | | | -0.80 (n=1) | |
| Posterior Gamma | | | | | | +0.54 (n=1) | | | +0.62 (n=1) | | | | | | |

**Supplementary Table T4. Shared and emotion-specific feature clusters in participant 4.** As in Table 1; only 10 emotions were modelled.

| | Admiration | Adoration | AestheticAppreciation | Amusement | Awe | Awkwardness | Confusion | Entrancement | Excitement | Interest |
|---|---|---|---|---|---|---|---|---|---|---|
| Posterior Alpha | | | | | +6.46 (n=7) | | | | | |
| Cardiovascular | | | | | | +1.77 (n=2) | -3.33 (n=2) | -6.06 (n=5) | -3.52 (n=2) | |
| Mid Theta | | | +6.03 (n=2) | | | +4.71 (n=3) | | | +4.18 (n=3) | |
| Frontal Theta | | | -3.79 (n=1) | | | -3.80 (n=1) | +5.06 (n=2) | +5.86 (n=7) | -3.25 (n=1) | |
| Posterior Beta | -3.93 (n=9) | -5.03 (n=13) | -3.80 (n=2) | -5.79 (n=17) | | -2.94 (n=2) | -3.68 (n=3) | -5.01 (n=2) | -5.13 (n=2) | |
| Mid Alpha | | | | | | | | | | -5.57 (n=17) |
| Frontal Gamma | | -5.24 (n=20) | | | -4.51 (n=10) | | | | -5.37 (n=3) | -5.06 (n=19) |
| Frontal Alpha | | | | | | -2.35 (n=2) | | -4.85 (n=5) | | |
| Mid Gamma | | | | -4.76 (n=28) | | | | -3.11 (n=4) | | |
| TREV | | | -4.72 (n=2) | | | -4.24 (n=2) | +2.60 (n=3) | | | |
| Frontal Delta | -4.34 (n=10) | | | | | -2.68 (n=2) | | | | |
| Posterior Gamma | | | | | | -3.78 (n=2) | +2.84 (n=2) | | | |
| EDA (Autonomic Arousal) | | | -3.12 (n=1) | | | -3.77 (n=2) | | -3.28 (n=3) | -2.46 (n=1) | |
| Blood Pressure | | | | | | -2.12 (n=1) | | | | |

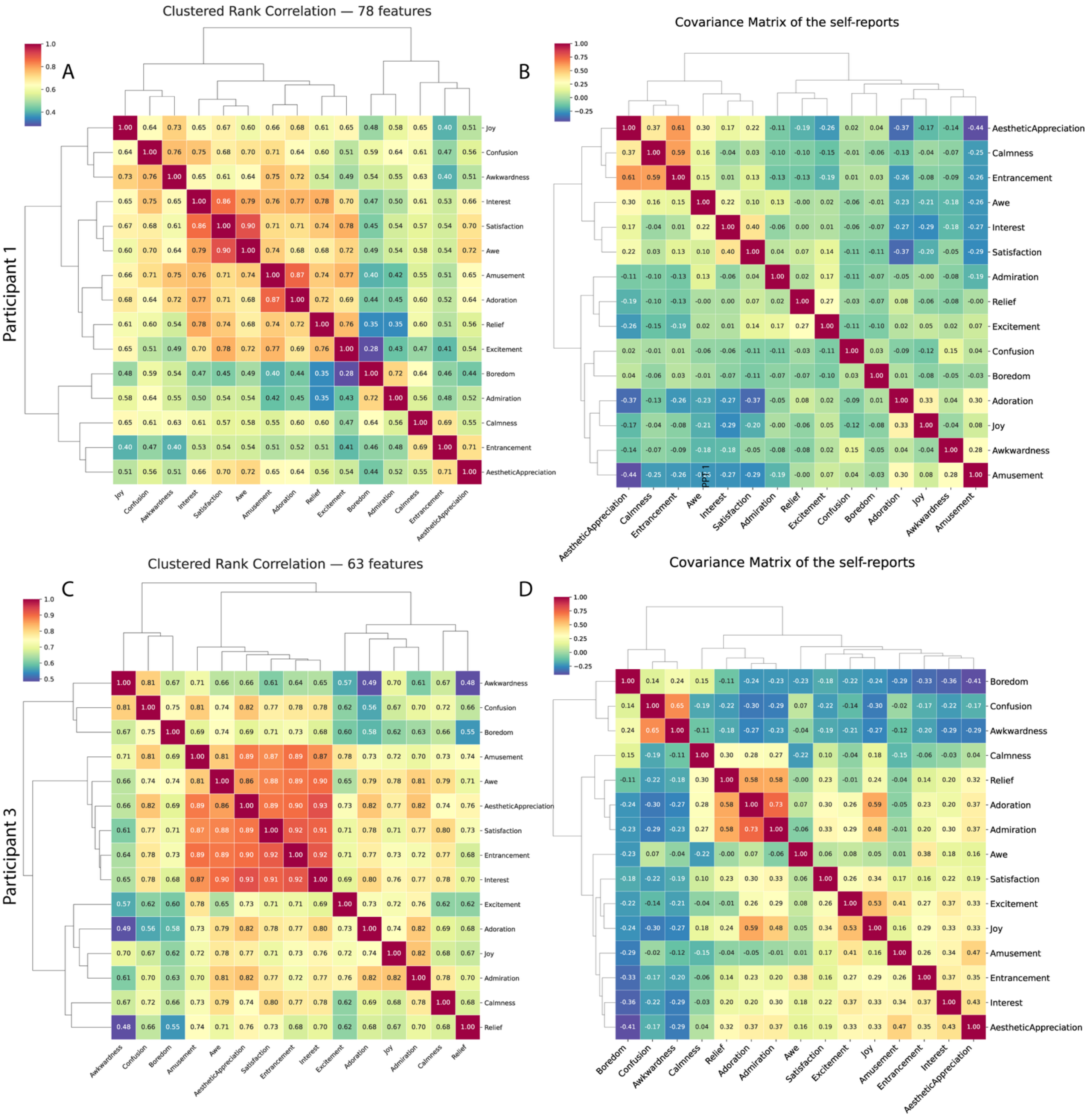


**Fig. S10.** Neurophysiological (A, C) and self-report (B, D) emotion similarity and clustering for participant 1 (A, B) and participant 3 (C, D), computed as in Fig. 5A,B. In participant 1, Awkwardness, Joy and Confusion clustered together neurophysiologically.

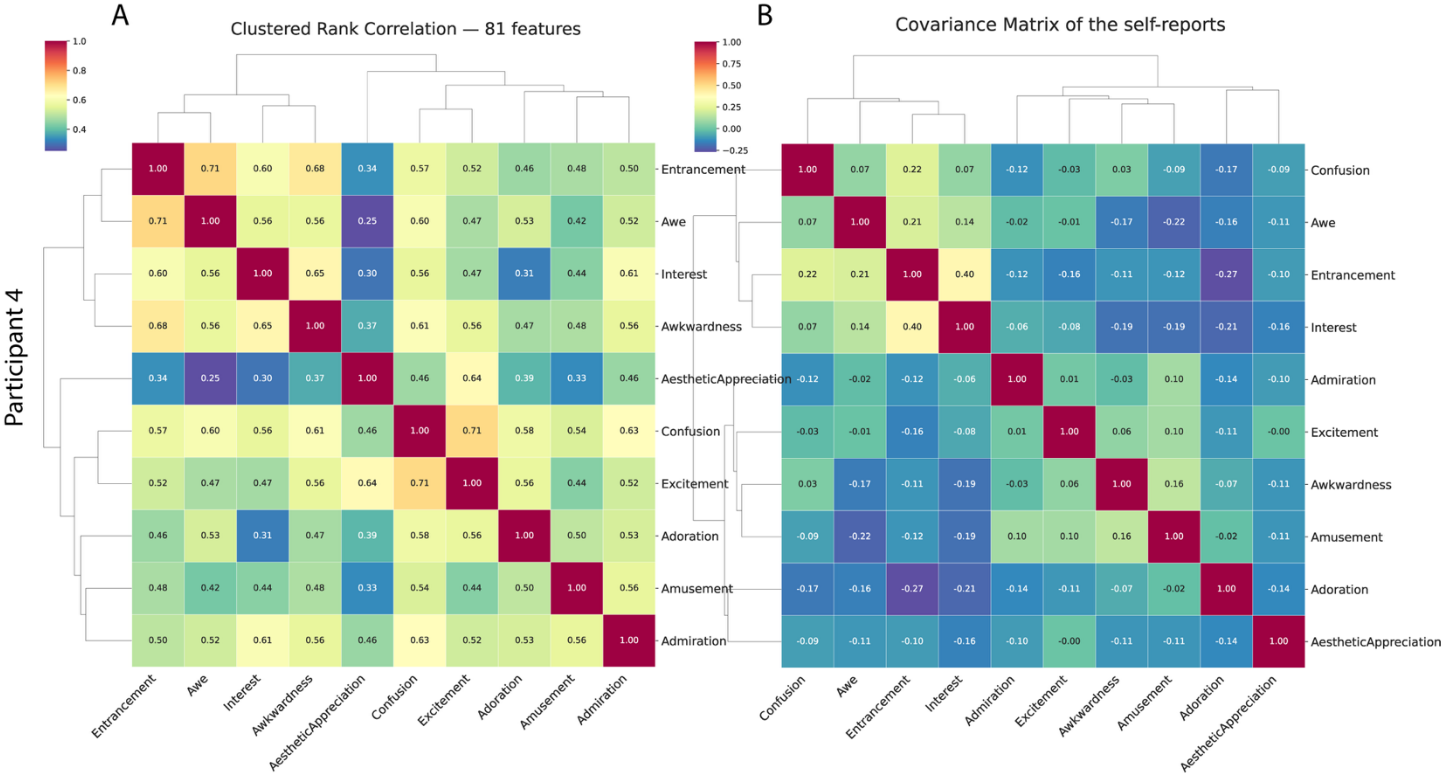


**Fig. S11.** As in Fig. S10, for participant 4 (10 emotions). Were modeled due to sample size.

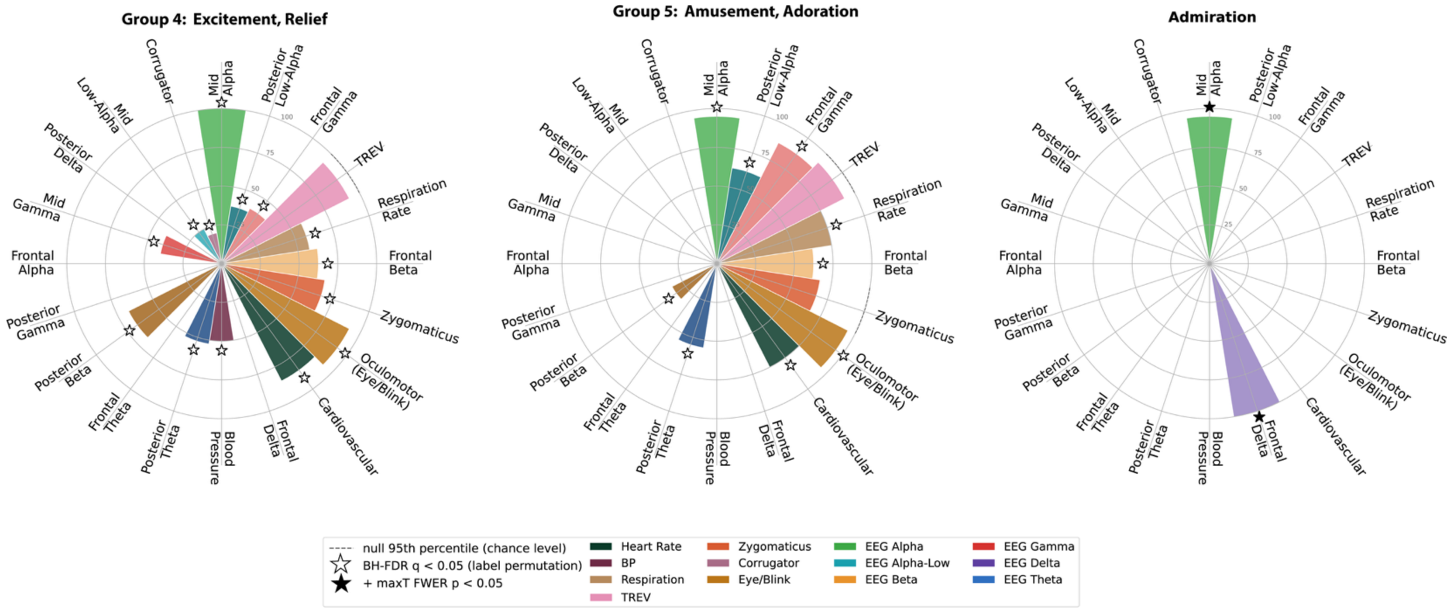


**Fig. S12.** Rose plots for the remaining emotion groups of participant 1, computed and tested as in Fig. 5D (dashed ring, null 95th percentile; open stars, FDR q < 0.05; filled stars, also max-T family-wise p < 0.05).

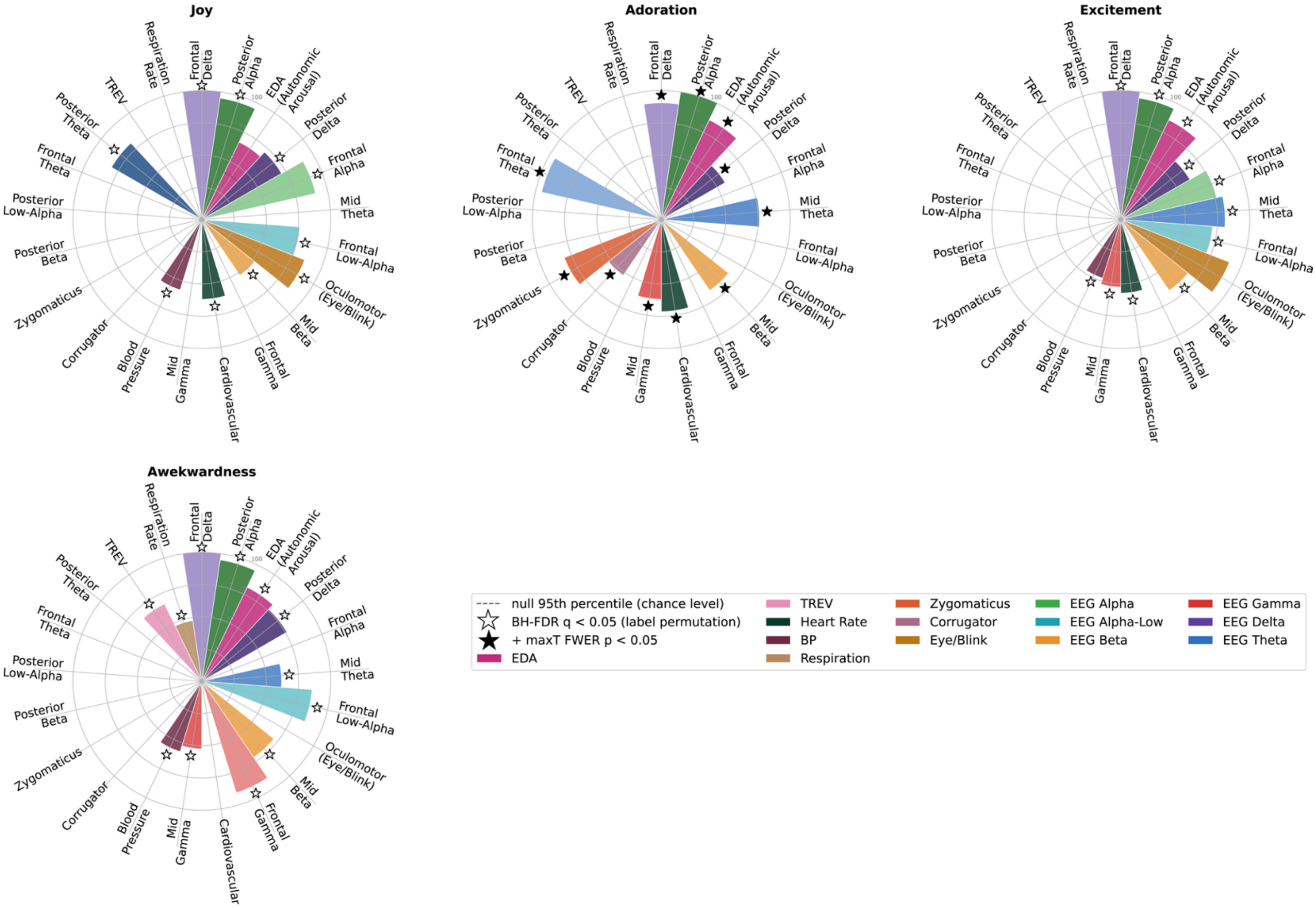


**Fig. S13.** Rose plots for the remaining emotion groups of participant 2, computed and tested as in Fig. 5D (dashed ring, null 95th percentile; open stars, FDR q < 0.05; filled stars, also max-T family-wise p < 0.05).

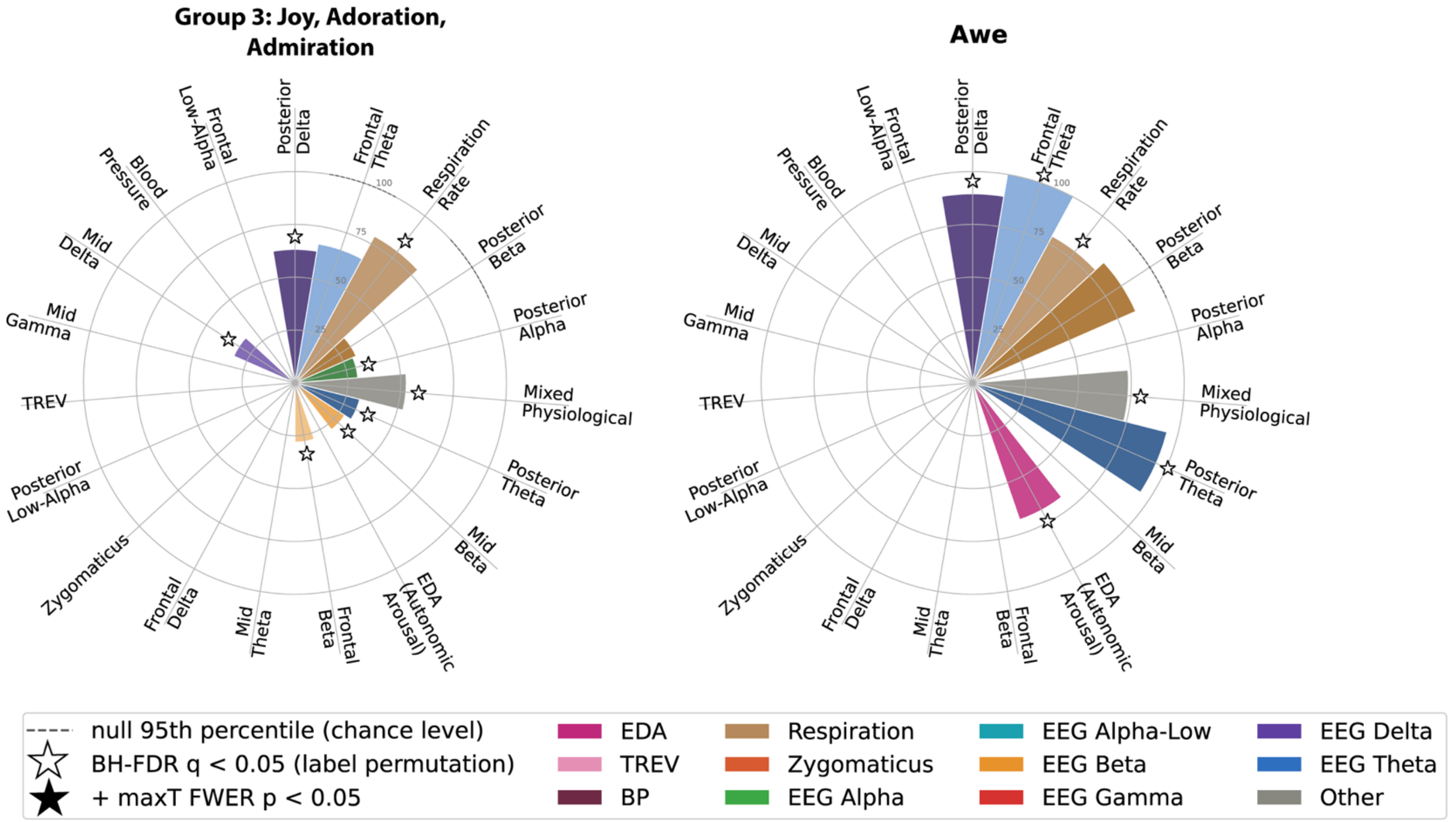


**Fig. S14.** Rose plots for the remaining emotion groups of participant 3, computed and tested as in Fig. 5D (dashed ring, null 95th percentile; open stars, FDR q < 0.05; filled stars, also max-T family-wise p < 0.05).

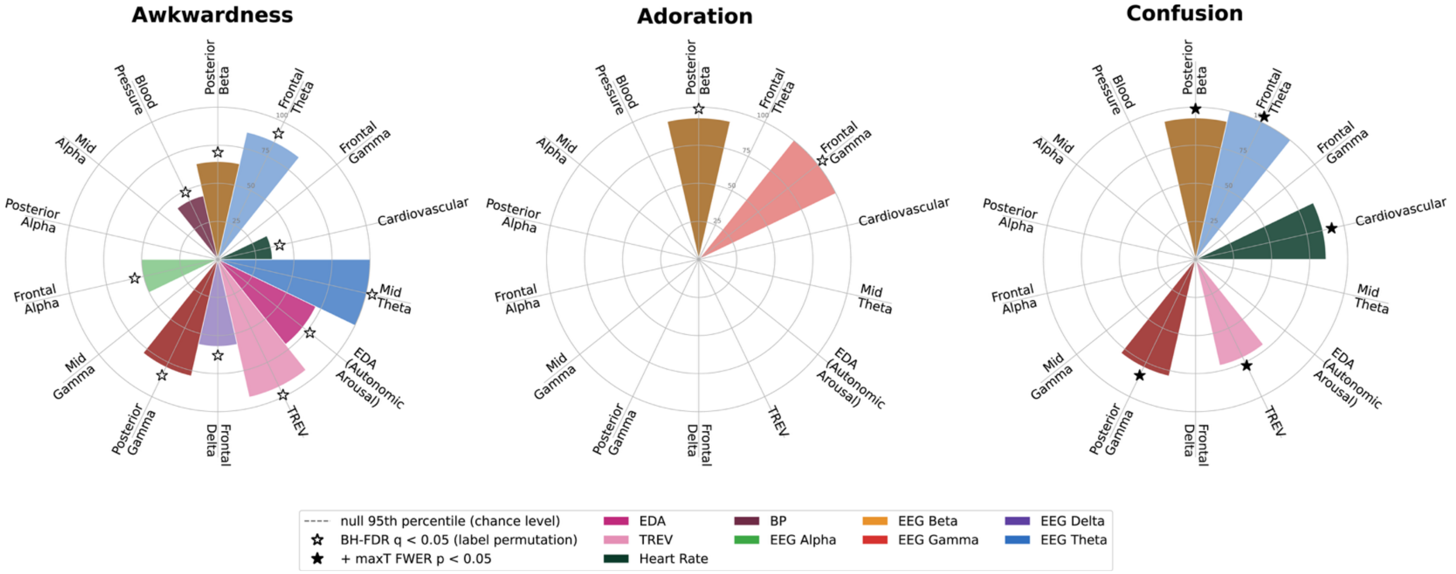


**Fig. S15.** Rose plots for the remaining emotion groups of participant 4, computed and tested as in Fig. 5D (dashed ring, null 95th percentile; open stars, FDR q < 0.05; filled stars, also max-T family-wise p < 0.05).

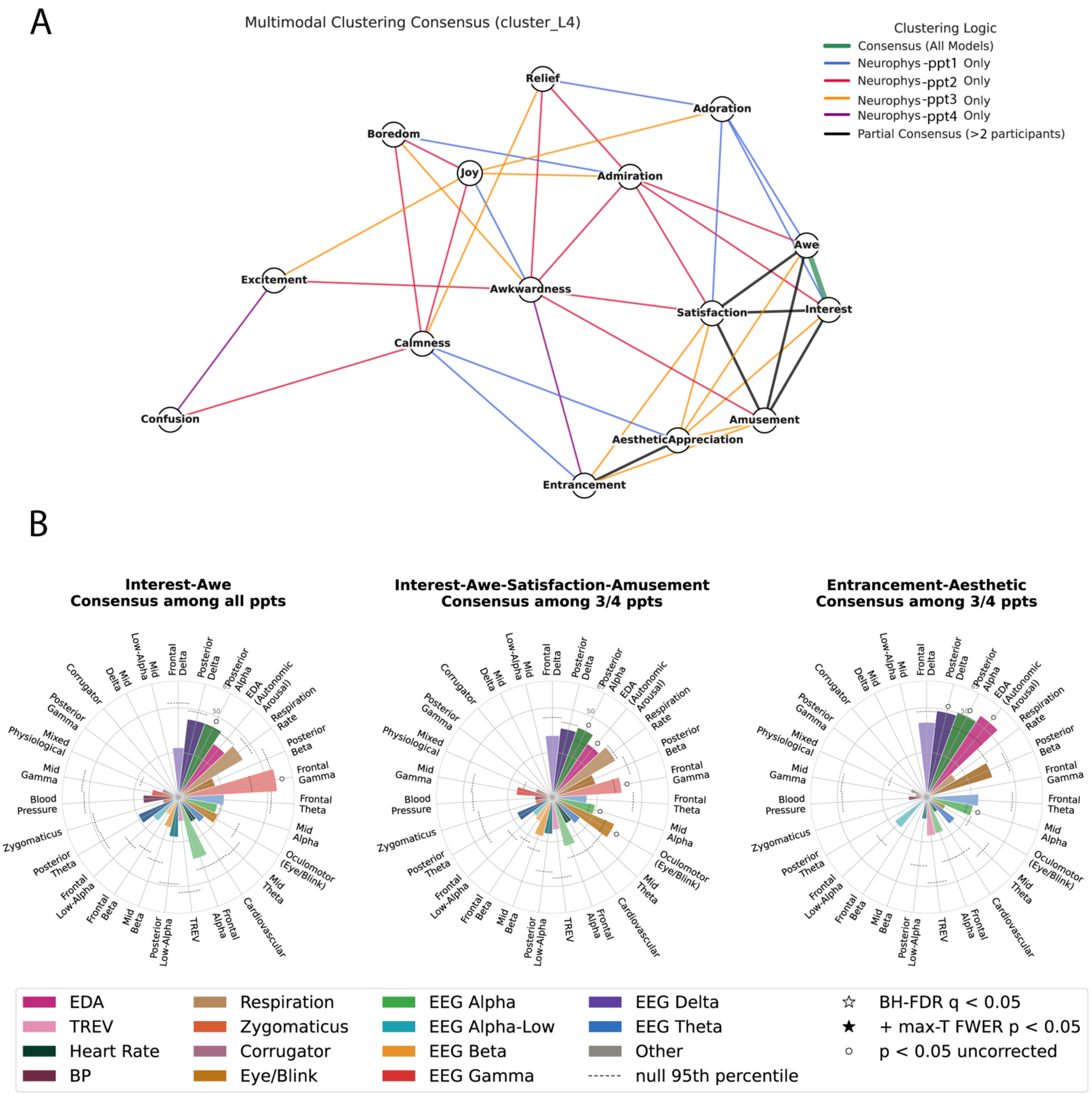


**Fig. S16.** Cross-participant emotion groupings including participant 4. A, Consensus network across all four participants, as in Fig. 6A (green, all participants; black, more than two participants; colored, a single participant). B, Cross-participant rose plots for Interest-Awe (all participants) and for Interest-Awe-Satisfaction-Amusement and Entrancement-Aesthetic Appreciation (three of four), computed and tested as in Fig. 6B; open circles, uncorrected $p < 0.05$. No cluster survived correction.

**Supplementary Table T5. Cross-participant consensus statistics for Fig. 6B.** For each consensus emotion group in Fig. 6 (columns), the feature clusters (rows) that participants consistently ranked as important. Interest-Awe-Satisfaction-Amusement and Entrancement-Aesthetic Appreciation were consensus groups in all three participants; Excitement with others (Excitement, Relief, Interest, Awe, Satisfaction, Amusement), Excitement-Admiration-Adoration and Boredom-Confusion-Joy in two of three. Rows are ordered by total consensus score across groups, so clusters near the top form the shared profile. Data are from ppt 1-3; ppt 4 was excluded because only 10 of 15 emotions were modelled. Each cell gives the cross-participant Borda score (percentage of the maximum attainable, equal to the wedge height in Fig. 6B) and, below it, the one-sided permutation p-value from 5,000 within-participant shuffles of Borda points across clusters[5]. Corrections were applied within each column, over clusters present in at least two participants: filled star (★), studentized max-T family-wise $p < 0.05$ (S4); open star (☆), Benjamini–Hochberg FDR $q < 0.05$ (S3); both shaded blue. Open circles (○, gray) mark uncorrected $p < 0.05$ (exploratory); dashes (–) mark clusters present in one participant only (not tested); empty cells, no points. This test establishes cross-participant agreement, not group specificity.

**Cross-participant agreement (rank-shuffle null)**
**★ max-T FWER p<.05 ☆ BH-FDR q<.05 ○ uncorrected p<.05 – not in test family**

| | Interest-Awe-Satisfaction-Amusement | Entrancement-Aesthetic Appreciation | Excitement-with others | Excitement-Admiration-Adoration | Boredom-Confusion-Joy |
|---|---|---|---|---|---|
| **Frontal Delta** | 42.5 p=0.211 | 51.7○ p=0.035 | 40.0 p=0.303 | 45.0 p=0.186 | **66.7★ p=0.002** |
| **Posterior Delta** | 47.9 p=0.102 | 60.0○ p=0.016 | 50.0 p=0.064 | 41.1 p=0.268 | 40.6 p=0.176 |
| **Posterior Alpha** | 45.4○ p=0.012 | **62.5★ p=0.001** | 44.4○ p=0.005 | 43.3○ p=0.039 | 32.8 p=0.088 |
| **EDA (Autonomic Arousal)** | 44.2○ p=0.011 | 50.8○ p=0.013 | 38.9○ p=0.028 | 30.6 p=0.240 | 27.8 p=0.203 |
| **Respiration Rate** | 50.4 p=0.066 | 23.3 p=0.478 | 43.9 p=0.179 | 31.7 p=0.533 | 32.8 p=0.387 |
| **Oculomotor (Eye/Blink)** | 46.7○ p=0.009 | 14.2 p=0.553 | **51.9★ p=0.000** | 40.6 p=0.055 | 10.0 p=0.922 |
| **Mid Alpha** | 24.2 – | 32.5 – | 27.2 – | 32.2 – | 32.8 – |
| **Posterior Beta** | 25.4 p=0.742 | 27.5 p=0.420 | 31.7 p=0.651 | 15.0 p=0.938 | 44.4 p=0.119 |
| **Frontal Gamma** | 37.1 p=0.087 | | 29.2 p=0.310 | 18.3 p=0.656 | **51.7☆ p=0.004** |
| **Frontal Alpha** | 35.0 p=0.130 | 16.7 p=0.351 | 32.5 p=0.173 | 18.3 p=0.668 | 28.3 p=0.388 |
| **Frontal Theta** | 23.8 p=0.787 | 15.8 p=0.617 | 26.4 p=0.822 | 42.8 p=0.245 | 20.6 p=0.753 |
| **Cardiovascular** | 20.4 p=0.635 | | 26.4 p=0.443 | 33.3 p=0.181 | 37.8 p=0.123 |
| **Mid Theta** | 23.8 p=0.465 | 12.5 p=0.732 | 28.9 p=0.261 | 27.2 p=0.359 | 15.6 p=0.641 |
| **Posterior Low-Alpha** | 25.4 p=0.748 | 15.0 p=0.664 | 26.1 p=0.840 | 17.2 p=0.905 | 17.2 p=0.838 |
| **Mid Beta** | 27.9 p=0.305 | | 30.8 p=0.194 | 33.3 p=0.179 | 6.1 p=0.938 |
| **Frontal Low-Alpha** | 17.1 p=0.728 | 27.5 p=0.311 | 15.6 p=0.859 | 8.3 p=0.923 | 27.8 p=0.198 |
| **TREV** | 22.5 p=0.812 | 15.8 p=0.617 | 28.3 p=0.772 | 27.8 p=0.658 | |
| **Frontal Beta** | 18.8 p=0.506 | | 27.5 p=0.210 | 30.6 p=0.156 | 17.2 p=0.400 |
| **Posterior Theta** | 26.7 p=0.701 | | 28.6 p=0.757 | 22.8 p=0.785 | 15.0 p=0.891 |
| **Zygomaticus** | 12.1 p=0.975 | | 19.7 p=0.957 | 32.2 p=0.518 | 20.6 p=0.756 |
| **Blood Pressure** | 12.1 p=0.976 | 12.5 p=0.796 | 16.4 p=0.982 | 10.6 p=0.965 | 24.4 p=0.653 |
| **Mid Gamma** | 19.6 p=0.886 | | 23.3 p=0.909 | 18.9 p=0.858 | 5.6 p=0.991 |
| **Mixed Physiological** | 12.1 – | 11.7 – | 15.0 – | 23.3 – | |
| **Corrugator** | | | 2.2 p=0.999 | 6.1 p=0.970 | 12.8 p=0.887 |
| **Mid Delta** | | | | 10.0 – | |
| **Mid Low-Alpha** | | | 2.8 – | 5.6 – | |
| **Posterior Gamma** | | | | | 6.1 – |

**SI References**


1. Ojala, M. & Garriga, G. C. Permutation Tests for Studying Classifier Performance. in *2009 Ninth IEEE International Conference on Data Mining* 908–913 (IEEE, Miami Beach, FL, USA, 2009). doi:10.1109/ICDM.2009.108.
2. Combrisson, E. & Jerbi, K. Exceeding chance level by chance: The caveat of theoretical chance levels in brain signal classification and statistical assessment of decoding accuracy. *J. Neurosci. Methods* **250**, 126–136 (2015).
3. Benjamini, Y. & Hochberg, Y. Controlling the False Discovery Rate: A Practical and Powerful Approach to Multiple Testing. *J. R. Stat. Soc. Ser. B Methodol.* **57**, 289–300 (1995).
4. Westfall, P. H. & Young, S. S. *Resampling-Based Multiple Testing: Examples and Methods for p-Value Adjustment*. (John Wiley & Sons, 1993).
5. Phipson, B. & Smyth, G. K. Permutation p-values should never be zero: calculating exact p-values when permutations are randomly drawn. *Stat. Appl. Genet. Mol. Biol.* **9**, (2010).

#### Legends for Extended Figure and Table

**Extended Data Figure 1.** Demonstrates Machine Learning pipeline, refer to Methods

**Extended Data Table 1.** Variability of emotion ratings across participants. For each emotion, the mean, s.d. and coefficient of variation (CV) of raw and normalized counts across participants, and the mean and s.d. of within-participant ranks (1 = most frequent). Rank s.d. ranged from 6.05 (Calmness) to 1.00 (Confusion); agreement in rank order was moderate (Kendall's W = 0.504).
